# Deep learning-based aberration compensation improves contrast and resolution in fluorescence microscopy

**DOI:** 10.1101/2023.10.15.562439

**Authors:** Min Guo, Yicong Wu, Chad M. Hobson, Yijun Su, Shuhao Qian, Eric Krueger, Ryan Christensen, Grant Kroeschell, Johnny Bui, Matthew Chaw, Lixia Zhang, Jiamin Liu, Xuekai Hou, Xiaofei Han, Zhiye Lu, Xuefei Ma, Alexander Zhovmer, Christian Combs, Mark Moyle, Eviatar Yemini, Huafeng Liu, Zhiyi Liu, Alexandre Benedetto, Patrick La Riviere, Daniel Colón-Ramos, Hari Shroff

**Affiliations:** State Key Laboratory of Modern Optical Instrumentation, College of Optical Science and Engineering, Zhejiang University, Hangzhou, China; Laboratory of High Resolution Optical Imaging, National Institute of Biomedical Imaging and Bioengineering, National Institutes of Health, Bethesda, Maryland, USA; Advanced Imaging and Microscopy Resource, National Institutes of Health, Bethesda, Maryland, USA; Janelia Research Campus, Howard Hughes Medical Institute (HHMI), Ashburn, VA, USA; Laboratory of Molecular Cardiology, National Heart, Lung, and Blood Institute, National Institutes of Health, Bethesda, Maryland, USA; Center for Biologics Evaluation and Research, U.S. Food and Drug Administration, Silver Spring, MD, USA; NHLBI Light Microscopy Facility, National Institutes of Health, Bethesda, MD, USA; Department of Biology, Brigham Young University-Idaho, Rexburg, ID, USA; Department of Neurobiology, UMass Chan Medical School, Worcester, MA; Faculty of Health and Medicine, Division of Biomedical and Life Sciences, Lancaster University, UK; Department of Radiology, University of Chicago, Chicago, IL, USA; MBL Fellows Program, Marine Biological Laboratory, Woods Hole, MA, USA; Wu Tsai Institute, Department of Neuroscience and Department of Cell Biology, Yale University School of Medicine, New Haven, CT, USA

## Abstract

Optical aberrations hinder fluorescence microscopy of thick samples, reducing image signal, contrast, and resolution. Here we introduce a deep learning-based strategy for aberration compensation, improving image quality without slowing image acquisition, applying additional dose, or introducing more optics into the imaging path. Our method (i) introduces synthetic aberrations to images acquired on the shallow side of image stacks, making them resemble those acquired deeper into the volume and (ii) trains neural networks to reverse the effect of these aberrations. We use simulations and experiments to show that applying the trained ‘de-aberration’ networks outperforms alternative methods, providing restoration on par with adaptive optics techniques; and subsequently apply the networks to diverse datasets captured with confocal, light-sheet, multi-photon, and super-resolution microscopy. In all cases, the improved quality of the restored data facilitates qualitative image inspection and improves downstream image quantitation, including orientational analysis of blood vessels in mouse tissue and improved membrane and nuclear segmentation in *C. elegans* embryos.

## Introduction

Fluorescence microscopes offer diffraction-limited imaging only when optical aberrations are absent. Such aberrations can arise due to optical path length differences introduced anywhere in the imaging path, including from instrument misalignment, optical imperfections, or differences in refractive index between the heterogenous and refractile sample, immersion media, or objective immersion oil. Sample-induced optical aberrations usually dominate and are often the reason that three-dimensional (3D) fluorescence image volumes show obvious deterioration in image signal-to-noise ratio (SNR), contrast, and resolution deeper into the image volume.

One method of compensating for these aberrations is via adaptive optics (AO^1,2^), a broad class of techniques that measure the aberrated wavefront and subsequently apply an equal and opposite ‘corrective’ wavefront, restoring diffraction-limited^3^ or even super-resolution^4^ imaging throughout the image volume. Once the aberrated wavefront is determined, an adaptive element such as a deformable mirror or spatial light modulator is used to apply the correction. Although these methods are effective, the process of determining the wavefront typically slows acquisition and/or applies more illumination dose than imaging without AO. From a practical perspective, implementing AO is nontrivial and adds considerable expense to the underlying microscope. Thus, AO remains the province of relatively few labs, and there is a need for new methods that can reverse the effects of optical aberrations without sacrificing temporal resolution, imparting more dose to the sample, or adding additional hardware to the microscope.

Deep learning approaches can computationally reverse image degradation, and have been used successfully in denoising^5,6^, deconvolution^7,8^, and super-resolution applications^9,10^. By incorporating information about the underlying object, such methods can also learn to predict the wavefront associated with aberrated images^11-13^. With sufficient training data (matched pairs of diffraction-limited and aberrated data), we reasoned that a neural network ought to be able to directly predict the diffraction-limited image from the aberrated image. The challenge then becomes accumulating appropriate training data, which would ideally be obtained without relying on AO.

Here we address this problem by (i) introducing synthetic aberrations to easily obtained near-diffraction limited data so that they resemble aberrated data and (ii) training neural networks to reverse the effect of these aberrations. We use simulations to show that application of our ‘content-aware’ approach outperforms other image restoration methods, including deconvolution with the known aberrated point spread function (PSF). We also show that our method provides performance on par with direct wavefront sensing-based AO^3^, by comparing its output to experimental ground truth. We then apply our techniques to diverse volumetric data captured with confocal, light-sheet, multi-photon, and super-resolution microscopes, finding that in all cases, resolution and contrast are substantially improved over the raw data. In addition to facilitating biological inspection, the restored data also enhanced quantitative investigation, including orientational analysis of blood vessels in mouse tissue and improved accuracy of membrane and nuclear segmentation in *C. elegans* embryos.

## Results

### Compensating for aberrations with deep learning

First, we intentionally synthetically aberrate the images acquired by fluorescence microscopes given knowledge of the physics of image formation^14,15^ (**Fig. 1, Methods, Supplementary Note 1**). Aberrations are chosen so that the aberrated images resemble those acquired deeper into the sample, where aberrations are more pronounced. The key insight of our approach is that the ‘shallow’ images on the ‘near side’ of the three-dimensional fluorescence volume are usually near-diffraction-limited and thus provide ground truth data that can be used to train a network to reverse the effect of the synthetically introduced aberrations. The trained neural network model (termed ‘DeAbe’) can then be used to reverse depth-dependent blurring on data unseen by the network, effectively mitigating the effect of aberrations without recourse to AO.

**Fig. 1.**
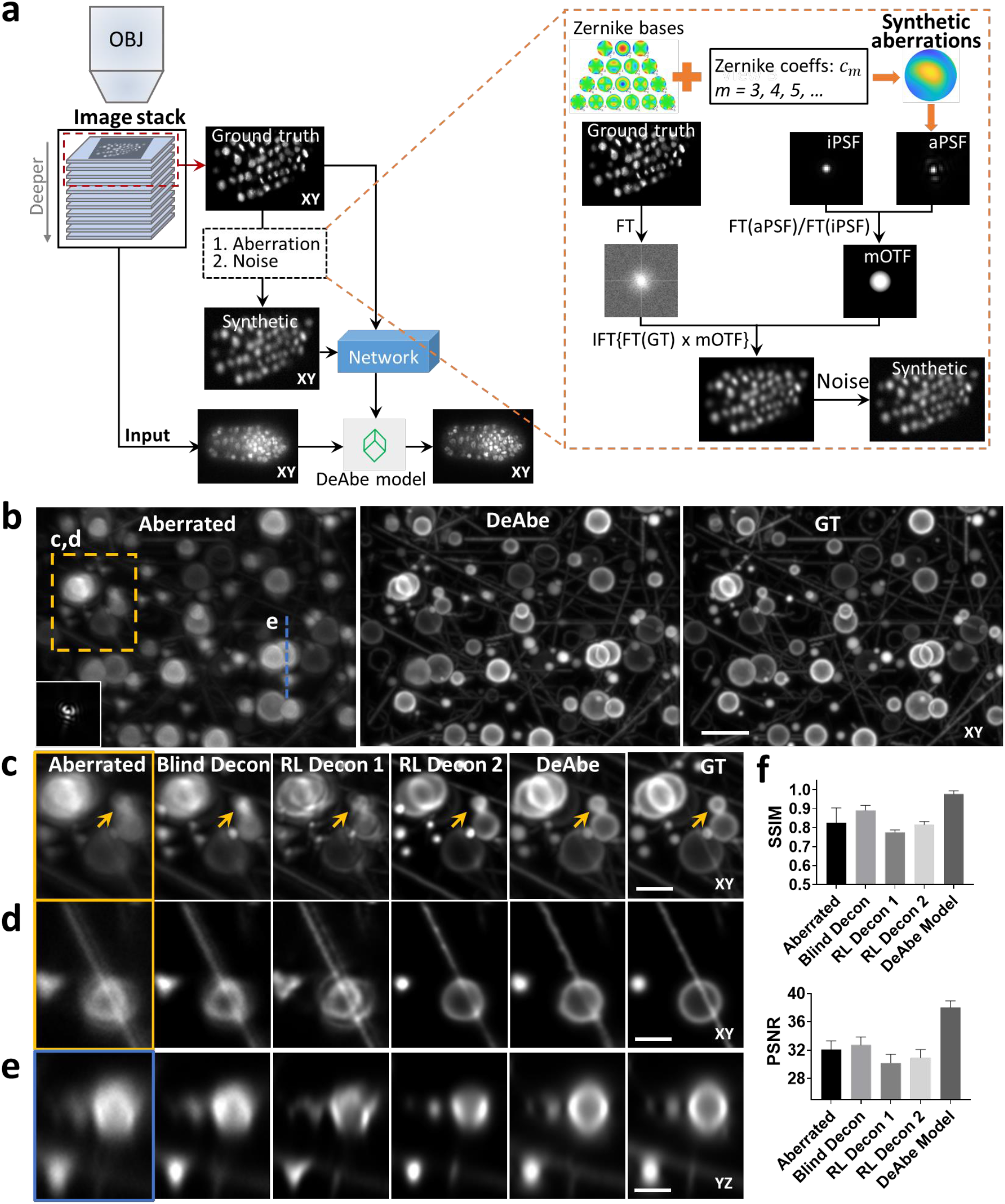
Concept and simulations illustrating deep learning-based aberration compensation. **a)** Schematic. *Left*: Fluorescence microscopy volumes are collected and near-diffraction-limited images from the shallow side of each stack are synthetically degraded to resemble aberrated images deeper into the stack. A neural network (e.g., a three-dimensional residual channel attention network, 3D RCAN) is trained to reverse this degradation given the ground truth on the shallow side of the stack, and the trained neural network (DeAbe model) subsequently applied to images throughout the stack, improving contrast and resolution. *Right*: More detailed view of synthetic degradation process. Zernike basis functions and associated coefficients (coeffs) are used to add random aberrations by modifying the ideal point spread function (iPSF) to generate an aberrated PSF (aPSF). Ground truth images (GT) are Fourier transformed (FT) and multiplied by the ratio of the Fourier transformed aberrated and ideal PSFs (essentially a modified optical transfer function, mOTF). Inverse Fourier transforming (IFT) the result and adding noise generates the synthetically aberrated images. See **Methods** for further detail on this procedure. OBJ: objective lens used to collect the stack. **b)** Simulated three-dimensional phantoms consisting of randomly oriented and positioned dots, lines, spheres, spherical shells, and circles comparing maximum intensity projections of aberrated input image (left, random aberration with root mean square (RMS) wavefront distortion of 2 radians and Poisson noise added for an SNR of ∼16, corresponding PSF in inset), network prediction (DeAbe) given aberrated input (middle), and ground truth (GT, right). Higher magnification views of dashed rectangular region are shown in **c)** (maximum intensity projection) and **d)** (single plane), additionally showing restoration given blind deconvolution (Blind Decon), Richardson-Lucy deconvolution with diffraction-limited PSF (RL Decon 1), Richardson-Lucy deconvolution with aberrated PSF (RL Decon 2). Yellow arrows indicate a reference structure for visual comparison. Twenty iterations were used for RL deconvolution and ten for blind deconvolution. **e)** As in **c, d**) but showing axial plane along dashed blue line in **b). f)** Quantitative comparisons for the restorations shown in **b-e**) using structural similarity index (SSIM, top) and peak signal-to-noise ratio (PSNR, bottom) with ground truth reference. Means and standard deviations are shown for 100 simulations (10 independent phantom volumes, each aberrated with 10 randomly chosen aberrations). Scale bars: 5 μm **b)** and 2.5 μm **c-e)**. See also **Supplementary Figs. 1-5**.

To benchmark our method, we began by simulating 3D phantoms consisting of randomly oriented and positioned dots, lines, spheres, circles, and spherical shells. We then degraded these structures by adding random aberrations and noise and evaluated the extent to which DeAbe could reverse the degradation (**Fig. 1b, Supplementary Figs. 1-7**). Visual assessments in lateral (**Fig. 1c, d, Supplementary Video 1**) and axial (**Fig. 1e, Supplementary Video 2**) views, as well as quantitative comparisons (**Fig. 1f**) demonstrated that the DeAbe model outperformed blind deconvolution^16^, Richardson-Lucy deconvolution with an ideal point spread function (PSF), Richardson-Lucy deconvolution with the aberrated PSF (known in these simulations, but unknown in general), and denoising methods (**Supplementary Figs. 6, 7**). We attribute the superior performance of DeAbe to its ability to learn a sample-specific prior, thereby better conditioning its solution relative to Richardson-Lucy deconvolution.

Importantly, simulations allowed us to further characterize DeAbe, offering insight into the regimes in which the method excels and where performance suffers. First, we found optimal performance when aberration magnitudes in the training data match the aberration magnitude in the test data (**Supplementary Fig. 1**). Over the conditions we tested, the model improved images contaminated with root mean square (RMS) wavefront distortion exceeding four radians (the highest value we tested), although performance degrades as wavefront distortion increases. Second, although we performed tests with training data containing up to the 7^th^ Zernike order, the improvement offered past order four (the value used in this work) is negligible (**Supplementary Fig. 2**). Third, DeAbe trained on a mixture of Zernike basis functions also provides notable improvement on images corrupted solely by individual Zernike functions (**Supplementary Fig. 3**), although dedicated models trained to correct specific Zernike modes are better if these modes are known in advance (**Supplementary Fig. 4**). Fourth, although DeAbe’s performance suffers in the presence of noise, it still offers noticeable visual and quantitative improvements in image quality for SNR above ∼5 (**Supplementary Fig. 5**). Finally, we explored different networks for implementing DeAbe, finding that our previous 3D RCAN^9^ offered better performance than CARE^5^, RLN^7^, or BasicVSR++^17^ architectures (**Supplementary Figs. 8, 9**).

### Comparing DeAbe predictions to experimental ground truth

We next benchmarked DeAbe against experimental datasets acquired with a lattice light sheet microscope^18^ equipped with adaptive optics for inducing and correcting aberrations (AO-LLSM^19^, **Fig. 2, Supplementary Table 1**). When imaging phalloidin-stained PtK2 cells (**Fig. 2a-f**), we induced aberrations that obscured the fine actin mesh at the cell periphery, filamentous actin, and stress fibers (**Fig. 2b-d**). Training a DeAbe model with a mixture of random aberrations restored these structures, improving contrast and resolution to a level approaching the aberration-free ground truth (**Fig. 2e, f**) or AO result (**Supplementary Fig. 10**). As for the simulations (**Supplementary Fig. 4, 6, 7**), we confirmed that training DeAbe with Zernike modes matching the underlying aberration enhanced performance compared to a random mixture of modes (**Supplementary Fig. 11**) and that DeAbe outperformed deconvolution (**Supplementary Fig. 12**) and denoising (**Supplementary Fig. 13**).

**Fig. 2.**
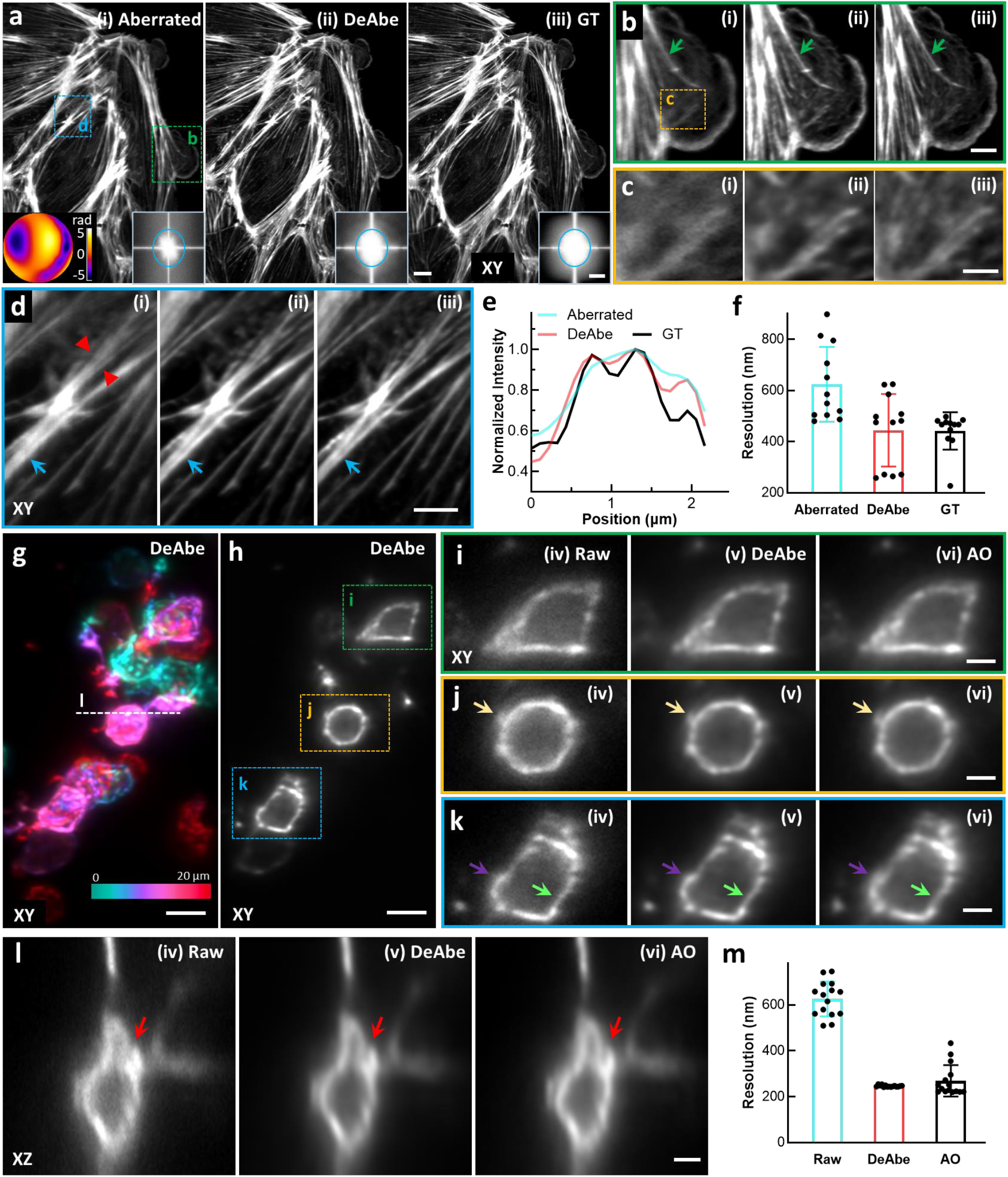
Benchmarking DeAbe against experimental ground truth and adaptive optics (AO) correction. **a)** Fixed Ptk2 cells were stained for actin using Phalloidin Alexa Fluor 488 and imaged with an AO-lattice light sheet microscope. Aberrated (i), DeAbe prediction using a model trained on random aberrations (ii), and ground truth (GT, iii) are shown. Inset in (i) shows applied aberration; right hand insets in i)-iii) show Fourier transforms, blue ellipse with 1/500 nm^-1^ horizontal extent and 1/400 nm^-1^ vertical extent. Note images have been rotated so viewing is normal to the coverslip surface, which results in anisotropic resolution in the lateral plane. **b)** Higher magnification insets of green rectangular region in **a). c)** Higher magnification views of the yellow rectangular region in **b**). **d)** Higher magnification view of blue rectangular region in **a). e)** Line profiles along red arrowheads in **d)** comparing aberrated image (blue), DeAbe prediction (red), and ground truth (GT, black). **f)** Decorrelation resolution analysis of images in **a)**. Means, standard deviations and individual data points from 12 images are shown. Green arrows in **b)** and blue arrows in **d)** highlight features improved in DeAbe or GT relative to aberrated image. XY: lateral views of sample (single planes). See also **Supplementary Figs. 10-13**. 5 dpf zebrafish embryos expressing a GFP membrane marker labeling glutamatergic neurons were fixed and imaged in an AO-lattice light sheet microscope. Image volumes were collected 40-140 μm from the surface of the fish and passed through DeAbe or corrected via AO. **g)** Depth coded lateral (XY) maximum intensity projection of volume after DeAbe compensation. Volume spans 20 μm. **h)** Single lateral plane 13 μm into imaging volume. DeAbe prediction is shown. Note images are displayed in the native view so axial direction is along optical axis of detection objective, resulting in isotropic resolution in the lateral plane. **i-k)** Higher magnification views of green, orange, and blue rectangular regions in **h)**, comparing raw (iv), DeAbe prediction (v), or AO correction (vi). **l)** Axial cross section along dashed white line in **g)**. Arrows in **i-l)** highlight membrane regions for comparisons. **m)** Lateral resolution estimates from decorrelation analysis. Means, standard deviations, and individual data points derived from 15 volumes are shown. See also **Supplementary Fig. 14**. Scale bars: 10 μm and 0.4 μm^-1^ vertical/ 0.5 μm^-1^ horizontal (insets) **a**); 5 μm **b, d, g, h**); 2 μm **c, i, j, k, l**). Data shown are representative samples from N = 12 experiments for **a-d)** and N=15 for **g-l)**.

We compared the performance of DeAbe to AO on a more challenging sample, fixed 5dpf zebrafish embryos expressing a GFP membrane marker labeling glutamatergic neurons (**Fig. 2g-m**), When acquiring image volumes 40-140 μm from the surface of the fish, AO correction and the DeAbe prediction improved lateral (**Fig. 2i-k**) and axial (**Fig. 2l**) views of the raw data, enhancing spatial resolution (**Fig. 2m**). Intriguingly, we also found examples in which the visual clarity of the DeAbe prediction appeared better than the AO correction (**Supplementary Fig. 14**), perhaps reflecting imperfect AO correction. The cell and fish samples also allowed us to investigate whether models trained on one sample type generalized to the other. As we^20^ and others^21^ have reported, we obtained superior results when training models specific to each sample type (**Supplementary Fig. 15**).

### Computational aberration compensation improves image quality on diverse volumetric data

We subsequently applied DeAbe to diverse datasets acquired with different microscope modalities, in each case training models on images derived from the shallow side of image volumes (**Fig. 3, Supplementary Fig. 16-17, Supplementary Table 1**). First, we imaged live *C. elegans* embryos expressing a pan-nuclear GFP-histone marker with inverted selective plane illumination microscopy (iSPIM)^22,23^, finding that the raw image data displayed progressive loss of contrast and resolution as a function of increasing depth, making it difficult or impossible to discern subnuclear structure (or even individual nuclei) at deeper imaging planes (**Fig. 3a, i, Supplementary Video 3**). By contrast, the DeAbe prediction restored these structures, also improving axial views (**Fig. 3a, iii**). Richardson-Lucy deconvolution also offered some improvement in image quality, albeit not to the extent of the DeAbe prediction, while also undesirably amplifying noise (**Fig. 3a, ii**). Second, we used spinning-disk confocal microscopy to image thicker adult *C. elegans* expressing the multicolor NeuroPAL transgene^24^, used for resolving neuronal identities. Depth-dependent image degradation produced raw images with dim or diffuse nuclear signal in each color channel. The DeAbe prediction improved SNR dramatically (**Supplementary Fig. 18, Supplementary Video 4**), which we suspect may prove useful in improving the accuracy of neuronal identification. Third, we applied DeAbe to images of NK-92 cells stained with Alexa Fluor 555 wheat germ agglutinin and embedded in collagen matrices, acquired with instant SIM^25^, a super-resolution imaging technique (**Fig. 3b-d, Supplementary Fig. 19, Supplementary Video 5**). Post deconvolution, the DeAbe prediction better resolved clusters of membrane-bound glycoproteins, intracellular vesicles, and membranes (‘DeAbe+’, **Fig. 3c, d**) than the raw (or deconvolved raw, **Supplementary Fig. 19**) data, especially near the limits of the 45 μm thick imaging volume. Fifth, we verified that the DeAbe prediction restored the shapes of neuronal nuclei located on the ‘far side’ of anesthetized adult *C. elegans* imaged with instant SIM, matching ground truth experiments in which we flipped the worm over (**Supplementary Fig. 20**). Sixth, we used two-photon microscopy to image live murine cardiac tissue expressing Tomm20-GFP, marking the outer mitochondrial membrane (**Fig. 3e**). Although mitochondrial boundaries were evident in the raw data 20 μm into the volume, aberrations caused a progressive loss in resolution that hindered visualization of subcellular structure at greater depths (**Fig. 3e, f**). The DeAbe prediction restored resolution throughout the 150 μm thick volume (**Fig. 3f, Supplementary Fig. 21, Supplementary Video 6**), unlike Richardson-Lucy deconvolution (**Fig. 3f**) which amplified noise without restoring the mitochondria. The DeAbe prediction similarly improved contrast and resolution when applied to volumes of fixed mouse liver stained with membrane labeled tdTomato, imaged with two-photon microscopy (**Supplementary Video 7)**. Quantitative contrast metrics (**Methods, Supplementary Fig. 22**) confirmed our visual impressions of contrast improvement provided by DeAbe.

**Fig. 3.**
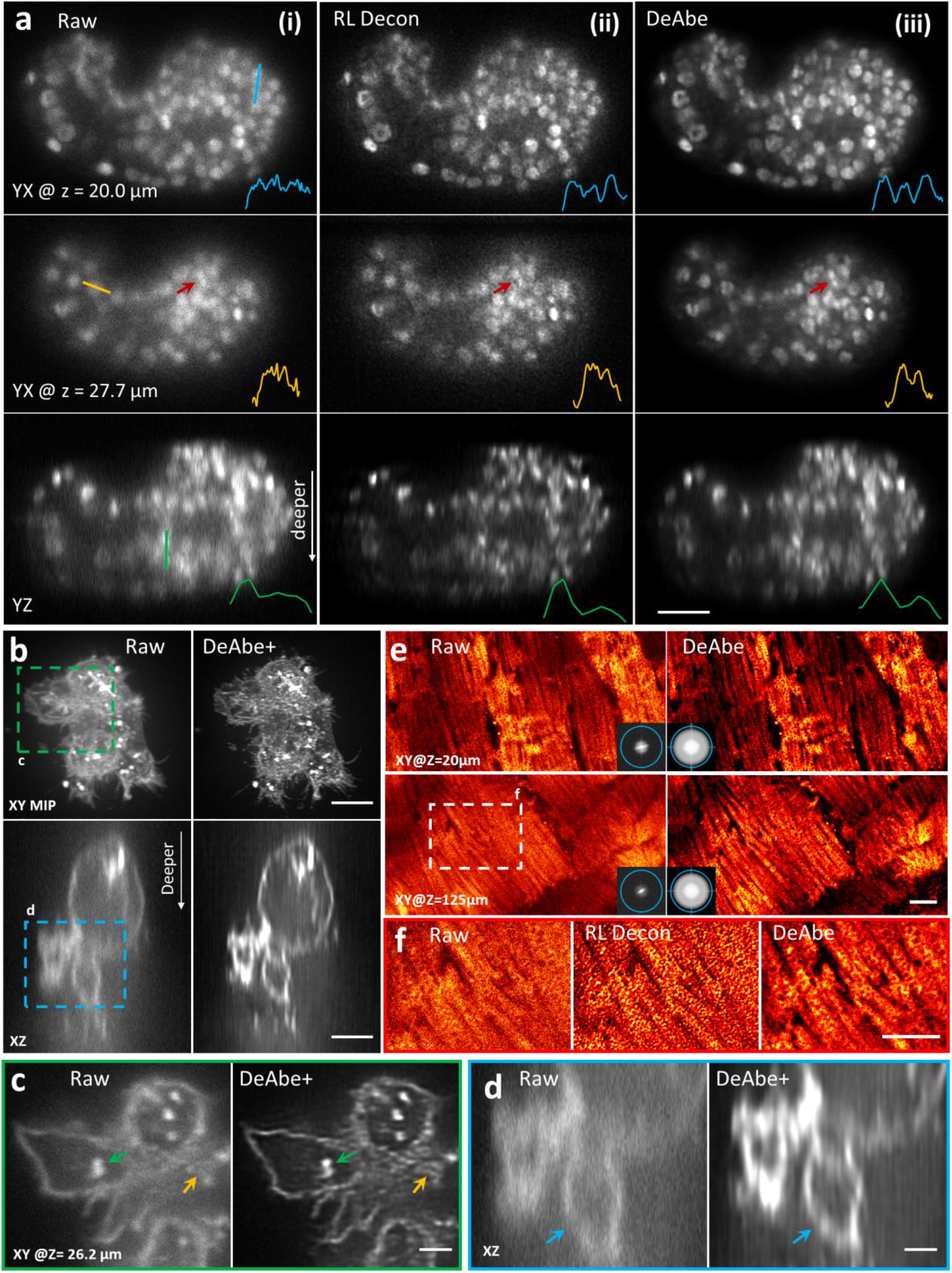
Computational aberration compensation on variety of fluorescence microscopy image volumes. **a)** Live *C. elegans* embryos expressing a pan-nuclear GFP histone marker were imaged with light sheet microscopy (**i**, left column) and the raw data processed with Richardson-Lucy deconvolution (**ii**, 10 iterations, middle column) or with a trained DeAbe model (**iii**, right column). First two rows show single planes 20.0 and 27.7 μm into the sample, third row shows axial view. Comparative line profiles through blue, yellow, and green lines are shown in insets, comparing ability to discriminate nuclei. Red arrow highlights nuclei for visual comparison. See also **Supplementary Video 3. b)** NK-92 cells stained with Alexa Fluor 555 wheat germ agglutinin and embedded in collagen matrices were fixed and imaged with instant SIM, a super-resolution imaging technique. Left: raw data, right: after application of DeAbe and deconvolution (DeAbe+, 20 iterations Richardson-Lucy). Lateral maximum intensity projections (MIP, top) or single axial planes (bottom) are shown in **b)**, and **c, d** show higher magnification views corresponding to green **c)** or blue **d)** dashed rectangular regions in **b)**. Colored arrows in **c, d** highlight fine features obscured in the raw data and better revealed in the DeAbe+ reconstructions. See also **Supplementary Video 5, Supplementary Fig. 19. e)** Live cardiac tissue containing cardiomyocytes expressing Tomm20-GFP was imaged with two photon microscopy. Raw data (left) are compared with DeAbe prediction (right) at indicated depths, with insets showing corresponding Fourier transform magnitudes. Blue circles in Fourier insets in **e)** indicate 1/300 nm^-1^ spatial frequency just beyond resolution limit. See also **Supplementary Video 6. f)** Higher magnification views of white dashed rectangular region in **e)**, emphasizing recovery of mitochondrial boundaries by DeAbe model. See also **Supplementary Fig. 21, Supplementary Video 7**. Scale bars: 10 μm **a, e)**; 5 μm **b, f)**; 2 μm **c, d)**; **e**) diameter of Fourier circle: 300 nm^-1^. Data shown are representative samples from N = 3 experiments.

Next, we applied DeAbe to samples ∼10,000-fold larger in volumetric extent (**Fig. 4a, Supplementary Video 8**). We fixed and iDISCO^26^-cleared E11.5 mouse embryos immunostained for neurons (Alexa Fluor TuJ1) and blood vessels (Alexa Fluor 594) and imaged them with low magnification confocal microscopy. Although tissue clearing nominally produces a sample with the same refractive index everywhere, we still observed pronounced depth-dependent degradation from the ‘near’ to ‘far’ side of the embryo, including in intensity (likely due to photobleaching during the acquisition) and resolution. We were able to largely reverse this deterioration by digitally compensating for photobleaching^27^ (**Methods**), applying DeAbe, and finally deconvolving the data (**Fig. 4b, Supplementary Fig. 23**). While the improvement in image quality was particularly striking in axial views (**Fig. 4b**), restorations also improved the appearance of fibrillar structures in lateral views, in both channels, throughout the volume (e.g., the vicinity of the vagus nerve and its associated nerve roots, **Fig. 4c, d**).

**Fig. 4.**
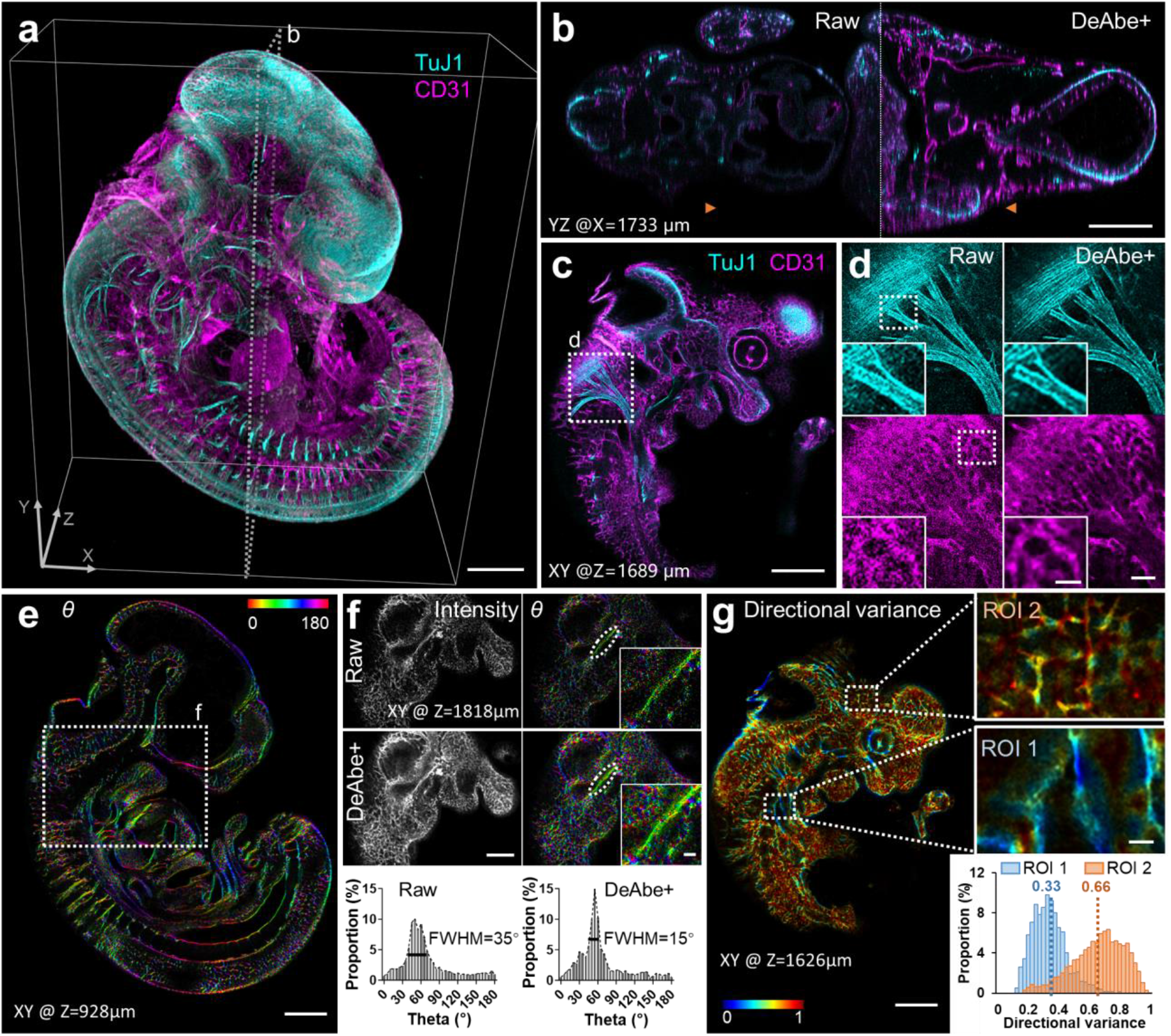
Computational aberration compensation on mm-scale cleared mouse embryo volumes. **a)** Fixed and iDISCO-cleared E11.5-day mouse embryos were immunostained for neurons (TuJ1, cyan) and blood vessels (CD31, magenta), imaged with confocal microscopy and processed with a trained DeAbe model. See also **Supplementary Video 8. b)** Axial view corresponding to dotted rectangular region in **a)**, comparing raw data and depth-compensated, de-aberrated, and deconvolved data (DeAbe+). See also **Supplementary Fig. 23. c)** Higher magnification lateral view at axial depth of 1689 μm indicated by the orange double headed arrowheads in **b). d)** Higher magnification views of white dotted region in **c)**, comparing raw (left) and DeAbe+ processing (right) for neuronal (top) and blood vessel (bottom) stains. **e)** Orientation (*θ*, transverse angle) analysis on blood vessel channel of DeAbe+ data, here shown on single lateral plane at indicated axial depth. See also **Supplementary Fig. 24, Supplementary Video 9. f)** Higher magnification lateral view of white dotted region in **e)** (note that axial plane is different), comparing intensity (left) and orientation (right) views between raw (top row) and DeAbe+ prediction (middle row). Righthand insets show higher magnification views of vessel and surrounding region highlighted by dotted lines. Bottom row indicates histogram of all orientations in the vessel highlighted with dotted ellipse, full-width-at-half maximum (FWHM) in peak region of histogram is also shown. **g)** Directional variance of blood vessel stain within the indicated plane, with higher magnification region of interest (ROI) views at right. Histogram of directional variance in both regions also shown. See also **Supplementary Fig. 25**. Scale bars: 500 μm **a, b, c, e)**; 100 μm **d)**, 50 μm inset; 300 μm **f)**, 50 μm inset; 300 μm **g)**, 50 μm inset. Data shown are representative samples from N = 3 experiments for **a-d)** and N=1 for **e-g)**.

We further investigated this qualitative impression by using automated tools^28,29^ to quantitatively assess the mean 3D orientation and directional variance (a measure of the spread in angular orientation) at each voxel in the blood vessel channel (**Fig. 4e-g, Supplementary Figs. 24, 25, Supplementary Video 9**). The DeAbe restoration resulted in cleaner separation between vessels, which aided voxel-wise quantification of these metrics even in dense regions containing many crisscrossing vessels (**Fig. 4e, Supplementary Video 9**). In deeper regions of the volume (**Fig. 4f**), the DeAbe results produced narrower angular histogram distributions of vessels than the noisy raw data (**Fig. 4f**). The improvement in quantification was also reflected in directional variance analysis. For example, when visually inspecting different regions of interest (ROI) with differential vessel alignment (**Fig. 4g**, comparing vicinity of aortic arches, (ROI 1), to diencephalon, (ROI 2)) we observed a greater difference in mean directional variance when using the DeAbe reconstruction vs. the raw data (**Supplementary Fig. 25**).

### Incorporating DeAbe in multi-step restoration further enhances resolution and contrast in 4D imaging applications

Given the performance of DeAbe thus far, we wondered if we could further boost image quality by combining DeAbe with additional networks designed to enhance spatial resolution. To test this possibility, we acquired dual-view light sheet microscopy (diSPIM^30,31^) volumetric time-lapse (‘4D’) recordings of *C. elegans* embryos expressing labels marking cell membranes and nuclei, and then passed the raw single-view data through three networks designed to sequentially compensate for aberrations (i.e., DeAbe), deconvolve the resulting predictions (‘DL Decon’), and improve resolution isotropy^5^ (‘DL Iso’, **Fig. 5a-d, Supplementary Figs. 26-29**). As expected, (**Fig. 5a**), the raw data showed increasing depth-dependent degradation in resolution and contrast, which confounded our ability to discern distinct nuclei or cell boundaries on the ‘far’ side of the volume. In comparison, the multi-step procedure offered striking improvements in resolution and contrast in both nuclear and membrane channels, largely alleviating the degradation (**Fig. 5a, b, Supplementary Figs. 27, 28, Supplementary Video 10**). Ablation experiments in which one or more of the networks were removed produced inferior results, further substantiating our hypothesis that the gains in image quality benefited from applying DeAbe (**Supplementary Fig. 30**). In the membrane channel, the multi-step restoration enabled us to automatically segment cell boundaries more accurately than in the raw data and further refine the segmentations manually up to 421 cells (**Fig. 5c, Supplementary Fig. 29, Supplementary Video 11**), exceeding previous efforts limited to the 350-cell stage^32^. Automated segmentation by successively applying DeAbe and DL Decon additionally provided a cell count closer to manual ground truth^33^ than the raw data (**Fig. 5d**) or DL Decon alone, with DL Iso providing no benefit to automated segmentation (**Supplementary Fig. 31**).

**Fig. 5.**
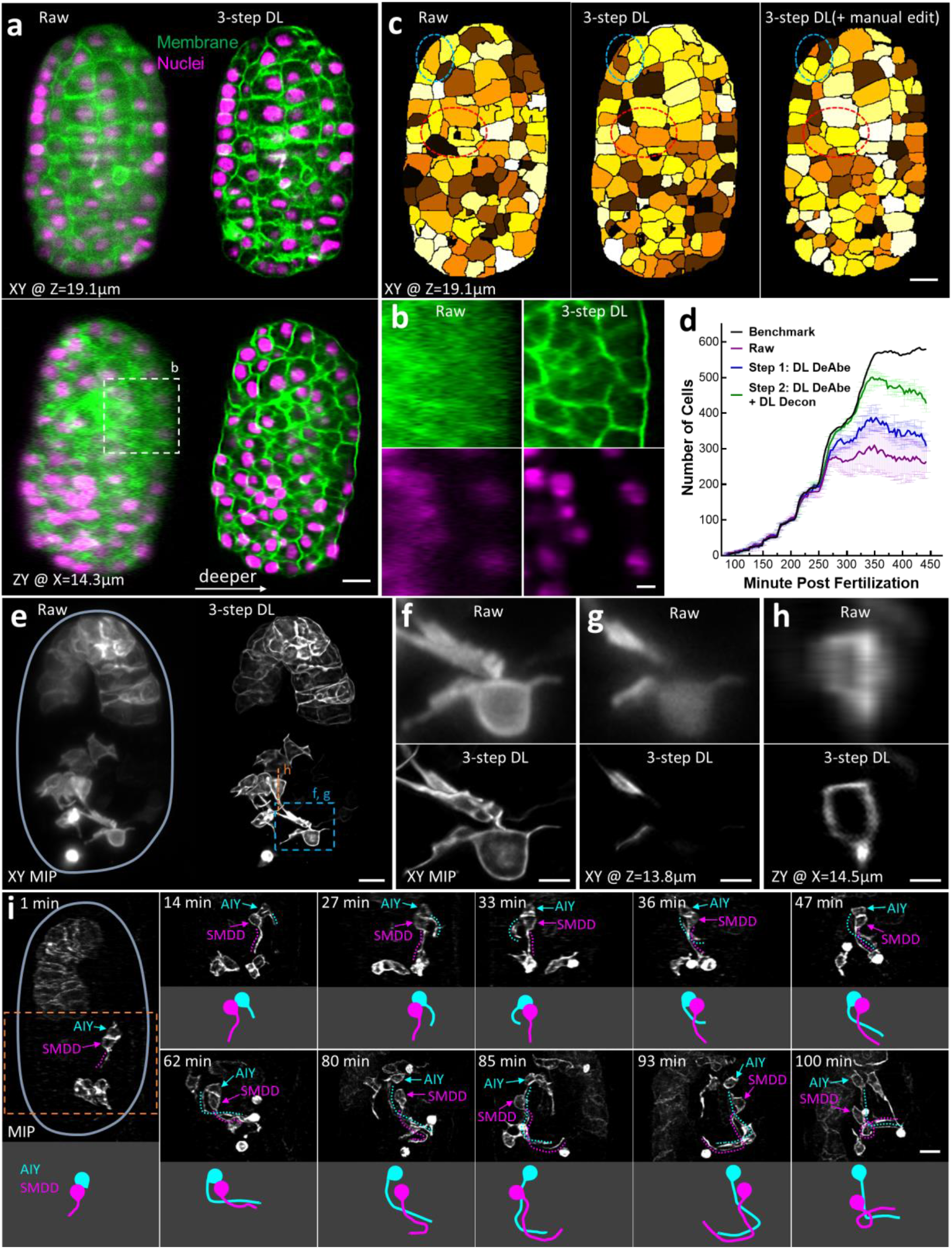
Incorporating aberration compensation into multi-step restoration dramatically improves image quality in volumetric time-lapse imaging. **a)** *C. elegans* embryos expressing GFP-labeled membrane marker (green) and mCherry-labeled nuclear marker (magenta) were imaged with dual-view light-sheet microscopy (diSPIM) and the raw data (left) from single-view recordings processed through neural networks that progressively de-aberrated, deconvolved, and isotropized spatial resolution (3-step DL, right). Single planes from lateral (top) and axial (bottom) perspectives are shown, with arrow in lower panel indicating direction of increasing depth. See also **Supplementary Video 10, Supplementary Figs. 27, 28. b)** Higher magnification axial views of membranes (top) and nuclei (bottom) deep into embryo, corresponding to dashed rectangle in **a). c)** Examples of automatic segmentation on raw (left, 319 cells), 3-step DL prediction (middle, 421 cells), and manually corrected segmentation based on DL prediction (right, 421 cells). Single planes corresponding to the upper planes in **a)** are shown. Red and blue dashed ellipses highlight regions for visual comparison. See also **Supplementary Video 11. d)** Number of cells detected by automatic segmentation of membrane marker vs. time for raw data (purple), and after successively applying the first two steps in the multistep restoration (Steps 1-2, blue and green curves), with means and standard deviations statistically derived from 3 different embryos. Ground truth from manual expert (black curve) is also shown for comparison. Inset (ellipse with dotted blue lines) highlights number count at early timepoints. See also **Supplementary Fig. 31. e)** Maximum intensity projection (MIP) images of *C. elegans* embryos expressing membrane-localized GFP under control of the *ttx3-3b* promoter, imaged with diSPIM, comparing raw single-view recordings (left) and multi-step restoration that progressively de-aberrated, deconvolved, and super-resolved the data (right, 3-step DL). Boundary of the embryo has been outlined in light blue for clarity. See also **Supplementary Figs. 33, 34, Supplementary Video 12**. Higher magnification MIP (**f**) or single lateral (**g**) or axial (**h**) plane comparisons corresponding to dashed lines or rectangle in **e**) are also shown. **i)** Time series based on 3-step DL MIP predictions highlight developmental progression of AIY (blue) and SMDD (magenta) neurites as they enter the nerve ring region. Top and bottom parts of each panel at each time point show MIP (neurites highlighted as dotted lines) vs. model of the neurites, respectively. See also **Supplementary Fig. 35**. Scale bars: 5 μm **a, c, e, f, h**); 2 μm **b, d, g**). Data shown are representative samples from N = 3 experiments.

Next, we explored replacing the final network (DL Iso) with a network designed to further enhance resolution based on ground truth acquired with expansion microscopy^9,34^ (‘DL Expan’, **Supplementary Fig. 26b**). After verifying that DL Expan improved resolution more than 2-fold on data unseen by the model (**Supplementary Fig. 32**), we applied the new multi-step restoration method to *C. elegans* embryos expressing a GFP-membrane marker labeling head neurons and gut cells (**Fig. 5e**). Compared to the raw data, the enhanced resolution offered by the deep learning prediction better resolved closely spaced membranes within and between cells (**Fig 5f-h, Supplementary Figs. 33, 34**). This capability proved especially useful when tracking the development of neurites projecting in the nerve ring, a neuropil that constitutes the brain of the animal, and which is composed of hundreds of tightly packed interwoven neurites. While the position of the neurites within the neuropil determines circuit identity and connectivity, the sequence of events leading to its innervation has not been described because of limitations in resolving these structures. We focused our analyses on the closely positioned neurons AIY and SMDD, which we identified based on morphology by comparison to labeled images in ref.^35^ and ref.^36^. SMDD is a central pioneering neuron in the nematode brain^36-38^, while its sister cell AIY^35^ is a first layer interneuron^39^ involved in thermotaxis and locomotion^40^. Observing both neurons over our 120-minute recording, we found that SMDD’s neurites grew out first, followed by AIY’s neurite. AIY’s neurite entered the nerve ring after SMDD, consistent with the SMDD’s role as a pioneer neuron (**Fig 5i, Supplementary Video 12**). Such developmental dynamics were difficult or impossible to observe in the raw data (**Supplementary Fig. 35**), or joint deconvolutions of the dual-view data due to artifacts resulting from motion between the two views (**Supplementary Fig. 36**). To illustrate that these gains in image quality can be extended to a different label imaged in a different microscope, we also restored images of nuclei labeled with a GFP histone marker and acquired with high NA diSPIM^23^, finding similarly dramatic improvements in contrast and resolution (**Supplementary Fig. 37, Supplementary Videos 13, 14)**.

In these neuronal (**Fig. 5e-i, Supplementary Videos 12**) and nuclear (**Supplementary Fig. 37, Supplementary Videos 13, 14**) recordings, although the inter-volume recording time spanned several minutes, the volume acquisition time was 1 s and 1.2 s (10 ms and 20 ms per plane, respectively), necessary to ameliorate motion blur in these rapidly repositioning^30^ embryo samples. As DeAbe is applied after data acquisition, there is no loss in temporal resolution relative to raw image capture. This capability is advantageous over AO, which always entails additional temporal cost due to the need for wavefront sensing and correction (e.g., several seconds for a single loop of correcting aberrations in the AO-LLSM experiments presented in **Fig. 2**). While this cost may be acceptable for correcting aberrations in static or slowly moving samples prior to image acquisition (by far the most common use case in AO enabled microscopy), it is too slow for the highly dynamic embryos imaged here, which would ideally benefit from rapid AO correction at each plane, at each time point.

To further underscore this point, we used the iSIM to image adult worms with a GCaMP6 marker targeted to neurons. On anesthetized (**Supplementary Fig. 38**) or partially immobilized (**Supplementary Fig. 39**) worms, DeAbe restored fine structure otherwise masked by aberrations. When performing continuous volumetric recordings at 1.5 Hz, necessary to follow calcium transients in the moving worm head and pharynx, DeAbe improved quality sufficiently that we could resolve structural details in the nerve ring that was obscured in the raw data (**Supplementary Videos 15-17, Supplementary Fig. 39**). As for the embryos, such restoration is currently infeasible with AO, due to its slow speed.

## Discussion

As we show on diverse microscopes and samples spanning multiple spatial and temporal scales, DeAbe can compensate for optical aberrations without recourse to AO: improving SNR, contrast, and resolution in fluorescence microscopy volumes without compromising the temporal resolution of data acquisition. We anticipate this capability will be useful for most labs, which lack access to sophisticated AO setups but still need to improve the quality of imaging volumes acquired using existing hardware. Besides improving the qualitative appearance of images (**Fig. 1-5**), which facilitates inspection of biological features deep within imaging volumes, DeAbe also quantitatively improves downstream image analysis. We highlight this capability by refining vessel segmentation in large, cleared tissue samples (**Fig. 4e-g**) and in enhancing the segmentation of densely packed nuclei and membranes in *C. elegans* embryos (**Fig. 5**). The latter capability may prove particularly useful in the creation or extension of 4D morphological atlases^32^, which depend on high quality image data.

Several caveats are worth noting in the context of current limitations and with an eye towards future applications. First, as for any deep learning method, DeAbe provides a *prediction* at best and cannot fully recover lost information that is not present in the raw data. Second, the performance of DeAbe depends critically on the quality of the training data, and specifically on the assumption that fluorescently labeled structures are similar throughout the image volume. While this assumption was met for the samples in this work, we encourage caution when applying DeAbe on highly heterogenous specimens (or when applying DeAbe trained on one sample type to another, **Supplementary Fig. 15**), lest hallucinations arise. Third, although here we mainly trained on semi-synthetic data (**Fig. 2-5**), it would also be worth investigating how well the training derived from fully synthetic data^7^ (**Fig. 1**) generalizes to experimental data. Such an approach might prove useful in ameliorating system aberrations introduced by microscope hardware. Fourth, we focused here on correcting depth-dependent aberrations, in which the training data was corrupted by a constant aberration in each image plane. A useful future direction would be to extend our approach to explicitly account for laterally varying aberrations, as such aberrations are problematic particularly for large specimens. Finally, although we used a mixture of random low-order aberrations to train our model, enhanced performance is likely if aberrations specific to the sample (or instrument) can be inferred and used in the training procedure (**Supplementary Fig. 4, 11, 15**).

## Author Contributions

Conceived project and directed research: H.S. Implemented DeAbe framework: M.G. Designed simulations: M.G., Y.W., H.S. Wrote software: M.G., Y.W., J.L., X.Han, S.Q., Z.L. Designed experiments: M.G., C.M.H., Y.S., R.C., X.M, A.Z., C.C., M.M., E.Y., A.B., D.C-R., H.S. Prepared samples: M.G., C.M.H., Y.S., E.K., R.C., G.K., J.B., M.C., L.Z., Z.Lu, X.M., A.Z., C.C., M.M., E.Y., A.B. Performed experiments: M.G., C.M.H., Y.S., E.K., R.C., G.K., J.B., M.C., X.Han, X.M., A.Z., C.C., M.M., E.Y., A.B. Designed and performed segmentation analysis: M.G., Y.S., J.L., X. Hou, S.Q., Z.Liu. All authors examined and analyzed data. Wrote manuscript: M.G. and H.S., with advice from all authors. Provided biological insight and advice: R.C., X.M., A.Z., C.C., M.M., E.Y., A.B., D.C-R. Supervised research: E.Y., H.L, Z.Liu, A.B., P.L-R., D.C-R., H.S.

## Acknowledgments

We thank Oliver Hobert for supporting the NeuroPAL work in his lab and allowing us to use the data generated therein for this paper; Robert Weigert for the gift of the fixed mouse liver sample; Manuel Zimmer for providing strain ZIM1997; Daniela Malide for helping to prepare and image the cleared mouse tissue datasets; Leanna Eisenman for the PtK2 cell sample preparation; Emmanuel Marquez-Legorreta for helping us to prepare the zebrafish samples; Teng-Leong Chew, the Advanced Imaging Center, and the Light Microscopy facility at HHMI Janelia Research Campus for supporting experiments with the AO-LLSM system; Steve Coleman for assisting us with the Visitech iSIM; Dan Milkie for his assistance with generating and interpreting wavefront images from the modified AO-LLSM system; Nikolaj Reiser for helpful discussions; and Courtney Johnson and Xuesong Li for their comments on the manuscript. This research was supported by the intramural research programs of the National Institute of Biomedical Imaging and Bioengineering and the National Heart, Lung, and Blood Institute within the National Institutes of Health (NIH). This work was supported by the Howard Hughes Medical Institute (HHMI). This article is subject to HHMI’s Open Access to Publications policy. HHMI laboratory heads have previously granted a non-exclusive CC BY 4.0 license to the public and a sub-licensable license to HHMI in their research articles. Pursuant to those licenses, the author-accepted manuscript of this article can be made freely available under a CC BY 4.0 license immediately upon publication. This research is funded in part by the Gordon and Betty Moore Foundation. We thank the Office of Data Science Strategy, NIH, for providing a seed grant enabling us to test and validate the initial deep learning frameworks using cloud-based computational resources. H.S., P.L.R. and D.C.-R. acknowledge the Whitman and Fellows program at MBL for providing funding and space for discussions valuable to this work. M.G. acknowledges the funding support from the Hundred Talents Program of Zhejiang University. We thank the Janelia Visiting Scientist program for supporting M.G. and A.B. Z.L. acknowledges the funding support from Natural Science Foundation of Zhejiang Province (LR20F050001). H.L. acknowledges the funding support from the National Key Research and Development Program of China (2020AAA0109502), the National Natural Science Foundation of China (U1809204) and the Talent Program of Zhejiang Province (2021R51004). E.Y. acknowledges support from the Esther A. and Joseph Klingenstein Fund, the Simons Foundation, and the Hypothesis Fund. X.M. and A.Z. were supported by Intramural FDA funding. A.B. acknowledges UKRI BBSRC (project grant BB/S017127/1).

## Methods

### Deep learning-based de-aberration model

Building a de-aberration model (DeAbe) requires appropriate training data and the use of a neural network. First, based on the physics of image formation, we derived forward imaging models that allowed us to synthetically aberrate the data produced for multiple systems, including wide field, light sheet, confocal, two photon, and super-resolution structured illumination microscopes (**Supplementary Note 1**). Second, we extracted subvolumes from the shallow side of the experimentally acquired image stacks, using these data as ground truth; alternatively, when we could obtain whole aberration-free volumes, we used them as ground truth (e.g., aberration-free images of synthetic phantoms in **Fig. 1b, Supplementary Figs. 1-9**, and stacks of cells in **Fig. 2a and Supplementary Figs. 10-13**, where the aberrations are negligible due to the thickness of the sample). Third, based on the forward imaging models, we synthetically added aberrations to the ground truth images so that they resembled aberrated data present deeper within the image stacks. Together, the paired ground truth data and associated synthetically degraded data constitute training pairs. Fourth, we used these training pairs in conjunction with our 3D RCAN network^9^ to train a DeAbe model to reverse the effect of synthetic aberration. Finally, we applied the trained network to reduce the effects of aberrations in experimentally acquired image volumes unseen by the network.

We define the ‘shallow side’ of an image stack by the planes nearest to the detection objective, which are typically contaminated with least aberration and thus offer the best image quality. We then selected subvolumes on the shallow side (‘shallow subvolumes’) by visually inspecting image quality in real and Fourier space. We also examined quantitative metrics for this choice, finding that our visual impression usually coincided with a resolution degradation of ∼20% (**Supplementary Fig. 16, 17, Supplementary Table 1**). We extracted shallow subvolumes from image stacks by manually cropping with ImageJ when image size and content differed substantially across a given specimen type, or automatically with customized ImageJ macros when considering specimens with more stereotyped image size and content (e.g., as for time-lapse image volumes). For the cleared mouse embryo images (**Fig. 4**), the shallow subvolumes were further divided into smaller subvolumes (∼80 MB/volume) due to their large volume size in raw data (**Supplementary Table 1**).

As described in **Supplementary Note 1**, we expressed the aberrated wavefront *ϕ*(*r,θ*) at the back focal plane of the objective using Zernike basis functions *ϕ*_*m*_(*r,θ*) and associated Zernike coefficients *c*_*m*_

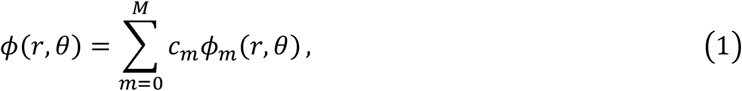

with *M* the maximum Zernike index chosen in our aberration.

We generated synthetic aberrations by using semi-randomly generated Zernike coefficients (**Fig. 1a**). We used the ANSI convention^41^ when indexing the Zernike coefficients, customizing aberrations by using different Zernike coefficients for different datasets acquired from different microscopes. For all experimental datasets, we added aberrations up to the 4^th^ Zernike order (i.e., *M* = 14), except for piston and tilt components (Z = 0, 1, 2). The amplitudes of the Zernike coefficients were randomly generated, but subject to pre-defined bounds. We initially set an upper bound of 0.5 rad for all Zernike coefficients, then added an additional 1 rad for defocus (Z = 4) and spherical (Z = 12) components to mimic the more severe contamination caused by defocus and spherical aberrations commonly encountered in experimental datasets, i.e:

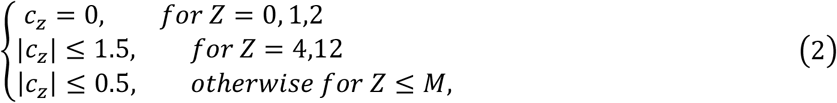

with *M* = 14 for all experimental datasets.

For each shallow side subvolume, 10 independent sets of aberrations were generated and used for synthetic degradation, thereby augmenting the data 10-fold. Processing was performed with custom MATLAB code (MathWorks, R2022b), with further details provided in the *Code availability* section.

We employed 3D RCAN (https://github.com/AiviaCommunity/3D-RCAN), appropriate for 3D image volumes, for generating the DeAbe model based on the training data pairs. We trained individual DeAbe models for each microscope and each sample type. For training, we set the number of epochs to 200; the number of steps per epoch to 400; the training patch size to 64 × 64 × 64; the number of residual blocks to 5; the number of residual groups to 5; and the number of channels to 32. When applying the model, the patch size was set to 256 × 256 × 256. Image volumes larger than this patch size were divided into patches, the network applied to each patch, and the patches stitched together via linear blinding to minimize boundary artifacts^8^ (unless specified otherwise, we used this setting for applications of 3D RCAN). Training and model application was performed within Python 3.7.0 on a Windows 10 workstation (CPU: Intel Xeon, Platinum 8369B, two processors; RAM: 256 GB; GPU: NVIDIA GeForce RTX 3090 with 24 GB memory). More details on datasets and training parameters are listed in **Supplementary Table 1**.

### Multi-step image restoration with deep learning

The multi-step image restoration pipeline combines the DeAbe model with two additional networks to progressively improve image resolution and contrast: (1) the DeAbe model to reverse degradation from aberrations (“DL DeAbe”); (2) a deconvolution network designed to mimic the image quality improvement afforded by multiview imaging (“DL Decon”, see the section *Deep learning-based deconvolution*); (3) an axial resolution enhancement network to improve resolution isotropy (“DL Iso”, see the section *Deep learning-based axial resolution enhancement*); or a network designed to predict the improved resolution provided by expanded samples (“DL Expan”, see the section *Deep learning-based expansion*).

### Deep learning-based deconvolution

As for our previous attempts at deep-learning based multiview deconvolution^8^, we used a single-view image volume as input, and attempted to restore image resolution and contrast that approximated the result from multiview joint deconvolution. The training data were acquired by dual-view light sheet microscopy^30^, either a ‘symmetric’ diSPIM equipped with 0.8/0.8 NA objectives^31^ (**Fig 5e-i, Supplementary Figs. 30, 32-36**) or a higher NA ‘asymmetric’ diSPIM equipped with 1.1 / 0.67 NA objectives^23^ (**Fig 5a-d, Supplementary Figs. 27-29, 37**). First, raw images were de-aberrated with the DeAbe model. Then de-aberrated images from the two views were jointly deconvolved to achieve reconstructions with near isotropic spatial resolution and good image quality throughout the reconstruction. With training data consisting of the single-view de-aberrated images as input and the jointly deconvolved images as ground truth, we then used another 3D RCAN for the deconvolution model (DL Decon). For all datasets, the number of epochs for training was 200; the number of steps per epoch was 400; the training patch size was 64 × 64 × 64; the number of residual blocks was 5; the number of residual groups was 5; and the number of channels was 32. Training and model application was performed within Python 3.7.0 on a Windows 10 workstation (CPU: Intel Xeon, Platinum 8168, two processors; RAM: 512 GB; GPU: Nvidia Quadro RTX6000 with 24 GB memory). We note that although training DL Decon required dual-view image volumes, applying DL Decon needs only single-view image volumes acquired from single-view light sheet microscopy (iSPIM)^22^.

### Deep learning-based axial resolution enhancement

The images predicted by the DL Decon model were not perfectly isotropic, i.e., the axial resolution (although improved over the raw input images) is worse than the lateral resolution. Thus, for some experiments we used an additional network to enhance axial resolution (DL Iso, **Fig. 5a, b, Supplementary Figs. 27-30, Supplementary Videos 10, 11**). CARE^5^ software (https://github.com/CSBDeep/CSBDeep) was employed to train the a ‘DL Iso’ model based on the predictions derived from serially applying the DeAbe and Decon models to raw input images. We used 100 3D volumes, each spanning 360 × 480 × 310 voxels, for training data. Training was performed on the xy planes (lateral views), using a 2D PSF (consisting of a point blurred with a 1D Gaussian function, sigma = 2.5 pixels along the y dimension) an axial downsampling factor of 6, and a patch size of 64 × 64 to create training pairs. The training was performed within Python 3.7.0 on a Windows 10 workstation (CPU: Intel Xeon, Platinum 8168, two processors; RAM: 512 GB; GPU: Nvidia Quadro RTX6000 with 24 GB memory).

### Deep learning-based expansion

As an alternative to DL Iso, we also trained a model to improve the resolution based on data acquired with expansion microscopy (DL Expan). First, physically expanded samples (**Supplementary Fig. 32**) were imaged on the symmetric 0.8 NA diSPIM. Second, dual-view raw images were jointly deconvolved and used as ground truth images. Third, the ground truth images were synthetically degraded to resemble low-resolution conventional images acquired on the diSPIM, following our previous procedure^9^. Last, the 3D RCAN network was employed to train the DL Expan model based on the training data (i.e., synthetically degraded and ground truth pairs).

For the worm embryo data with DAPI labeled nuclei (**Supplementary Fig. 37**), dual-view raw image volumes from 15 expanded worm embryos were acquired and jointly deconvolved to produce 15 high-resolution image volumes. These 15 volumes were then synthetically degraded to generate low-resolution images. For the worm embryo data with TTX3B neurites labeled (**Fig 5e-i, Supplementary Figs. 32-35**), dual view image volumes from 71 expanded worm embryos were acquired and manually cropped to select regions containing TTX3B neurites (this was necessary given the sparsely labeled neurites present in the raw images). Cropped images were jointly deconvolved to produce 71 high-resolution image volumes. These 71 volumes were then synthetically degraded to generate synthetic low-resolution image data. For each dataset, the low-resolution and high-resolution paired volumes were then used to train the 3D RCAN based DL Expan model. The number of epochs for training was set to 300; the number of steps per epoch to 400; the training patch size to 64 × 64 × 64; the number of residual blocks to 5; the number of residual groups to 5; and the number of channels to 32. The training was performed within Python 3.7.0 on a Windows 10 workstation (CPU: Intel Xeon, Platinum 8369B, two processors; RAM: 256 GB; GPU: NVIDIA GeForce RTX 3090 with 24 GB memory).

### Simulations on phantom objects

To evaluate the quality and performance of our DeAbe model, we generated 3D phantom objects consisting of five types of structures in MATLAB (Mathworks, R2022b, with the Image Processing Toolbox): dots, lines, circles, spheres, and spherical shells^27^. Phantoms were randomly oriented and located in a volume of 256 × 256 × 256 voxels, with voxel size 0.13 × 0.13 × 0.13 μm^3^. We simulated the blurring introduced by light sheet microscopy (**Supplementary Note 1**) by convolving the phantom with an ideal, noise-free PSF resembling that of our light sheet system (with 1.1 NA water dipping objective, detection wavelength of 0.532 μm and an illumination light sheet thickness of 2 μm). Aberrated data was generated by altering the ideal PSF according to the synthetic aberration procedure described above.

To create synthetic aberrations, we adopted Equation (1) and generated Zernike coefficients semi-randomly in MATLAB, with each Zernike coefficient *c*_*m*_ subject to a pre-defined upper bound *T*_*m*_:

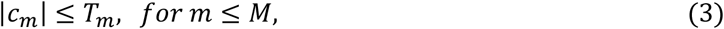

with *m* the Zernike index following the ANSI convention and *M* the maximum Zernike index.

We omitted piston and tilt components (*m* = 0, 1, 2) and weighted lower order Zernike components (Defocus *m* =4, astigmatism *m*=3,5, and spherical *m*=12) more as these aberrations are commonly observed in real samples:

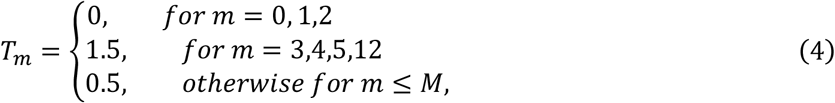

with *M* defined based on the desired Zernike order:

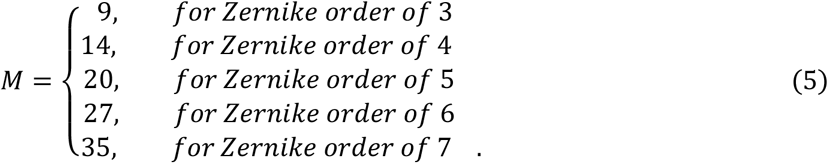

For **Supplementary Fig. 2**, we varied *M* to explore the effect of different Zernike orders on de-aberration performance by setting *M* = 9, 14, 20, 27, and 35 corresponding to Zernike orders 3-7. For all other simulations, we set *M* = 14.

The Root Mean Square (RMS) wavefront distortion of an aberration with Zernike coefficients *c*_*m*_ (*m* = 3, 4,5, …, *M*) is:

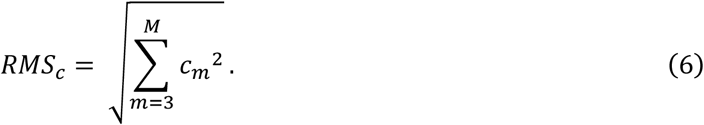

The RMS wavefront distortion for aberrations defined by upper bounds *T*_*m*_ (*m* = 3, 4,5, …, *M*) is:

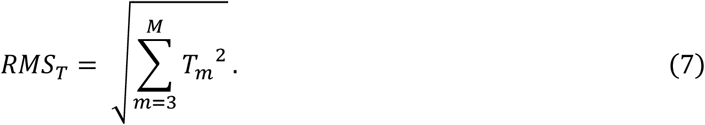

To create training data, we synthetically aberrated phantoms with two types of aberrations:

1) a random mixture of aberrations containing different Zernike components, with the amplitude of the aberrations subject to upper bounds. This type of aberrations was first generated with a set of initial Zernike coefficients *c*_*m*_ based on Equations (3-5), and then rescaled to a maximum RMS of Ω wavefront distortion (e.g., Ω = 1,2, or 4 rad) to obtain the final Zernike coefficients *c*_*m*−*final*_:

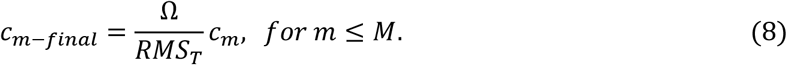 These aberrated training data were used to train the general DeAbe models (i.e., all but the model trained to counter the defocus mode specifically) used in all figures and videos showing simulated phantoms.
2) a single aberration mode of defocus with amplitude subject to upper bounds, i.e., the upper bounds of each Zernike coefficient were zeros except for the defocus mode (*m* =4):

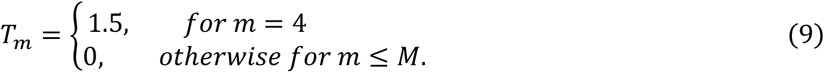

By replacing Equation (4) with Equation (9), we could generate the defocus aberration the same way as for the first aberration type (1). These training data were only used to train the specific defocus DeAbe model used in **Supplementary Fig. 4**.

For each training session, we created 50 phantoms, each consisting of different random objects. For each phantom, we generated 10 independent aberrated images with each image containing random mixtures of aberrations (**Fig 1, Supplementary Figs. 1-9, Supplementary Videos 1-2**) or only defocus aberrations (**Supplementary Fig. 4**), for a total of 500 training data pairs per session. We also added Poisson noise to the aberrated images by defining the SNR as

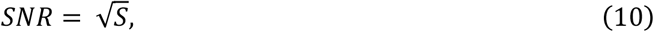

where *S* is the signal defined by the average of all pixels with intensity above a threshold (here set as 1% of the maximum intensity of the blurred objects in the noise-free image).

We employed 3D RCAN to train the DeAbe model based on simulated training data. We set the number of epochs to 200; the number of steps per epoch to 400; the training patch size to 64 × 64 × 64; the number of residual blocks to 5; the number of residual groups to 5; and the number of channels to 32. Training was performed with Python 3.7.0 on a Windows 10 workstation (CPU: Intel Xeon, Platinum 8369B, two processors; RAM: 256 GB; GPU: NVIDIA GeForce RTX 3090 with 24 GB memory).

To benchmark the performance of the DeAbe model, we created synthetic phantoms with three types of aberrations:

1) a random mixture of aberrations containing different Zernike components, with the amplitude of the aberrations subject to upper bounds. This type of aberration is the same used for training the general DeAbe models and was generated following Equations (3-5) and (8). This aberration mixture was used in **Supplementary Fig 2**.
2) a random mixture of aberrations containing different Zernike components, with the amplitude of the aberrations fixed at a certain RMS value. This aberration mixture was first generated with a set of initial Zernike coefficients *c*_*z*_ based on Equations (3-5), and then rescaled to a fixed amplitude with RMS ϒ (e.g., ϒ = 1,2, or 4 rad) wavefront distortion to obtain the final Zernike coefficients *c*_*m*−*final*_:

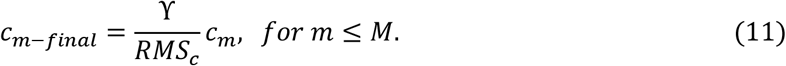 This aberration mixture was used for **Fig 1, Supplementary Figs 1**,**3**,**5, and Supplementary Videos 1-2**.
3) single aberration modes with a fixed RMS value, i.e., Zernike coefficients were set to zero except for the desired aberration mode. The single aberration modes tested in the paper include defocus (*m*=4), astigmatism (*m*=3,5), coma (*m*=7,8), trefoil (*m*=6,9), and spherical (*m*=12). If the RMS wavefront distortion is defined as ϒ (e.g., ϒ = 1,2, or 4 rad), each single aberration mode’s Zernike coefficients are:

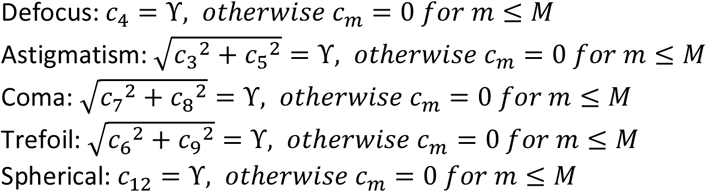

These aberrations were used to test the DeAbe performance on single aberration modes (**Supplementary Figs. 3**,**4**).

For quantitative analysis, we used structural similarity index (SSIM) and peak signal-to-noise ratio (PSNR) to evaluate the restored images provided by deep learning as well as by traditional deconvolution. The SSIM and PSNR were calculated based on image volumes with MATLAB (Mathworks, R2022b). Their mean value and standard deviation were computed from 100 simulations, each with random object structures and input aberrations.

To benchmark the performance of DeAbe using different neuronal networks, we compared our default 3D-RCAN choice with three other state-of-the-art 3D networks including CARE, RLN, and BasicVSR++. For a fair comparison, the training data pairs of phantom objects for **Fig. 1** (generated with random mixtures of aberrations) were used to train the CARE, RLN, and BasicVSR++ in addition to 3D-RCAN. Then models trained using different networks were applied to aberrated images and the prediction results compared in **Supplementary Figs. 8, 9**. 1) The CARE package was downloaded from https://github.com/CSBDeep/CSBDeep. The patch size was set to a 3D shape of 64 × 64 × 64 and the patch number was set to 32; the training epoch was 50 and the training steps per epoch was 30; and all other parameters were set to default values. 2) The RLN package was downloaded from https://github.com/MeatyPlus/Richardson-Lucy-Net. The training files and folders were reorganized to fit the input format as required by RLN. All training parameters were set to default values. 3) The BasicVSR++ package was downloaded from https://github.com/XPixelGroup/BasicSR. The batch size was set as 2 and the patch size of the 3D shape was 10 × 256 × 256; the learning rate for all modules was set to 1×10^−4^; and all other parameters were set at default values.

To distinguish de-aberration from denoising (**Supplementary Figs. 6, 7, 13**), we compared DeAbe performance with nonlocal means (NLM) and an unsupervised deep learning network, Noise2Void (N2V). The NLM denoising algorithm was implemented using the OpenCV library (https://docs.opencv.org/3.4/d5/d69/tutorial_py_non_local_means.html). We used the function fastNlMeansDenoising with the parameters h as 5, templateWindowSize as 7, and searchWindowSize as 21. The N2V package was downloaded from https://github.com/hanyoseob/pytorch-noise2void. The training files and folders were reorganized to fit the input format as required by N2V. The training epoch was 5000 and the batch size was 4; and all other parameters were set to default values.

### Preprocessing, attenuation correction, traditional deconvolution, and multiview fusion

Raw images acquired with iSIM and light sheet imaging were preprocessed by subtracting a uniform background with intensity equivalent to the average of 100 dark (no excitation light) background images. When diSPIM was operated in stage scan mode, the images were also deskewed to correct the distortion induced by stage-scan acquisition before further processing.

For the cleared mouse embryos imaged with confocal microscopy (**Fig 4, Supplementary Fig. 23, Supplementary Video 8**) and nematodes imaged with iSIM (**Supplementary Fig. 20)**, raw data was additionally preprocessed with intensity attenuation correction. The attenuation correction was performed by multiplying the raw intensity values with an exponential compensation factor:

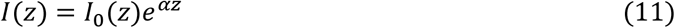

with *I*_0_(*z*) the raw intensity, *z* the depth and *α* the attenuation factor. We set *α* = 0.01 for all datasets. For the comparison of DeAbe with traditional deconvolution, we implemented both Richardson-Lucy (RL) deconvolution^42,43^ (**Fig. 1c-f, Fig. 3, Supplementary Figs. 12, 19, 21, 23**) and blind deconvolution^16^ (**Fig. 1c-f, Supplementary Fig. 12**) on the raw aberrated images. For blind deconvolution, we used the MATLAB function *deconvblind* with default settings (https://www.mathworks.com/help/images/ref/deconvblind.html). For RL deconvolution, we adopted our previously developed deconvolution package^8^ (https://github.com/eguom/regDeconProject). In one synthetic dataset (‘RL Decon 2’, **Fig. 1c-f**), we used an aberrated PSF that was generated as described in **Supplementary Note 1** and matched the aberrations in the synthetic dataset; otherwise, we used an aberration-free ideal PSF for all other datasets (**Fig. 1c-f, Fig. 3 and Supplementary Figs. 19, 21, 23**). Additionally, we also performed RL deconvolution on several datasets after DeAbe processing (**Fig. 3b-d, Supplementary Fig. 19, 23**), setting the number of iterations to 20 unless specified otherwise. All deconvolution was performed in MATLAB (MathWorks, R2022b) on a Windows 10 workstation (CPU: Intel Xeon, Platinum 8369B, two processors; RAM: 256 GB; GPU: NVIDIA GeForce RTX 3090 with 24 GB memory).

For data acquired by diSPIM, we performed multiview fusion on several datasets either for generating DL training data (**Fig. 5, Supplementary Figs. 27, 28, 33-35, 37**) or for comparisons to the DL Decon model (**Supplementary Figs. 30, 36**). The diSPIM data typically contain two view volumes, referred to as View A and View B volumes. The multiview fusion process involves registration and joint deconvolution to combine two views into a single volumetric image stack with improved resolution. The registration first rotates View B by 90 degrees along the Y-axis to align View B’s orientation with View A and then maximizes the cross-correlation function between View A and View B with affine transformations. After registration, View A and registered View B were deconvolved jointly using a modified Richardson–Lucy deconvolution algorithm as previously described^30^. Multiview fusion was achieved using custom software (https://github.com/eguom/diSPIMFusion) on a Windows 10 workstation (CPU: Intel Xeon, Platinum 8369B, two processors; RAM: 256 GB; GPU: NVIDIA GeForce RTX 3090 with 24 GB memory).

### Sample preparation and imaging

#### Live nematode embryos imaged with light sheet microscopy

Nematode strains were kept at 20°C, and grown on NGM media plates seeded with *E. coli* OP50. Strains used in this paper included BV514 (ujIS113 [*pie-1p::mCherry::H2B + unc-119(+); Pnhr-82::mCherry::histone + unc-119(+)*]), OD58 (ltIs38 [*pie1p::GFP::PH(PLC1delta1) + unc-119(+)*]), DCR6268 (olaEx3632 [*pttx-3b::SL2::PHD::GFP:: unc-54 3’ UTR + pelt-7::mCh::NLS::unc-54 3’ UTR*]), and SLS164 (ltIS138[*pie-1p::GFP::PH(PLC1delta1) + unc-119(+)*]; ujIS113 [*pie-1p::mCherry::H2B + unc-119(+); Pnhr-82::mCherry::histone + unc-119(+)*]). SLS164 was made by crossing together strains BV514 and OD58 and may have unc-119(ed3) III in the background. Strains BV514 and OD58 were gifts from Dr. Zhirong Bao.

Nematode samples were prepared for diSPIM imaging as previously described^22,31,44^: gravid adult hermaphrodites were picked into a watch glass with M9 buffer, adults were cut in half to liberate embryos, and embryos were transferred onto a poly-L-lysine coated coverslip in a diSPIM imaging chamber. For strain DCR6268 ((olaEx3632 [*pttx-3b::SL2::PHD::GFP:: unc-54 3’ UTR + pelt-7::mCh::NLS::unc-54 3’ UTR*]), labeling neuron and gut cells), embryos were imaged once they reached the bean stage of development using a fiber-coupled symmetric diSPIM (with 0.8NA/0.8NA objectives)^31^. Volumes were captured once per minute over two hours in light sheet scan mode. Each volume comprised 50 slices, with a 1 μm step size and a total acquisition time per volume of ∼1 second. For strain SLS164 (labeling cell membrane and nuclei), embryos were imaged from the 2-or 4-cell stage using a fiber-coupled asymmetric diSPIM (with 1.1NA/0.67NA objectives)^23^. Volumes were captured once every 3 minutes over 450-minute duration in stage scan mode. Each volume comprised 70 slices, with a 1.1 μm stage step size and a total acquisition time of ∼1.4 s per volume. For strain BV514 (labeling cell nuclei), embryos were imaged from the bean stage to hatching using the asymmetric diSPIM. Volumes were captured every 5 minutes in stage scan mode. Each volume comprised 60 slices, with a 1.4 μm stage step size and a total acquisition time per volume of ∼1.2 seconds. For strain OD58 (labeling cell membranes), embryos were imaged from the 4- or 8-cell stage using a symmetric diSPIM. Volumes were captured once every 3 minutes over a 450-minute period in light sheet scan mode. Each volume comprised 45 slices, with a 1 μm step size and a total acquisition time per volume of ∼0.9 seconds. For all imaging, images were acquired using 488 nm excitation (for GFP labels) or 561 nm excitation (for mCherry labels).

#### Expanded nematode embryos

*C. elegans* embryos from strain DCR6268 (labeling neurites and gut cells) were immobilized on Poly-L-Lysine (PLL) coated glass bottom dishes, bleached, digested by yatalase, fixed, and expanded. The procedure takes approximately 2 days, and is adapted from our published method^27^.

First, glass bottom dishes were coated with PLL. PLL (Sigma, Cat# P5899) powder was reconstituted in distilled water to 1mg/mL, aliquoted, and stored at −20°C. Prior to experiments, 30-50 μL of PLL was placed on the glass bottom dish (MatTek, Cat# P35G-1.5-14-C) and air dried at room temperature (RT). Coated coverslips were usually prepared up to 1 day before pre-treatment of *C. elegans* for expansion microscopy.

Second, embryos were digested, fixed, and stained with DAPI. Gravid adult *C. elegans* worms were deposited in a petri dish in PBS buffer and cut with a surgical blade to release eggs. Eggs were immobilized on a PLL coated glass bottom dish in PBS and could be processed immediately or stored at 25°C in M9 buffer until the embryos developed to the desired stage. Embryos were treated with a bleaching mixture containing 1% sodium hypochlorite (Sigma, Cat# 425044) in 0.1M NaOH/water for 2-3 minutes, rinsed 3 times in PBS, digested in 50 mg/mL Yatalase in PBS (Takara Bio, Cat# T017) for 40 minutes at RT and rinsed 3 times with PBS. It was important to treat eggs with bleach only after immobilization on the PLL surface, otherwise embryos tended to detach from the glass at later steps. Digested embryos were fixed in 4% paraformaldehyde/PBS (Electron Microscopy Sciences, Cat# RT15710) for 1 hour, then rinsed 3 times with PBS to remove fixative. Fixed embryos were permeabilized in 0.1% Triton X-100/PBS (Sigma, Cat# 93443) for 1 hour at RT with 1 μL/mL of DAPI (Thermo Fisher Scientific, Cat# D1306).

Optionally, GFP signal can be boosted by immunolabeling. Yatalase digested embryos were permeabilized with staining buffer (0.1% Triton X-100/PBS) for 1 hour before immunolabeling. Embryos were stained by an anti-GFP primary antibody (Abcam, Cat# ab290) in the staining buffer at 4°C overnight at 1 μg/mL. After primary antibody labeling, embryos were washed 3 times (30 min intervals between washes) in the staining buffer and labeled using donkey-anti-rabbit-biotin secondary antibody (Jackson ImmunoResearch, Cat# 711-067-003) in the staining buffer at 4°C overnight at 1 μg/mL. After secondary antibody labeling, the embryos were washed 3 times in the staining buffer (30 mins intervals between washes) and labeled with Alexa Fluor 488 Streptavidin in the staining buffer at 4°C overnight at 2 μg/mL (Jackson ImmunoResearch, Cat# 016-540-084). Labeled embryos were washed 3 times in the staining buffer (30 minutes between washes) before being processed for expansion microscopy. Immunolabeling was only performed on the data shown in **Supplementary Fig 32a**.

Finally, embryos were expanded. Embryos were treated with 1 mM MA-NHS (Sigma, Cat# 730300) in PBS for 1 hour at RT. Samples were rinsed 3 times in PBS, and treated with monomer solution, which was made up of acrylamide (Sigma, Cat# A9099), sodium acrylate (Santa Cruz Biotechnology, Cat# 7446-81-3), N, N’-methylenebis(acrylamide) (Sigma, Cat# 146072) and 4-Hydroxy-TEMPO (Sigma, Cat# 176141), diluted with PBS, with a final concentration of 10%, 19%, 0.1%, and 0.01%, respectively. After the treatment for 1 hour at RT, the monomer solution was replaced by gelation solution. The gelation solution shared the same reagents and concentrations as monomer solution, with the addition of tetramethylethylenediamine (TEMED, Thermo Fisher Scientific, Cat# 17919, reaching a final concentration of 0.2%) and ammonium persulfate (APS, Thermo Fisher Scientific, Cat# 17874, reaching a final concentration of 0.2%). APS was added at last, and the fresh gelation solution was immediately applied to the embryos sandwiched between the glass bottom dish and another coverslip surface for 2 hours at RT. It was important to control the gelation speed with 4-hydroxy-TEMPO as premature gelation can distort embryos and result in poor expansion quality. The polymerized embryo-hydrogel hybrid was cut out by a razor blade and digested with 0.2 mg/mL Proteinase K (Thermo Fisher Scientific, Cat# AM2548) in digestion buffer (0.5 M sodium chloride (Quality Biological, Cat # 351-036-101); 0.8 M guanidine hydrochloride (Sigma, Cat# G9284); and 0.5% Triton X-100) at 45°C overnight. Digested embryos were expanded ∼3.3-3.7 fold in distilled water, exchanging the water every 30 min until expansion was complete. Expanded samples were flipped over so that embryos were ‘on top’ (suitable for diSPIM imaging), mounted on PLL coated #1.5 coverslips (VWR, Cat# 48393-241) and secured in an imaging chamber filled with distilled water. Finally, samples were imaged using the symmetric 0.8/0.8 NA diSPIM in stage scan mode. Depending on the orientation of embryos, ∼200-300 planes were acquired for each embryo, with 1.414 μm stage step size and 20 ms per-plane exposure time.

#### PtK2 cells imaged with adaptive optical lattice light-sheet microscopy (AO-LLSM)

PtK2 cell samples were prepared by placing one 25 mm round coverslip (Warner Instruments, CS-25R17) into a 35 mm culture dish (Corning, 430165) and seeding cells at 100k cells per dish the day before fixation. Cells were washed quickly 3 times with pre-warmed PBS before fixing in 4% formaldehyde for 5 minutes at room temperature. 3 additional PBS washes were performed, and cells were permeabilized in 0.1% IGEPAL (Sigma-Aldrich, I8896) for 5 minutes at room temperature. Cells were washed with PBS 3 times, after which 250 µl of a primary antibody solution of 0.1% iGf-free BSA (Jackson ImmunoResearch, 001-000-162) and 1:400 Phalloidin Alexa Fluor 488 (ThermoFisher Scientific, A12379) in PBS was added to each coverslip. Cells were incubated at 37C for 1 hour, and a final wash of PBS with 0.05% Tween-20 (Sigma-Aldrich, P1379) and 2 additional PBS washes were performed.

Cells were imaged in PBS on a modified adaptive optical lattice light-sheet microscope^18,19^. First, a system correction was performed as previously described^19^. Lattice light sheet excitation was performed using a 488 nm laser line, a Thorlabs TL20®-MPS 0.6 NA objective lens, and a square lattice pattern (Outer NA: 0.4, Inner NA: 0.3, Cropping: 10, Envelope: 5). Image stacks (256×1500 pixel field of view (FOV) with 401 z steps) were acquired by scanning the sample stage horizontally at an angle of 32.45° relative to the optical axis of the detection objective (Zeiss Plan-Apo 20x, NA 1.0 DIC M27 75 mm) with a step size of 0.4 µm and an exposure time of 20 ms. Emission light was filtered through a Semrock BrightLine 523/40-25 emission filter and reflected onto a Hamamatsu Orca Flash 4.0 sCMOS camera via a Semrock Di03-R561-t3-32×40 dichroic. After data collection, images were deskewed using a custom analysis pipeline (https://github.com/aicjanelia/LLSM). The final voxel size after deskewing was 0.108 × 0.108 × 0.215 nm.

For training data, 40 random FOVs were selected and imaged as described above. For aberration experiments (**Fig. 2a-f, Supplementary Figs. 10-12**), a random FOV was selected and a ground truth data set was acquired. Next, an aberration was applied to the deformable mirror (DM; ALPAO DM69). These aberrations were either random, wherein each actuator on the mirror was pushed or pulled by a random amount with a fixed maximum amplitude, or a predefined Zernike mode (astigmatism, coma, or spherical). For each type of aberration, 3 different magnitudes were used, and for each magnitude 3 different FOVs were selected, yielding a total of 36 experiments. After the aberration was applied to the DM, a stack was collected. The microscope configuration was then changed to the adaptive optics (AO) configuration.

The methods for AO correction have been described previously^19^. A focused two photon (Coherent 1335240 Chameleon) spot was directed through the detection objective and scanned through the same FOV to be imaged. The collected emission was passed through a microlens array and imaged to the same camera used for image collection to function as a Shack-Hartmann (SH) wavefront sensor. The distance each spot in the SH image moves is calculated relative to a reference image, after which the DM is updated to correct the measured aberration. This process is repeated 2 additional times as the AO correction will iteratively improve until it converges. The microscope is then switched back to LLSM mode, and a final stack is acquired.

For comparative denoising experiments (**Supplementary Fig. 13**), a random FOV was selected and a ground truth stack was acquired. Then, aberrations (random, astigmatism, and coma) were applied to the DM at a single magnitude; 3 separate FOVs were examined per aberration. Once the aberration was applied to the DM, stacks were acquired with the original laser power (high SNR, **Supplementary Fig. 13c**) as well as 1/5 laser power (low SNR, **Supplementary Fig. 13b**).

#### Zebrafish embryos imaged with adaptive optical lattice light-sheet microscopy

Transgenic Zebrafish Tg(vGlut2a:Gal4); (UAS:CoChR-eGFP), featuring eGFP localized in the membrane of glutamatergic neurons, were fixed overnight at 5 dpf in 4% PFA at 4C and subsequently washed with and stored in PBS. A total of n=6 fish were used for experiments. To mount the fish onto 25 mm round coverslips, the coverslips were first treated with Poly-l-lysine, after which a thin layer of 1.5% agarose (ThermoFisher Scientific, 16520050) was cured onto the coverslip. A small channel was carved into the center of the agarose, and the fish was placed ventral side down into the channel. Finally, a small drop of 1.5% agarose was placed on top of the fish.

Fish were imaged in milliQ water on the modified AO-LLSM described above. In this case, a square lattice pattern (Outer NA: 0.4, Inner NA: 0.34, Cropping: 10, Envelope: 10) was used for excitation. Image stacks (256×512 pixel FOV with 101 z steps) were acquired by scanning the sample stages horizontally and vertically simultaneously such that the sample moved directly along the optical axis of the detection objective with a step size of 0.2 µm and an exposure time of 100 ms. Emission light was captured as described above. In this instance, deskewing of the data was not necessary and the final voxel size was 108 × 108 × 200 nm.

For training data, 42 FOVs were selected near the surface (∼0-20 µm) of the fish and imaged as described above. Next, 15 FOVs deeper within the fish (∼40 – 120 µm) were imaged first without AO correction, and next with an identical AO correction procedure as described above (**Fig. 2g-m, Supplementary Fig. 14**).

#### Live nematode adults imaged with spinning disk confocal microscopy

*C. elegans* strain OH15500 (*otIs669[NeuroPAL]; otIs672[panneuronal::GCaMP6s]*) were raised at 20°C and grown on NGM media plates seeded with OP50 *E. coli*. Young adult worms (with 2 or less visible eggs in their uterus) were picked and immobilized inside a microfluidic chip as previously described^24^. Worms were imaged by a spinning disk confocal microscope (Nikon, Ti-e) equipped with a 60×/1.2 NA water objective (Nikon, CFI Plan Apochromat VC 60XC WI), a confocal scan unit (Yokagawa, CSU-X1) and an electron multiplying CCD (EM-CCD, Andor, iXon Ultra 897). Four excitation lasers (405 nm, 488 nm, 561 nm, and 640 nm) were used for illumination, in conjunction with emission filters spanning 420-470 nm, 500-545 nm, 570-650 nm, and 660-800 nm bandwidths, respectively. The pixel size was 0.27 μm in the XY dimension and each Z-stack volume comprised 21 slices for each color, with1.5?μm step size. Each multicolor Z-stack volume was captured at a rate of just over 1 minute.

#### Fixed WGA-labeled NK-92 samples imaged with instant structured illumination microscopy

NK-92 cells (ATCC®, CRL-2407™) were rinsed with 1× PBS, and fixed with 1 ml of 4% paraformaldehyde in 1× PBS for 30 min at room temperature, rinsed in 1 ml of 1x PBS, and permeabilized in 0.1% Triton X-100 in 1× PBS for 15 min. Next, samples were rinsed with 1× PBS, and blocked with buffer containing 1% BSA (Fisher, Cat# BP9700100) in 1× PBS for 1 hour. Blocking buffer was removed, and the samples were stained with 500 μl of 1x PBS with a 1:100 dilution of Alexa Fluor 555 labelled WGA (Invitrogen, Cat# W32464), 10 U/mL phalloidin-ATTO 647N conjugate (Millipore-Sigma, Cat #65906), and 1:1000 dilution of Hoechst solution (Tocris, Cat#5117) for 1 h. Cells were washed in 1× PBS three times. We mounted samples using 90% Glycerol (Sigma, Cat# G5516) in 1x PBS.

In preparation for imaging, cells were cultured in collagen-I gels in the ImmunoCult-XF T Cell Expansion Medium (STEMCELL Technologies, Cat# 10981) with the addition of Human Recombinant Interleukin 2 (STEMCELL Technologies, Cat# 78036.3). To prepare 3 mg/ml collagen-I gel, we assembled a gel premix on ice in a prechilled Eppendorf tube. Briefly, to 1 volume of CellAdhere™ type I bovine (STEMCELL Technologies, Cat# 07001) we added 8/10 volume of DMEM, 1/10 volume of 10x PBS, 1/20 volume of 1M HEPES, and 1/20 volume of 1M (in DMSO) Alexa Fluor 488 ester (Molecular Probes, Cat# A20000). A drop of premixed gel (∼50 µL) was spread immediately on a glass surface of a plasma-treated glass-bottom 35 mm Petri dish (MatTek Corp., Cat# P35G-1.5-14-C) with a pipette tip. During polymerization (room temperature, for overnight), gels were covered with 1 mL of mineral oil (Sigma-Aldrich, Cat# M8410) to prevent evaporation of water. Before adding NK-92 cells, polymerized gels were rinsed with PBS to remove the unpolymerized gel components.

Instant structured illumination microscopy (iSIM) was performed using the commercial instant structured illumination microscope system (VisiTech Intl, Sunderland, UK) equipped with an Olympus UPlanSAapo 60×/1.3NA Sil objective, two Flash-4 scientific CMOS cameras (Hamamatsu, Corp., Tokyo, Japan), an iSIM scan head (VisiTech Intl, Sunderland, UK), and a Nano-Drive piezo Z stage (Mad City Laboratories, Madison, WI). The iSIM scan head included the VT-Ingwaz optical destriping unit. The exposure time was set to 250 ms per image frame. The voxel size was 64 × 64 × 250 nm, in x, y, and z, respectively.

#### Nerve ring calcium imaging of trapped C. elegans with instant structured illumination microscopy

Strain ABA0001 ((*lite-1(xu7); goeIs247 [ceh-24p::GCaMP6s::mKate2::unc-54 3’UTR + unc-119(+)]*) was generated by crossing TQ1101^45^ and HBR1077^46^. Adult day 1 (24 hours after late L4 stage) ABA0001 worms were raised at 20 C on standard 6 cm-diameter NGM plates seeded with *E. coli* OP50 bacteria^47^. Individual worms were picked for imaging using BIO-133 (MY Polymers) as sticky glue (in lieu of bacterial paste) into another drop of BIO-133^48^ set on a high-precision 50 × 24 mm^2^ #1.5 glass coverslip (Thorlabs, CG15KH1) between two 18 × 18 mm^2^ #1 glass coverslips (Brand, 470045) used as spacers. Another high precision 50×24 mm^2^ #1.5 glass coverslip was carefully laid on top and gently pressed downwards. The assembly was cooled on ice to ensure minimal worm movement, then flood-exposed on an aluminum sheet to 365 nm light dispensed by a LED array for 1-2 min until BIO-133 had cured^48^. The “coverslip-sandwiched” worms were then imaged with a qCMOS Orca Quest (Hamamatsu, C15550-22UP) through a 40x/1.15NA water objective (Olympus, UAPON-340) on a VisiTech iSIM imaging platform driven by Micro-Manager 2.0^49^, equipped with a 300 μm-range Z-piezo (ASI, PZ-2300FT) and 405 nm, 488 nm, and 561 nm lasers.

Image volumes of *Pceh-24::GCaMP6s* expression in the worm head were then acquired using the single-channel fast-sequence mode, with 1.2 μm axial spacing, yielding a volume acquisition rate of ∼1.5 Hz (voxel dimensions: 0.115 × 0.115 × 1.2 μm^3^). The exposure time was 14 ms. GCaMP6s fluorescence was filtered through a ET525/50m emission filter (Chroma).

#### Imaging anesthetized adult C. elegans with instant structured illumination microscopy

Adult day 1 ZIM1997 (*mzmIs52; lite-1(ce314);otIs670*)^50^ or ABA001 worms were raised at 20 C on standard 6 cm-diameter NGM plates seeded with *E. coli* OP50 bacteria^47^ and subsequently exposed to unseeded NMG plates containing 0.02% levamisole prepared in M9 buffer for 10 min. Worms were next mounted in BIO-133 as previously described, and imaged with the aforementioned VisiTech iSIM imaging platform. 3D volumes were acquired with 300 nm Z-steps at full XY-resolution (voxel dimensions: 0.115 × 0.115 × 0.300 μm^3^) sequentially (XY-Z-C) for each channel (starting with the longest excitation wavelength).

For ZIM1997, imaging was performed twice per worm (before and after flipping) so that both sides of the worm were imaged with the more favorable ‘near-side’ configuration (**Supplementary Fig. 20**). The imaging parameters for each label were as follows: 1) mTagBFP2 with 405 nm excitation, 40 ms exposure time, and an ET460/50m emission filter; 2) GCaMP6f with 488 nm excitation, 20 ms exposure time, and an emission filter of ET525/50m; 3) CYO1FP with 488 nm excitation, 30 ms exposure time, and an emission filter of ET600/50m; 4) TagRFP-T with 561 nm excitation, 40 ms exposure time, and an emission filter of ET600/50m; 5) mNeptune2.5 with 561 nm excitation, 60 ms exposure time, and an emission filter of ET690/50m. All emission filters were purchased from Chroma.

For ABA001, imaging parameters were: 1) GCaMP6s with 488 nm excitation, 30 ms exposure time, and an emission filter of ET525/50m; 2) mKate2 with 561 nm excitation, 30 ms exposure time, and an emission filter of ET600/50m. All emission filters were purchased from Chroma.

#### Two-photon microscopy on live and fixed mouse tissue

Fixed mouse liver samples and fresh ex-vivo mouse heart muscle strips were imaged with two-photon microscopy using a Leica SP8 two photon DIVE upright microscope (Mannheim, Germany), a pulsed dual beam Insight X3 Ti-Sapphire laser (MKS Spectra-Physics, Milpitas CA), a Leica 25x 1.0 NA (HC PL IRAPO) water dipping lens, and emission bandwidth tunable Leica HyD detectors in the non-descanned emission pathway. Liver samples were prepared from freshly excised liver from a 10 week-old mouse expressing a membrane-targeted peptide fused with tdTomato^51^. After excision, the mouse liver was washed in cold saline three times, fixed with 4% formaldehyde in PBS for 2 hours, and stored in PBS. Tissue harvesting procedures were approved by the NCI (for mouse liver) and NHLBI (for mouse heart) Animal Care User Committees (ACUC) respectively. Freshly excised heart muscle strips from transgenic mice expressing mitochondrial TOMM20-mNeonGreen were prepared for imaging as described^52^. tdTomato and mNeonGreen were excited using 1045 nm and 960 nm excitation with emission bandwidths of 550-700 nm and 500-600 nm, respectively. Laser excitation (ramped as a function of depth in some experiments and optimized by adjusting the objective motorized correction collar) were in the range of 1% for tdTomato and less than 20% for mNeonGreen. HyD detector gains were kept at 100% for tdTomato and 150% for mNeonGreen. Tiled images volumes of liver membrane expressing tdTomato were collected with voxels sizes set to 400 nm in the XY dimension and 500 nm in the z dimension. Z-stack volumes of mNeonGreen expressing heart strip were collected with voxels sizes set to 120 nm in the XY dimension and 500 nm in the z dimension. All imaging was conducted at an imaging speed of 600 Hz with a pinhole size of 1 A.U.

#### Cleared mouse embryos imaged with confocal microscopy

E11.5-day mouse embryos were collected in phosphate-buffered saline (PBS) and directly immersed in 4% paraformaldehyde (PFA) in PBS (pH 7.4) at 4°C overnight. Following fixation, the samples were washed with PBS and stored in PBS at 4°C for further analyses. Wholemount immunofluorescence staining was performed at 4°C. The mouse embryos were permeabilized with 0.2% Triton/PBS overnight and blocked with 10% normal goat serum and 1% BSA in 0.2% Triton/PBS overnight. The embryos were then stained with monoclonal antibody against PECAM1 (CD31, clone MEC 13.3, Cat# 553700, BD Pharmingen, 1:200 dilution) and monoclonal anti-β-tubulin III (TuJ1)) antibody (clone 2G10, Cat# T8578, Sigma-Aldrich, 1:500 dilution) in blocking buffer overnight. After washing with 0.2% Triton/PBS, the embryos were stained with secondary antibodies with Alexa 488 goat anti-rat IgG and Alexa 594 goat-anti-mouse IgG (1:250, Invitrogen, Carlsbad, CA) in blocking buffer overnight. The embryos were cleared with iDISCO^26^ and imaged using a Zeiss LSM 880 Confocal microscope with a 10X, 0.5NA air objective. To compensate for focal shift effects due to the refractive index difference between air and iDISCO we scaled the axial voxel size of images by 1.56 before processing for DeAbe.

### Quantitative image quality analysis

#### Decorrelation resolution metric

Decorrelation analysis^53^ was used to estimate image resolution (**Fig. 2f, m, Supplementary Fig. 13f, Supplementary Fig. 16**). Code was downloaded from https://github.com/Ades91/ImDecorr, and the MATLAB version of the code was used. For statistical analysis, the resolution of each image was estimated first, then means and standard deviations were calculated from N=12 (**Fig. 2f**) or N= 15 (**Fig. 2m**) images.

#### Normalized Discrete Cosine Transform Shannon Entropy

The Normalized Discrete Cosine Transform Shannon Entropy (DCTS) a helpful metric for quantifying image sharpness in the frequency domain. We used it to analyze image quality degradation vs. imaging depth (**Supplementary Fig. 17**). The definition of DCTS has been described in ref^54^, and we implemented it via customized MATLAB code.

#### Image contrast metric

We adopted a commonly used contrast metric – the root mean square (RMS) contrast (RMSC^55^) to quantify image contrast (**Supplementary Fig. 22**). The RMSC of an image is defined as:

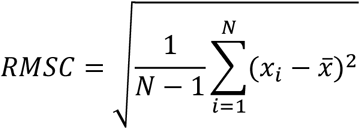

where *x*_*i*_ is the intensity of each pixel *i*, 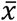 is the mean intensity of the image, and *N* is the total pixel number. To compare across different images, we first divide each image by its mean intensity and bin 3-fold to reduce noise before computing the RMSC.

#### Image intensity correction for time-lapse images

When applying the DeAbe model to predict images, the 3D-RCAN network automatically normalizes the input raw images to an intensity range of 0-1 by default. For time lapse images, this normalization was performed independently at each time point, resulting in additional intensity fluctuations. To compensate for these fluctuations, we applied corrections to the DeAbe predictions for the GCaMP calcium signal in the live worm experiments (**Supplementary Fig. 39, Supplementary Videos 15-17**).

We first calculated the normalization ratio at each time point:

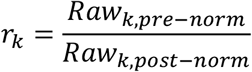

where *r*_*k*_ is the normalization ratio of time point *k*; *Raw*_*k,pre*−*norm*_ is the average intensity of the raw image volume before normalization and *Raw*_*k,post*−*norm*_ is the average intensity of the raw image volume after normalization.

Next, we rescaled the image intensity of the DeAbe images based on the normalization ratios by matching all time points to the first time point:

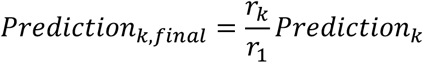

where *r*_1_ is the normalization ratio of the first time point; *Prediction*_*k*_ is the images predicted by DeAbe model; *Prediction*_*k,final*_ is the final images with intensity fluctuation compensation for quantitative analysis.

#### Calculation of vessel orientation and alignment

Orientations were estimated in 3D using a weighted vector summation algorithm^28^, adapting it for the volumetric images of fiber-like structures corresponding to the CD31 channel (i.e., blood vessel images) in iDISCO-cleared mouse embryos (**Fig. 4**).

For a given voxel within the 3D image, an *n* × *n* × *n* voxel window was generated surrounding the voxel under assessment. To segment the effective voxels, six-level Otsu intensity thresholding was applied to the image, with five thresholds dividing the intensity into six levels. The lowest level was designated as background noise, and regions assigned to the upper five levels defined the vessel signals. The window size *n* was typically set as two to three times the vessel thickness. All vectors passing through the center voxel were defined and weighted by their length and intensity variations, and the direction of the sum of all the weighted vectors was designated as the orientation of the center voxel^28^, with associated azimuthal angle *θ* (ranging from 0° to 180°) and polar angle *φ* (ranging from 0° to 180°). However, since the calculation of the polar angle *φ* was not straightforward, we defined two additional azimuthal angles, *β* and *γ* (**Supplementary Fig. 24a**), which were symmetrical to the azimuthal angle *θ*. *β* was defined as the angle between the projection of the vessel in the *zx* plane and the *x* axis, and *γ* was the angle between the projection in the *yz* plane and the − *y* axis. These two angles were related to the polar angle *φ* via:

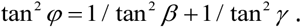

We also derived the 3D directional variance (DV) metric, quantifying the spread in orientations^29,56^. The value of DV ranges from 0 to 1, with 0 corresponding to perfectly parallel alignment, and 1 corresponding to complete disorder (**Supplementary Fig. 24b**). The directional variance 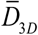 was defined as:

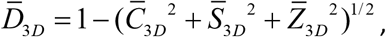

where:

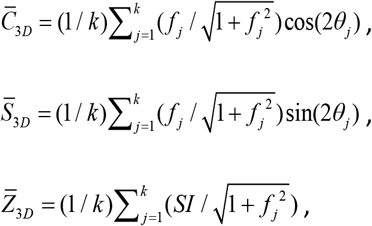

with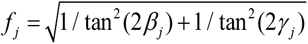, and *SI* = (−1) · (*φ* − 90) / |*φ* − 90|, where *φ* was acquired from the determination of *β* and *γ* as described above, *k* was the number of fiber voxels in the region, and *θ*, *β* and *γ* were calculated azimuthal angles as described above.

#### Membrane segmentation

For the images of live worm embryos dual-labeled with nuclear and membrane markers (**Fig. 5c, d, Supplementary Fig. 29**), raw data was restored using our multiple-step deep learning pipeline (Steps 1-3 in **Supplementary Fig. 26a**) prior to cell membrane segmentation. We performed automatic membrane segmentation using segmented nuclei as seeds:

First we used the Keras and Tensorflow-based implementations of Mask RCNN^57^ (https://github.com/matterport/Mask_RCNN) to perform nuclear segmentation (**Supplementary Fig. 29d**). We then manually segmented 8 volumes (3 acquired with diSPIM, 3 with iSPIM, and 2 from multiview confocal microscopy^27^ for a total of 1963 nuclei) for training. Of these 8 volumes, 6 volumes with a total of 1688 nuclei were used for training a segmentation network and 2 volumes with a total of 275 nuclei were used for validation. We used a ResNet-50 model as the backbone for our network, initialized the model using weights obtained from pretraining on the MS COCO dataset^58^, and proceeded to train all layers in three stages. Training took ∼10 hours and applying the model took ∼ 3 minutes per volume on a Windows workstation equipped with an Intel(R) Xeon(R) W-2145 CPU operating at 3.70 GHz, an Nvidia Quadro P6000 GPU, and 128 GB of RAM. After Mask RCNN segmentation, we applied a marker-controlled watershed operation (https://www.mathworks.com/help/images/marker-controlled-watershed-segmentation.html) to the nuclear segmentations to separate touching nuclei.

Second, we applied the vascular structure enhancement filter^59^ (https://github.com/timjerman/JermanEnhancementFilter) to the membrane data to enhance boundaries (**Supplementary Fig. 29c**). Scales were set to [2.0, 2.25, 2.5] and all other parameters were set to the default.

Third, the centroids of segmented nuclei were used as seeds, and we used the seeded watershed algorithm (https://github.com/danielsnider/Simple-Matlab-Watershed-Cell-Segmentation) for membrane segmentation (**Supplementary Fig. 29f**).

This workflow was applied both to the raw image data and restored images after each step in our multi-step pipeline to demonstrate the benefit of segmentation enhancement from DL processing.

For selected volumes (**Fig. 5c, Supplementary Video 11**), we also performed manual editing on the automatic segmentations produced by the multi-step deep learning pipeline. Manual editing was performed within the ImageJ plugin Labkit (https://imagej.net/plugins/labkit/). After automatic segmentations were imported to Labkit, segmentation labels were manually edited interactively in lateral views (XY planes), and then were edited in axial views (YZ planes). Since the manual editing was conducted in 2D views and initial editing in either view was not sufficient to ensure smoothness in 3D, we iterated twice to further improve our results.

## Code availability

Training and applying deep learning models were achieved using Python 3.7.0. Generation of synthetic aberrated data and quantitative image analysis was performed in MATLAB (Mathworks, R2022b). Customized code and software are available at https://github.com/eguomin/DeAbePlus/. RCAN and CARE software were installed from https://github.com/AiviaCommunity/3D-RCAN and https://github.com/CSBDeep/CSBDeep, and code for RL deconvolution and multiview fusion is available at https://github.com/eguomin/diSPIMFusion/.

## Data availability

The data that support the findings of this study are included in **Figs. 1-5, Supplementary Figs. 1–39** and **Supplementary Videos 1–17**. Some representative data from the figures (**Fig. 2a, Supplementary Figs. 16, 30**) are publicly available at https://doi.org/10.5281/zenodo.8424245. Other datasets (training data and intermediate data for deep learning) are available from the corresponding author upon reasonable request due to their large file size.

