## Supplementary File for "Deep learning-based aberration compensation improves contrast and resolution in fluorescence microscopy"

Supplementary Video Captions

**Supplementary Video 1, Lateral views of synthetic phantom data restored by DeAbe vs. other methods.** Phantoms consisting of randomly oriented and positioned dots, lines, spheres, spherical shells, and circles (ground truth, GT) were blurred to simulate microscopy data (Raw) and restored using blind deconvolution (Blind decon), Richardson-Lucy deconvolution with diffraction-limited PSF (RL Decon 1), Richardson-Lucy deconvolution with aberrated PSF (RL Decon 2), or our de-aberrating network (DeAbe). Lateral views though the volume are shown. Twenty iterations were used for RL deconvolution and 10 for blind deconvolution. See also **Fig. 1.**

**Supplementary Video 2, Axial views of synthetic phantom data restored by DeAbe vs. other methods.** As in **Supplementary Video 1**, but showing axial views through the volume.

**Supplementary Video 3, DeAbe restores images of *C. elegans* embryos expressing nuclear marker.** Images were acquired with single view light sheet microscopy (iSPIM, 1.1NA). Left: raw data, middle: same data after 10 iterations of Richardson-Lucy deconvolution (RL Decon), right: restoration after DeAbe. Lateral views through the image volume are shown. See also **Fig. 3a**.

**Supplementary Video 4, DeAbe restores images of adult *C. elegans* expressing NeuroPAL;GCaMP6s.** Images were acquired with spinning disk confocal microscopy. Left: raw data, right: restoration after DeAbe. Lateral views of individual color channels ($1^{st}$ – $3^{rd}$ row) and combined channels ($4^{th}$ row) through the image volume are shown. Note contrast has been increased to better visualize dim nuclei, this results in a large background in the red channel in the last few planes. See also **Supplementary Fig. 18**.

**Supplementary Video 5, DeAbe restores images of NK-92 cells fixed and stained with Alexa Fluor 555 wheat germ agglutinin.** Images were acquired with instant SIM. Volumetric maximum intensity projections (MIP), lateral views, and axial views through the image volume are shown in sequence. Left: raw data, middle: same data after restoration with DeAbe, right: same data after restoration with DeAbe+ (DeAbe followed by 20 iterations Richardson-Lucy deconvolution). See also **Fig. 3b-d**.

**Supplementary Video 6, DeAbe restores lateral views of live cardiac tissue expressing GFP-Tomm20.** Images were acquired with two photon microscopy. Left: raw data, middle: after 20 iterations of Richardson-Lucy deconvolution (RL Decon), right: restoration after DeAbe. Lateral views through the volume are shown. See also **Fig. 3e-f**.

**Supplementary Video 7, DeAbe restores axial views of fixed tissue expressing tdTomato membrane marker.** Images were acquired with two photon microscopy. Left: raw data, right: restoration after DeAbe.

**Supplementary Video 8, DeAbe ameliorates image degradation in mm-scale cleared mouse tissue embryo.** Fixed and iDISCO-cleared E11.5-day mouse embryo was immunostained for blood vessels (CD31, magenta) and neurons (TuJ1, cyan). Rendering compares raw and DeAbe+ (attenuation compensated, DeAbe, and deconvolved) data. See also **Fig. 4a-d**.

**Supplementary Video 9, DeAbe enhances quantification of vessel orientation and alignment in mm-scale cleared mouse tissue embryo.** Orientations (theta and phi) and 3D directional variance (DV) analysis on the blood vessel channel of the mouse embryo data in **Supplementary Video 8**. Rendering compares raw and DeAbe+ (attenuation compensated, DeAbe, and deconvolved) results. See also **Fig. 4e-g**.

**Supplementary Video 10, Multi-step deep learning restores images of *C. elegans* embryos expressing nuclear (magenta) and membrane (green) markers.** Images were acquired with single view light sheet microscopy (iSPIM, 1.1NA) and restored using a three-step deep learning pipeline. Maximum intensity projections (MIP, left column, shown only for nuclear channel) and single lateral plane 17.7μm into the volume (right column, both nuclei and membranes) are shown for raw (top row) and restored (bottom row) data. See also **Fig. 5a, b**.

**Supplementary Video 11, Multi-step deep learning improves image quality and cell segmentation for *C. elegans* embryos.** The $80^{th}$ time point volume (~320 min post fertilization) extracted from the time series data in **Supplementary Video 10**. Top row shows raw image (top left) and restored image (top right) using three-step deep learning pipeline, with nuclei in magenta and membrane in green. Bottom row shows the automated cell segmentation based on the raw image (bottom left) and the automated cell segmentation followed by manual editing based on restored image (bottom right). 319 cells are automatically segmented in the raw data, but 421 cells are segmented based on manual editing of the multi-step deep learning result. Lateral views through the image volume are shown. See also **Fig. 5c**.

**Supplementary Video 12, Multi-step deep learning restores images of *C. elegans* embryos expressing ttx-3B-GFP, marking neurons and gut cells.** Raw images (left) were acquired with single view light sheet microscopy (iSPIM, 0.8NA) and restored using three-step deep learning pipeline. Maximum intensity projections through time are shown; selected time points also show volumetric projections. See also **Fig. 5i**.

**Supplementary Video 13, Multi-step deep learning restores images of *C. elegans* embryos expressing nuclear marker.** Images were acquired with single view light sheet microscopy (iSPIM, 1.1NA) and restored using three-step deep learning pipeline. From left to right were shown the raw images, Step 1 DeAbe, Step 2 Decon, and Step 3 Expan results. Lateral views through the volume are shown.

**Supplementary Video 14, Multi-step deep learning restores time-lapse images of *C. elegans* embryos expressing nuclear marker.** Images were acquired with single view light sheet microscopy (iSPIM, 1.1 NA) and restored using three-step deep learning pipeline. From left to right were shown the raw images, Step 1 DeAbe, Step 2 Decon, and Step 3 Expan results. Maximum intensity projections through time are shown.

**Supplementary Video 15, DeAbe restores highly dynamic time-lapse images of live *C. elegans* expressing a GCaMP marker targeted to neurons.** Images were acquired with instant SIM (1.15 NA) at 1.5 volumes/s over 1000 time points. Top: raw data; Bottom: restoration after DeAbe. Maximum intensity projections through time are shown.

**Supplementary Video 16, Higher magnification GCaMP data shown in Supplementary Video 15.** Images from the same dataset as **Supplementary Video 15**, but higher magnification views of the nerve ring region from the period spanning 9.3 – 54.7 s, highlighting the rapid, fine details otherwise obscured in the raw images.

**Supplementary Video 17, Additional higher magnification GCaMP data shown in Supplementary Video 15.** Images from the same dataset as **Supplementary Video 15**, but higher magnification views of the nerve ring region from another period spanning 423.5 – 493.6 s, highlighting the rapid, fine details otherwise obscured in the raw images.

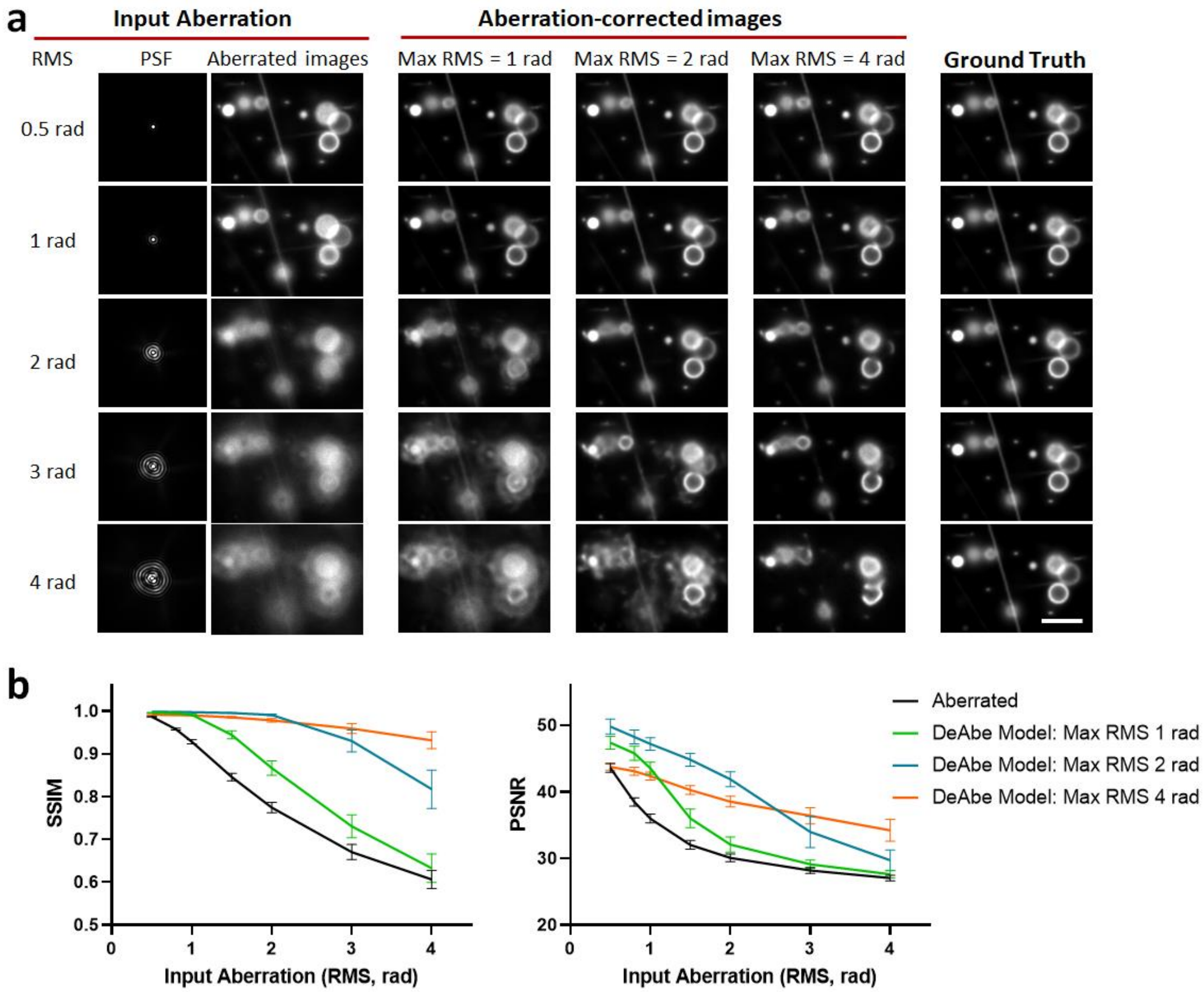


**Supplementary Fig. 1, DeAbe performance depends on aberration magnitude in input images and training data. A)** Example aberrated PSFs and associated aberrated input images (left); DeAbe model predictions yielding aberration corrected images (middle), shown for different training data consisting of mixed aberrations with indicated maximum root mean square (RMS) wavefront distortion (see also **Methods**); and ground truth synthetic objects (right). The example input aberration here is defocus, applied at an increasing aberration magnitude (0.5 rad – 4 rad RMS wavefront distortion). **b)** Quantification of **a)**. Structural similarity index (SSIM) and peak signal-to- noise ratio (PSNR) metrics are used to quantify the quality of DeAbe Net predictions under different training data regimes (green, blue, red curves at 1, 2, and 4 rad max RMS) vs. ground truth. All DeAbe predictions show improvement compared to input aberrated image (black), with training data containing larger magnitude aberrations better able to compensate for larger aberration magnitudes in the input images. For this work, we used the model corresponding to a maximum RMS wavefront distortion of two radians (blue curve). Means and standard deviations are shown from 100 independent simulations, each with randomized object structures and randomized input aberrations. Scale bar: 5 μm.

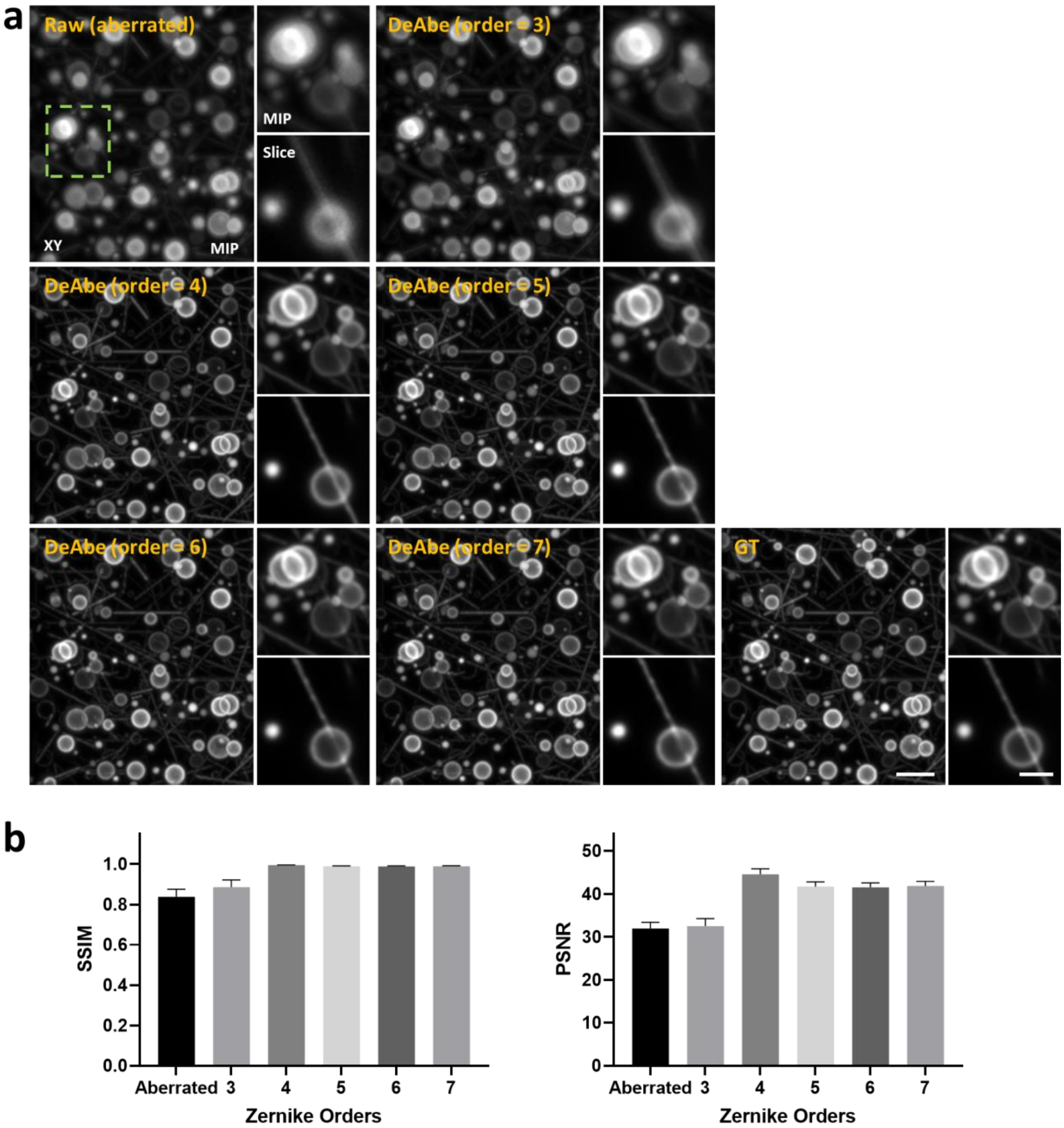


**Supplementary Fig. 2, Dependence of DeAbe prediction on number of Zernike orders in training data.** Synthetic aberrated phantom structures in **Fig. 1b** were input into DeAbe networks trained with progressively more Zernike basis aberrations. Visual **a)** and quantitative **b)** analysis indicate that performance plateaus after the 4$^{th}$ Zernike terms are added. Thus, in this work, we used four Zernike orders to generate training data. In **b)**, both SSIM (left) and PSNR (right) values are shown from 100 independent simulations. Scale bars: 5 μm and 2.5 μm (insets).

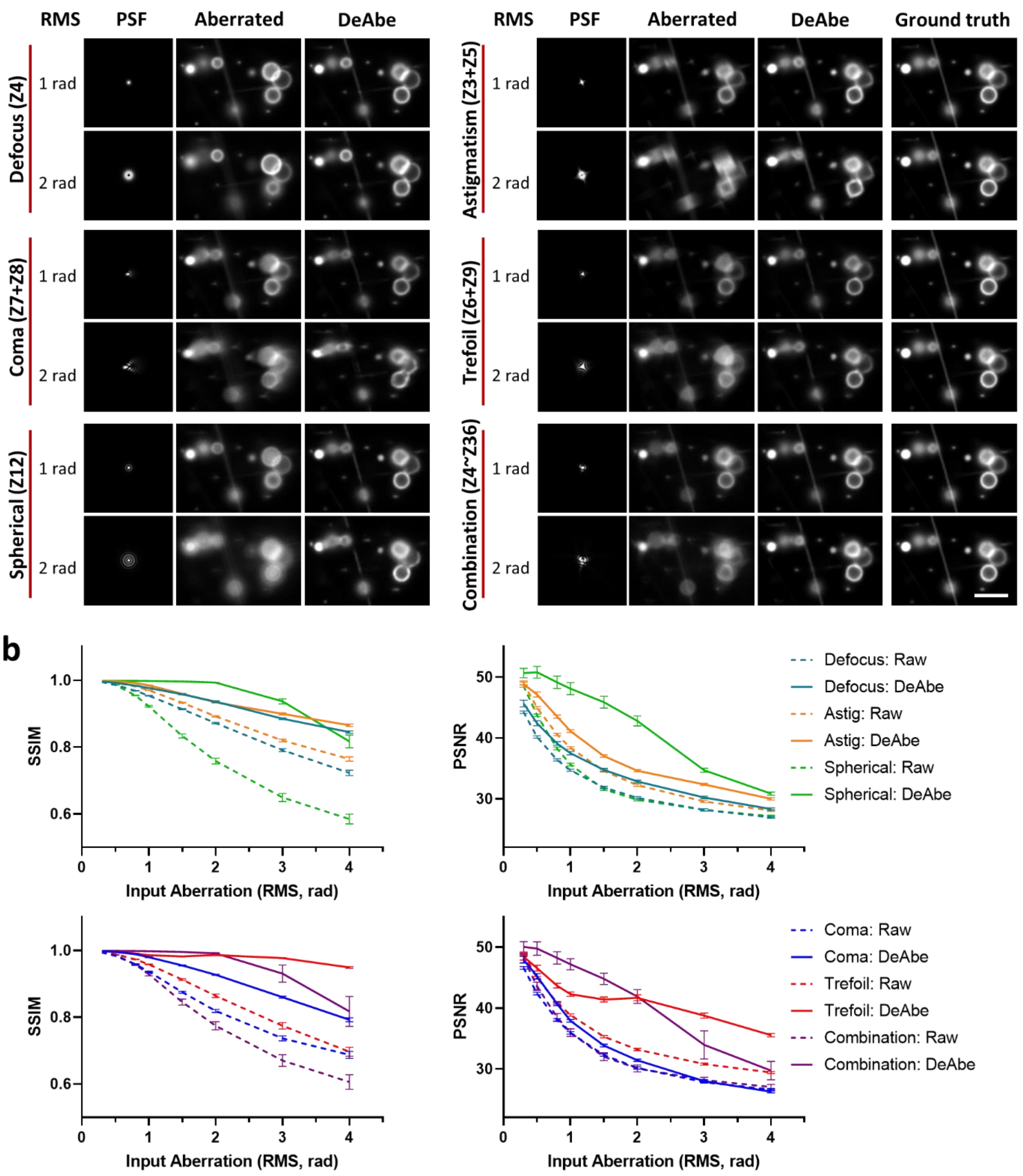


**Supplementary Fig. 3, DeAbe network predictions on images contaminated with specific aberration modes.** Synthetic phantoms were aberrated with defocus, coma, spherical, astigmatism, trefoil, and mixed aberrations with indicated RMS wavefront distortion. A DeAbe model trained on data contaminated with mixed aberrations (up to fourth order Zernike basis terms, maximum RMS wavefront distortion two radians) was then used to compensate for aberrations. **a)** Aberrated PSFs (left), aberrated images (middle) and DeAbe model predictions (right) are shown for each condition, as well as the ground truth reference (GT). Z in each vertical axis label refers to the Zernike order. **b)** Quantification using SSIM and PSNR metrics (means and standard deviations from 100 independent simulations) support the visual improvement after application of the DeAbe model. Scale bar: 5 μm.

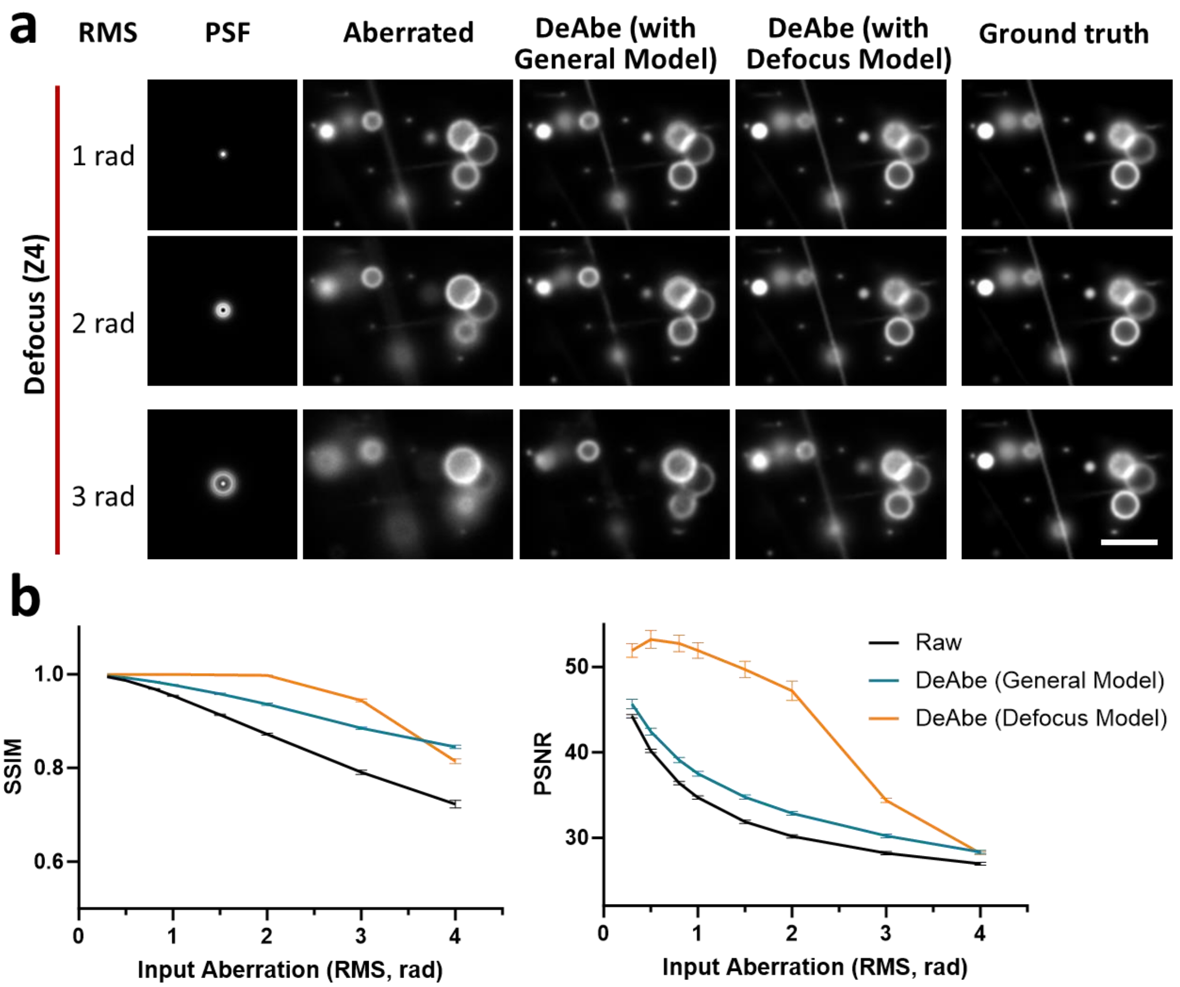


**Supplementary Fig. 4, DeAbe network predictions on defocused images improve if using a dedicated network trained purely on defocused images.** Synthetic phantoms were aberrated with defocus with indicated RMS wavefront distortion. A general DeAbe model trained on data contaminated with mixed aberrations (up to fourth order Zernike basis terms, maximum RMS wavefront distortion two radians), or a specific defocus DeAbe model trained on data contaminated with only defocus aberrations (maximum RMS wavefront distortion two radians), was then used to compensate for aberrations. **a)** From left to right: aberrated PSFs, aberrated images, predictions with a general DeAbe model, predictions with a specific defocus DeAbe model, and the ground truth reference (GT). **b)** Quantification using SSIM and PSNR metrics (means and standard deviations from 100 independent simulations) to compare the performance of the general DeAbe model vs. the specific defocus DeAbe model. Scale bar: 5 μm.

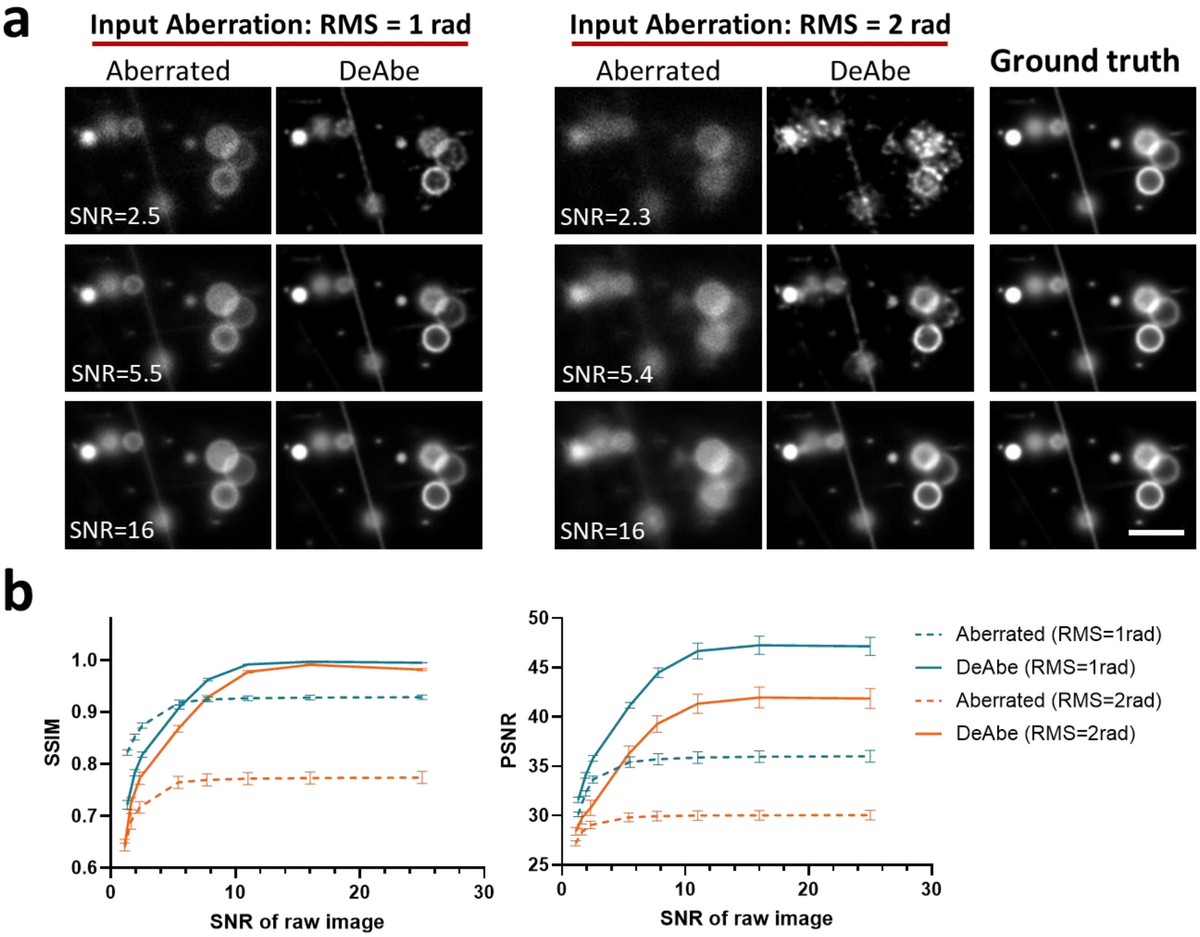


**Supplementary Fig. 5, Effect of noise on DeAbe model prediction. a)** Images of synthetic phantom structures with aberrations were additionally contaminated with noise to simulate different SNR levels and degraded with indicated RMS wavefront distortion. Ground truth structure (GT) is shown for comparison. **b)** SSIM and normalized PSNR analysis (means and standard deviations from 100 independent simulations) for data in **a),** confirming that performance of DeAbe network deteriorates in the presence of increasing noise and aberration. Scale bar: 5 μm.

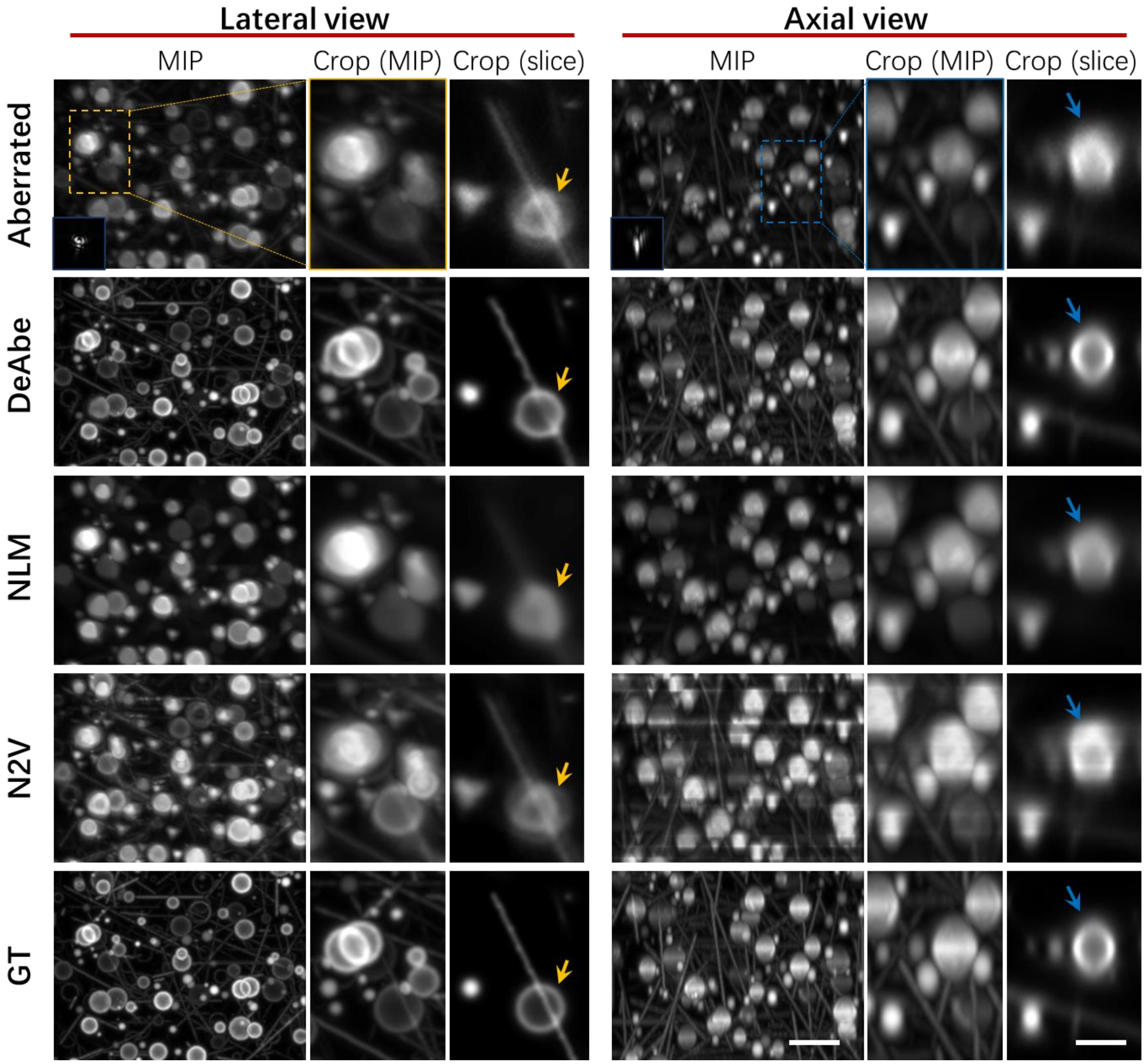


**Supplementary Fig. 6, DeAbe outperforms denoising methods.** Synthetic phantoms (Row 5, GT) were aberrated (Row 1) and subsequently restored with DeAbe (Row 2), nonlocal means (NLM, Row 3), or noise to void (N2V, Row 4). Lateral (left columns) or axial (right columns) are shown; orange and blue arrows highlight features better resolved with DeAbe than the denoising methods. Noise has been added to match the 'SNR 16' condition in **Supplementary Fig. 5.** Scale bars: 5 µm and 2.5 µm (insets). Data shown are representative results from N = 100 independent simulations.

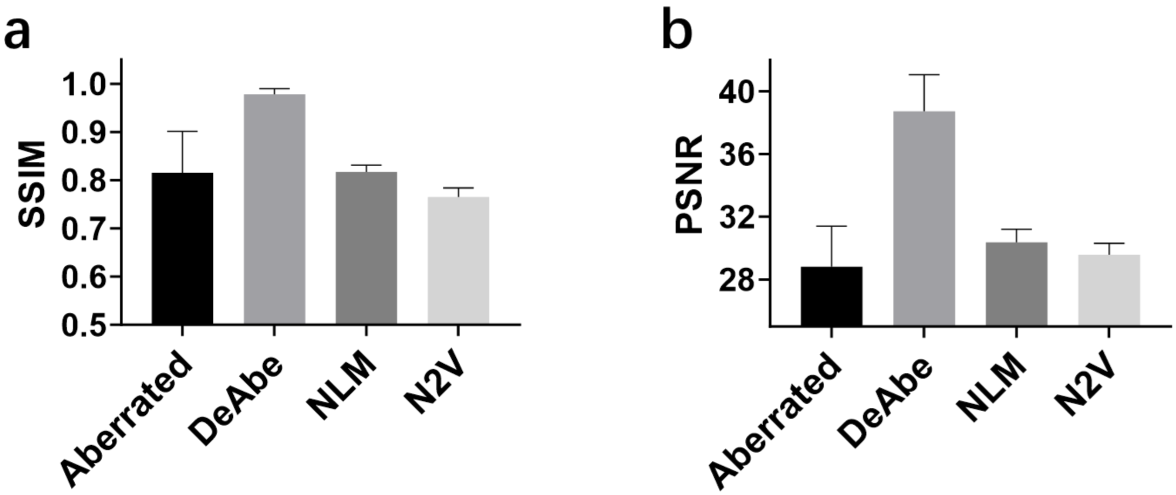


**Supplementary Fig. 7**, **Quantification of DeAbe vs. other denoising methods and aberrated input, to accompany Supplementary Fig. 6**. DeAbe offers higher structural similarity index (SSIM, **a)**) and peak signal-to-noise ratio (PSNR, **b)**) than other methods or aberrated input. Means and standard deviations from N = 100 volumes are shown.

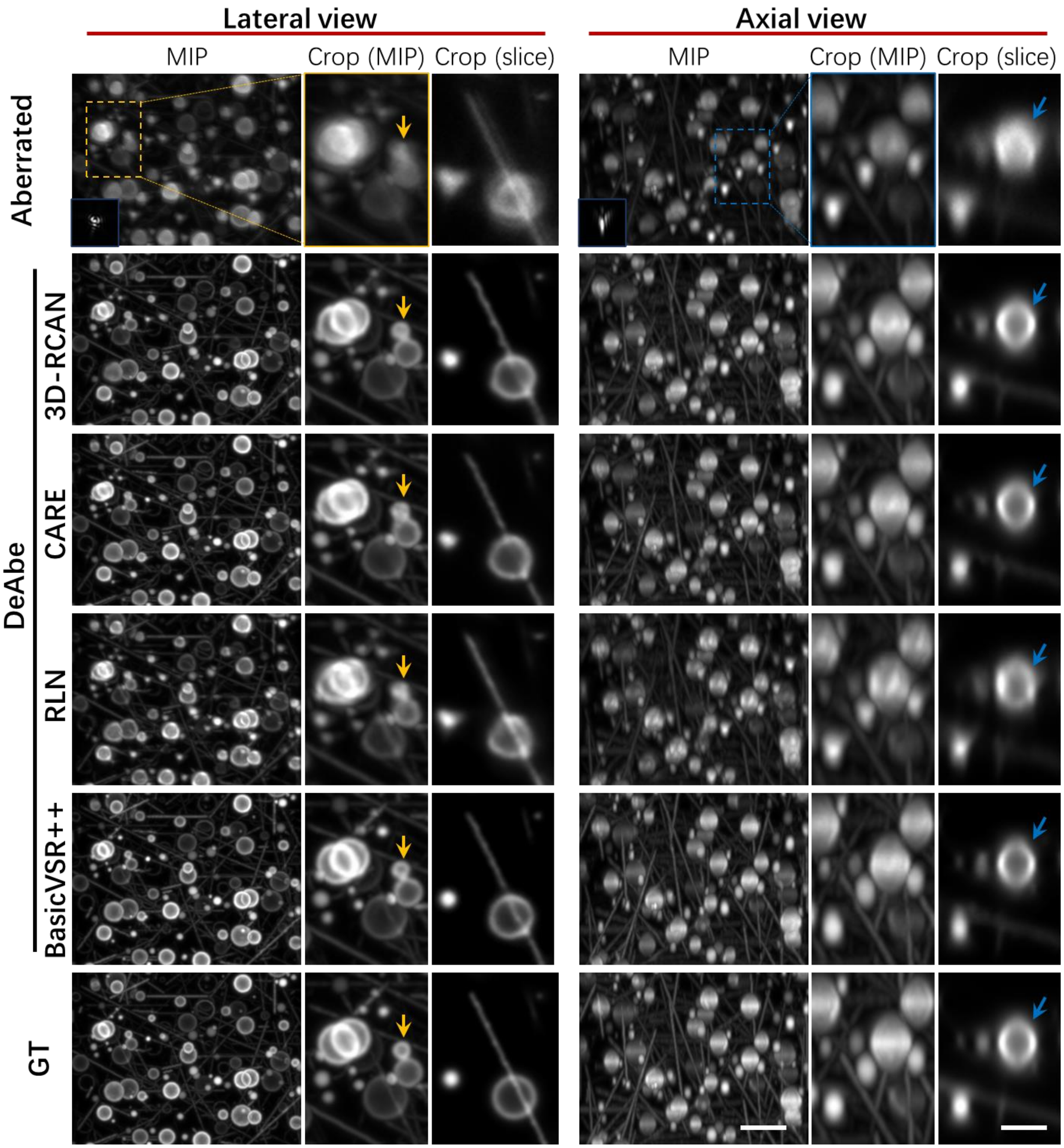


**Supplementary Fig. 8, The RCAN network compares favorably to other network architectures in implementing DeAbe.** Synthetic phantoms were aberrated and 3D-RCAN, CARE, RLN, and BasicVSR++ networks were used to restore the data (rows). Ground truth is also shown for a comparison. Lateral (left columns) and axial (right columns) are shown, including higher magnification views of orange and blue rectangular regions in lower magnification views. Maximum intensity projections (MIPs) and single planes (slice) are shown, arrows are intended to guide the eye for visual comparisons. Scale bar: 5 µm and 2.5 µm (insets). Data shown are representative results from N = 100 independent simulations.

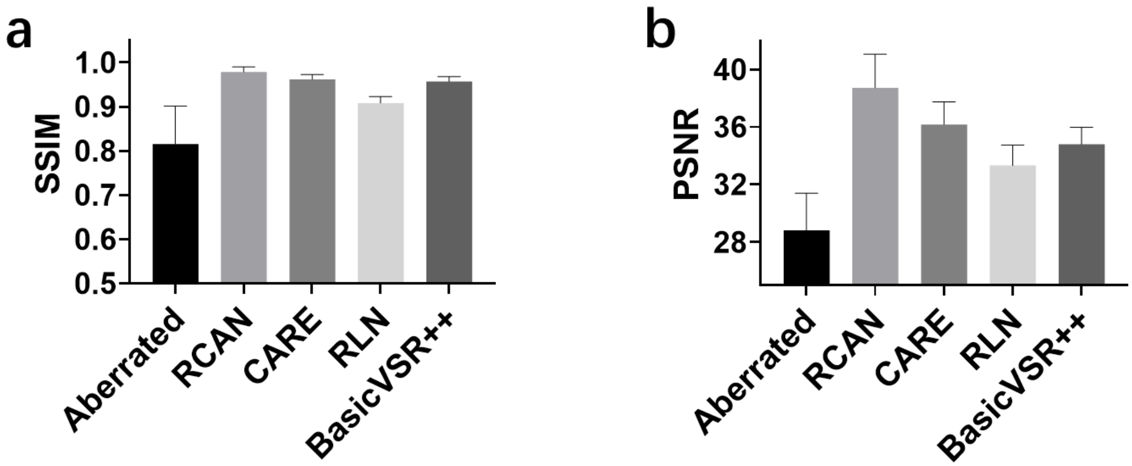


**Supplementary Fig. 9, Quantification corresponding to Supplementary Fig. 8**, showing that RCAN-based DeAbe offers improved SSIM **a)** or PSNR **b)** relative to CARE, RLN, or BasicVSR++ networks. Means and standard deviations from N = 100 3D measurements are shown.

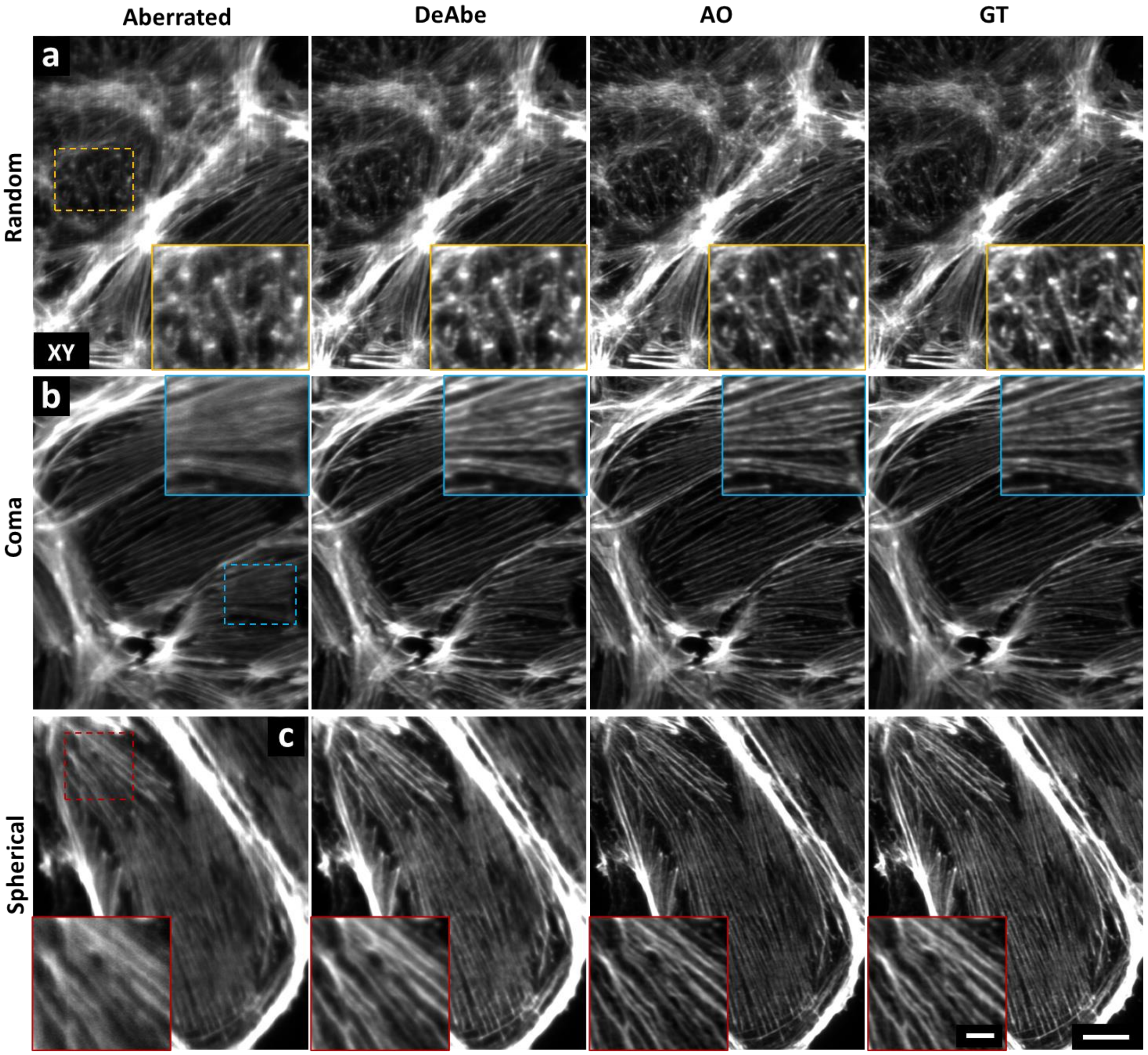


**Supplementary Fig. 10, Additional examples of DeAbe predictions on aberrated images.** PtK2 cells were fixed, stained for actin, and imaged with an AO lattice light sheet microscope. Random **a)**, coma **b)**, or spherical **c)** aberrations were applied (first column) and images were processed via DeAbe (second column, trained on random aberrations) or AO (third column). Ground truth (GT, fourth column) is also shown. Insets correspond to higher magnification views in first column. Scale bars: 10 µm and 3 µm (insets). Data shown are representative samples from N = 12 experiments.

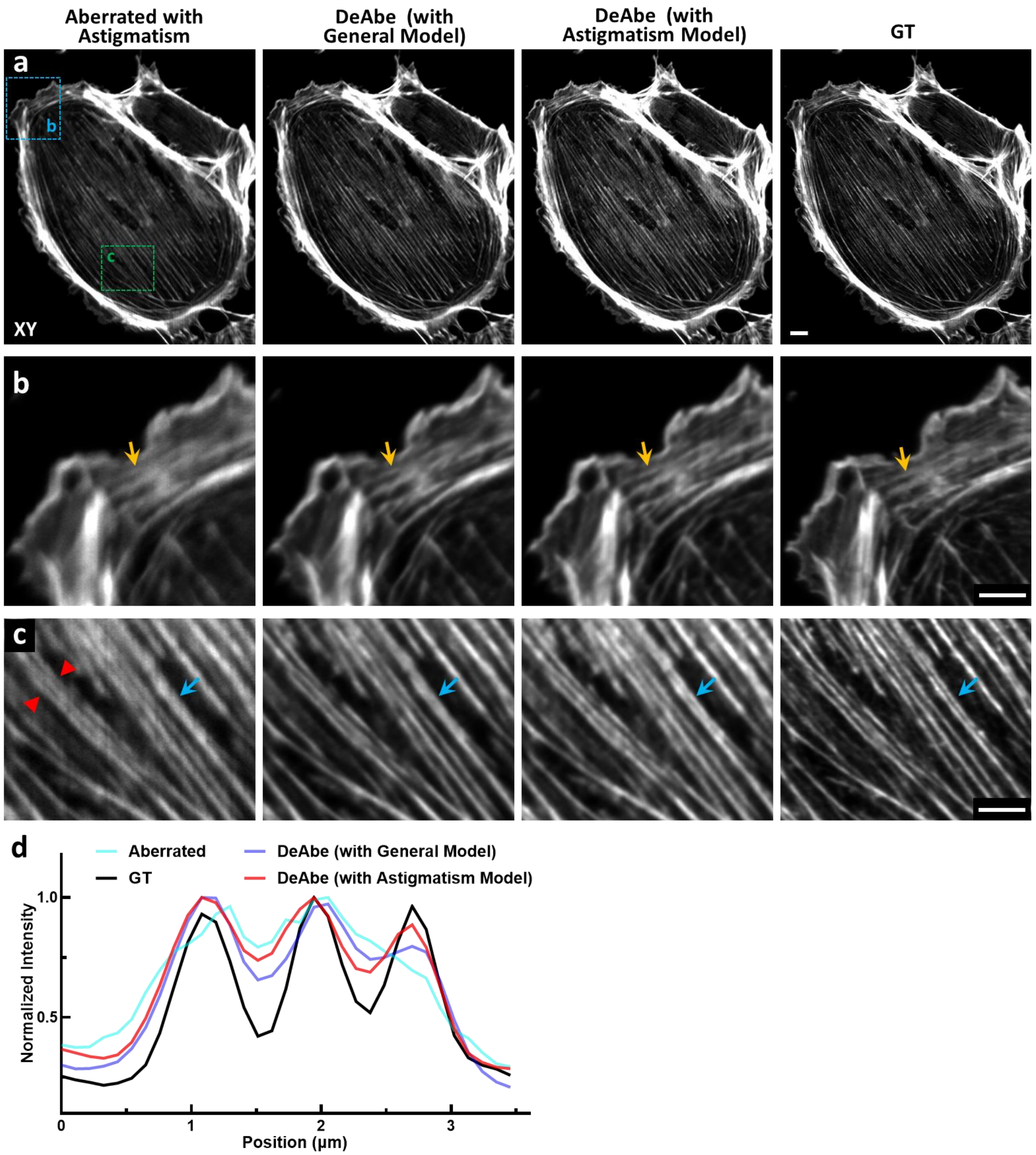


**Supplementary Fig. 11, Training DeAbe with specific aberrations improves performance relative to using a general model trained on a mix of aberrations.** PtK2 cells were fixed, stained for actin, and imaged with an AO lattice light sheet microscope. **a)** Left to right: Aberrated image (aberration is astigmatism), DeAbe prediction with a model trained on a mixture of aberrations, DeAbe prediction with a model trained only with astigmatism, and ground truth (GT). **b, c)** Higher magnification views of blue and green rectangular regions in **a)**. Yellow and blue arrows highlight features for visual comparisons. **d)** Line profile across region demarcated by red arrowheads in **c)**. Scale bars: 10 µm **a**); 5 µm **b, c**). Data shown are representative samples from N = 3 experiments.

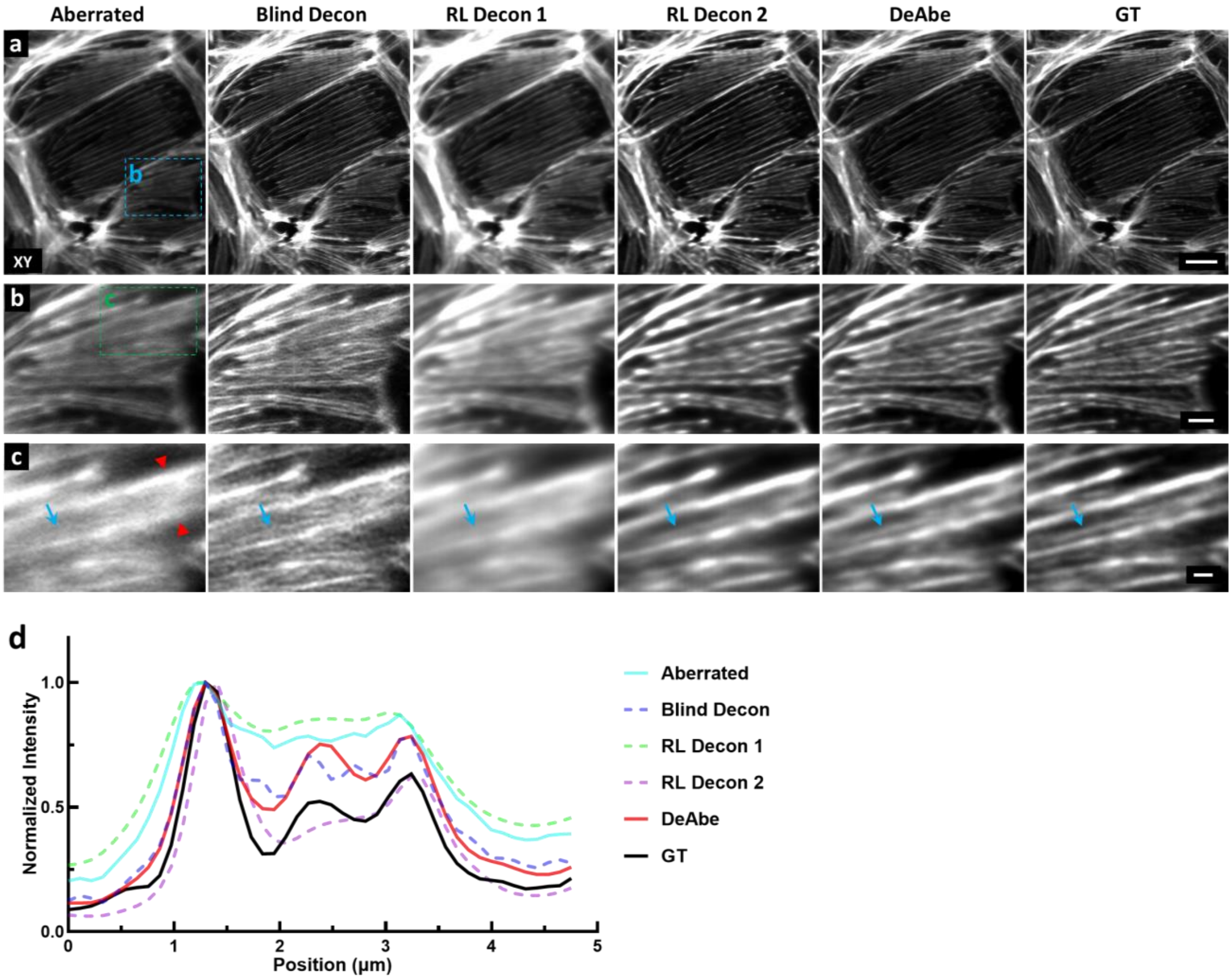


**Supplementary Fig. 12, Comparing DeAbe with deconvolution on experimental images.** PtK2 cells were fixed, stained for actin, and imaged with an AO lattice light sheet microscope. **a)** Left to right: Image aberrated with astigmatism, the result of blind deconvolution (Blind Decon), after deconvolution with an ideal PSF (RL Decon 1), after deconvolution with the aberrated PSF corresponding to the aberrated wavefront (RL Decon 2, note this is impractical in most applications as the actual PSF is usually difficult to obtain), DeAbe prediction using a general model trained on a random mixture of aberrations, and ground truth (GT, no aberrations added). **b)** Higher magnification views of the blue rectangular region in **a)**. **c)** Higher magnification views of the green rectangular region in **b)**. **d)** Line profile along region delineated by red arrowheads in **c)**, showing that DeAbe model trained with astigmatism resolves filament triplet, unlike deconvolution methods. Blue arrows emphasize features better resolved in DeAbe prediction vs. deconvolution. All deconvolution was performed with 10 iterations. Scale bars: 10 µm **a**); 3 µm **b**); 1 µm **c**). Data shown are representative samples from N = 3 experiments.

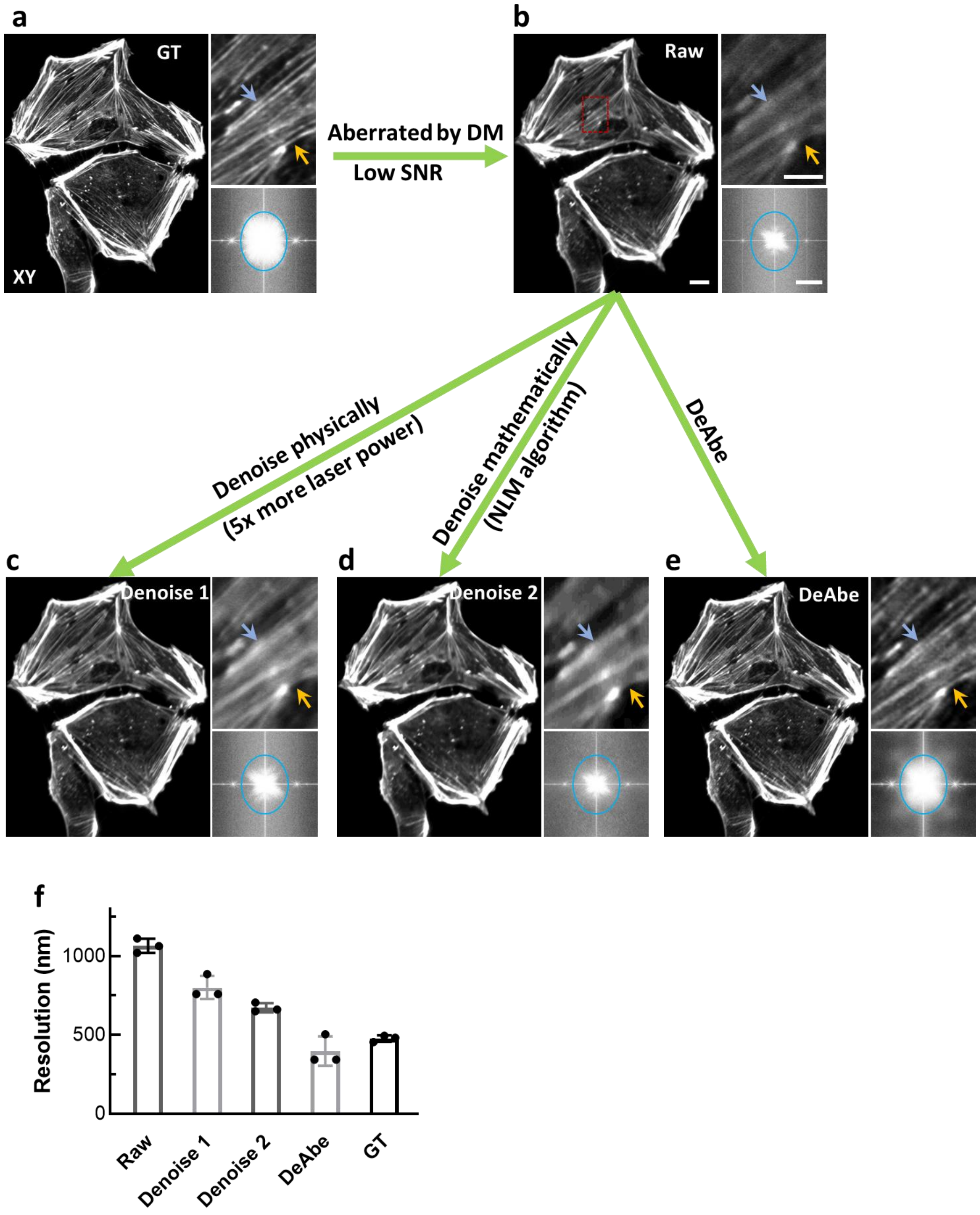


**Supplementary Fig. 13, DeAbe outperforms denoising methods on experimental data.** PtK2 cells were fixed, stained for actin, and imaged with an AO-lattice light sheet microscope at high SNR **a)** and aberrated and imaged at low SNR **b)**. We then denoised the low SNR, aberrated data physically by imaging at 5x higher laser power ('Denoise 1') **c)**; denoised digitally ('Denoise 2', via nonlocal means, **(d)**), or applied DeAbe after training with random aberrations **e)**; DeAbe provides better image restoration than either of the denoising methods. Insets at top right: higher magnification views of the dashed rectangle in **b)**. Arrows highlight actin fibers better resolved with DeAbe than via denoising methods. Insets at right bottom: Fourier transforms (FTs) of corresponding images, with blue circles indicate spatial frequency extents of 1/500 $nm^{-1}$ horizontally and 1/400 $nm^{-1}$ vertically. **f)** Decorrelation resolution analysis of the images. Means, standard deviations and individual data points from N = 3 images are shown. Scale bars: 10 μm; 5 μm in top right insets. Data shown are representative samples from N = 3 experiments.

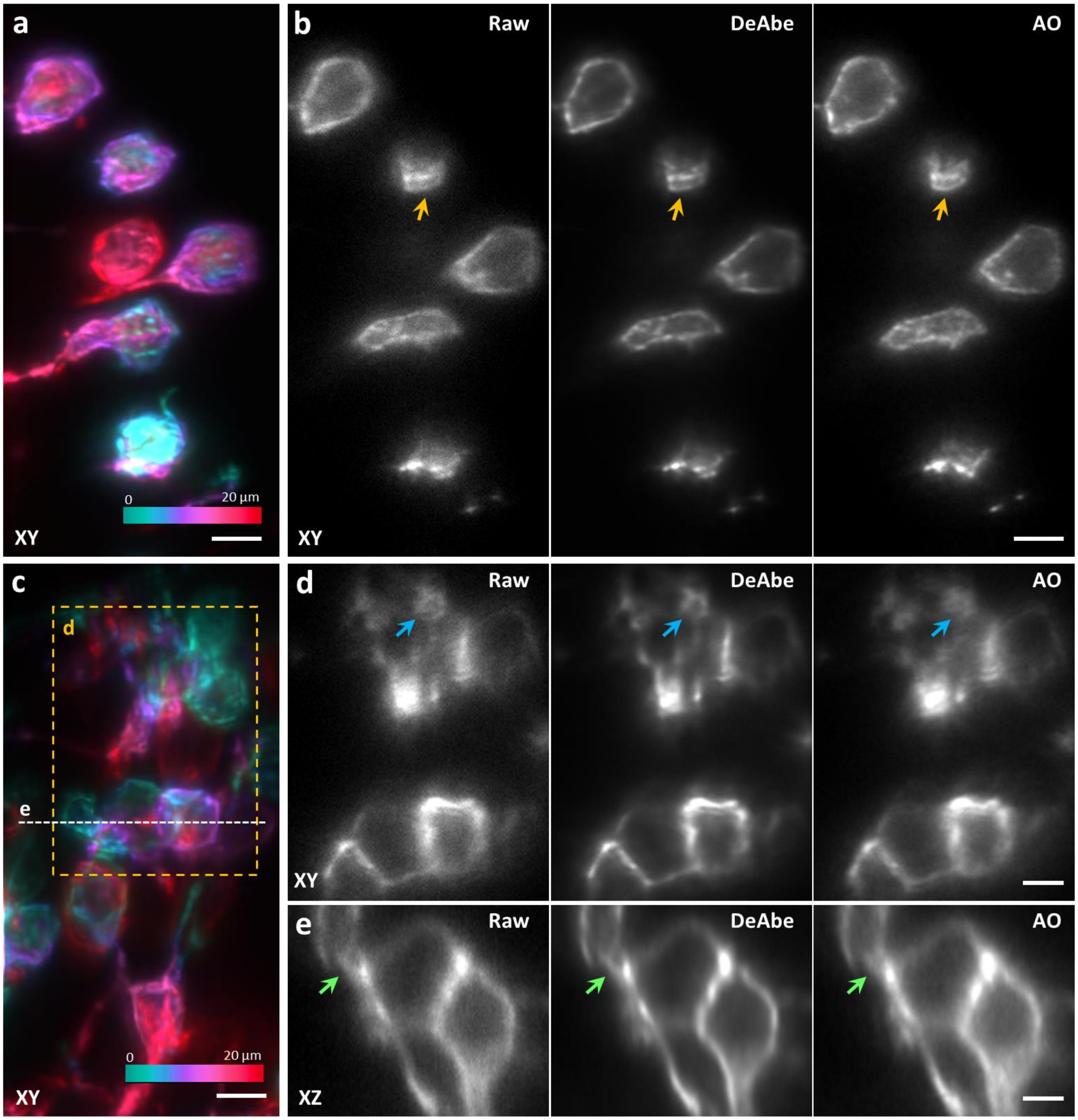


**Supplementary Fig. 14, Additional examples showing that DeAbe compares favorably to AO correction on zebrafish samples.** As in **Fig. 2,** 5 dpf zebrafish embryos expressing a GFP membrane marker labeling glutamatergic neurons were fixed and imaged in an AO-lattice light sheet microscope. Image volumes were collected 40-140 µm from the surface of the fish and passed through DeAbe or corrected via AO. **a)** Volume spanning 20 µm. Depth coded lateral (XY) maximum intensity projection. **b)** Single lateral plane 11.6 µm into imaging volume. Left to right: raw data, DeAbe prediction, and AO correction are shown. **c)** Another depth coded lateral maximum intensity projection from a different acquisition. Higher magnification lateral view (single plane) **d)** and axial view (single plane, (XZ)) **e)** views corresponding to orange rectangular region and dashed white line in **c)**, comparing raw, DeAbe prediction, and AO correction. Arrows in **b, d, e)** highlight membrane regions for comparisons. Scale bars: 5 µm **a, b, c**); 2 µm **d, e**). Data shown are representative samples from N = 15 experiments.

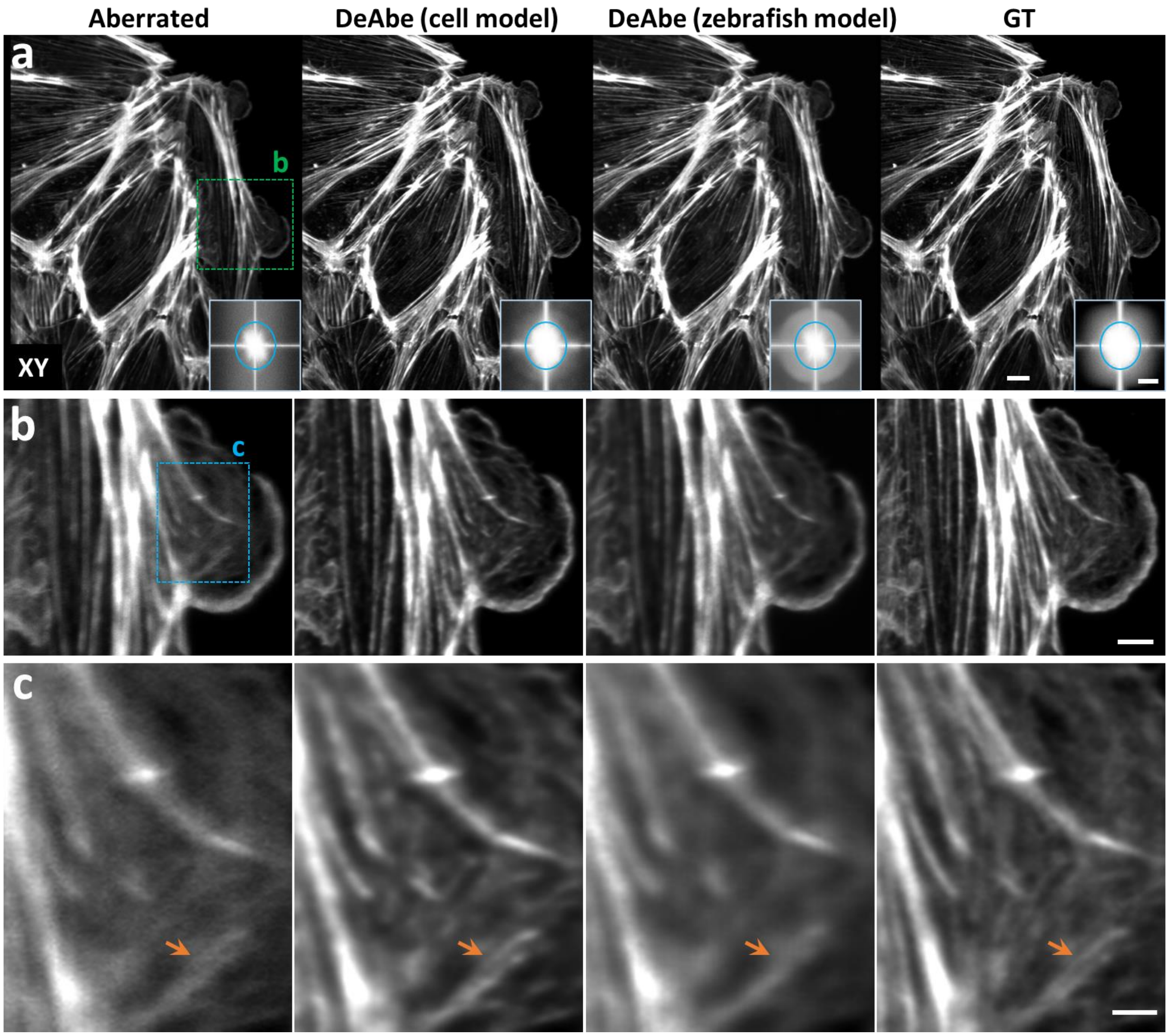


**Supplementary Fig. 15, Model generalization test on experimental dataset.** Fixed PtK2 cells were stained for actin and imaged in an AO lattice light sheet microscope. **a)** Left to right: Aberrated image, DeAbe prediction with model trained on similarly stained cells, DeAbe prediction applying the model trained on the zebrafish membrane marker (**Fig. 2g-m, Supplementary Fig. 14**), and ground truth (GT). **b, c)** Higher magnification views of dashed rectangular regions in **a, b)** respectively. As is clear in higher magnification views (e.g., orange arrows), it is best to train models that are matched to the data. Scale bars: 10 µm **a**) and 0.4 $\mu m^{-1}$ vertical/ 0.5 $\mu m^{-1}$ horizontal (insets); 5 µm **b**); 2 µm **c**). Data shown are representative samples from N = 12 experiments.

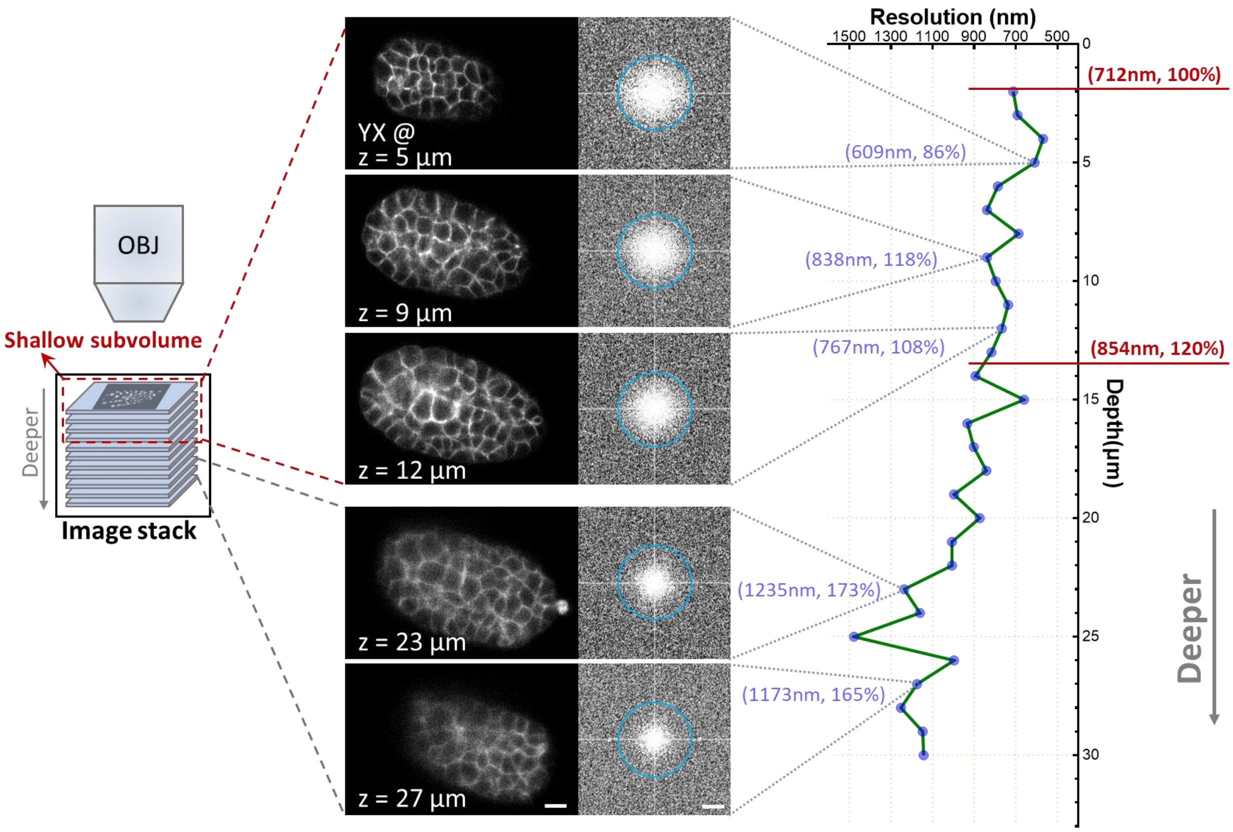


**Supplementary Fig. 16, Shallow subvolume selection.** Selected planes at different depths of an image stack from *C. elegans* embryos expressing membrane marker and accompanying decorrelation analysis showing degradation in spatial resolution as a function of depth. Right-hand side shows resolution estimated by decorrelation analysis as a function of depth (lower values are better), with absolute values and %age of initial value shown in blue and red. The middle column shows lateral views and corresponding Fourier transforms with progressive degradation in image quality from the shallow side (closest to the detection objective) to the deeper side of the stack due to aberrations. The slices on shallow side (referred to as a "shallow subvolume") are manually selected as ground truth and used to generate synthetically aberrated images. The cutoff chosen in this experiment corresponds to plane at which resolution degrades by 20%. Blue circles in right panel indicate 1/0.6 $\mu m^{-1}$ spatial frequency. Scale bars: 5 μm (left) and 1/2.5 $\mu m^{-1}$(right). Data shown are representative samples from N = 3 experiments.

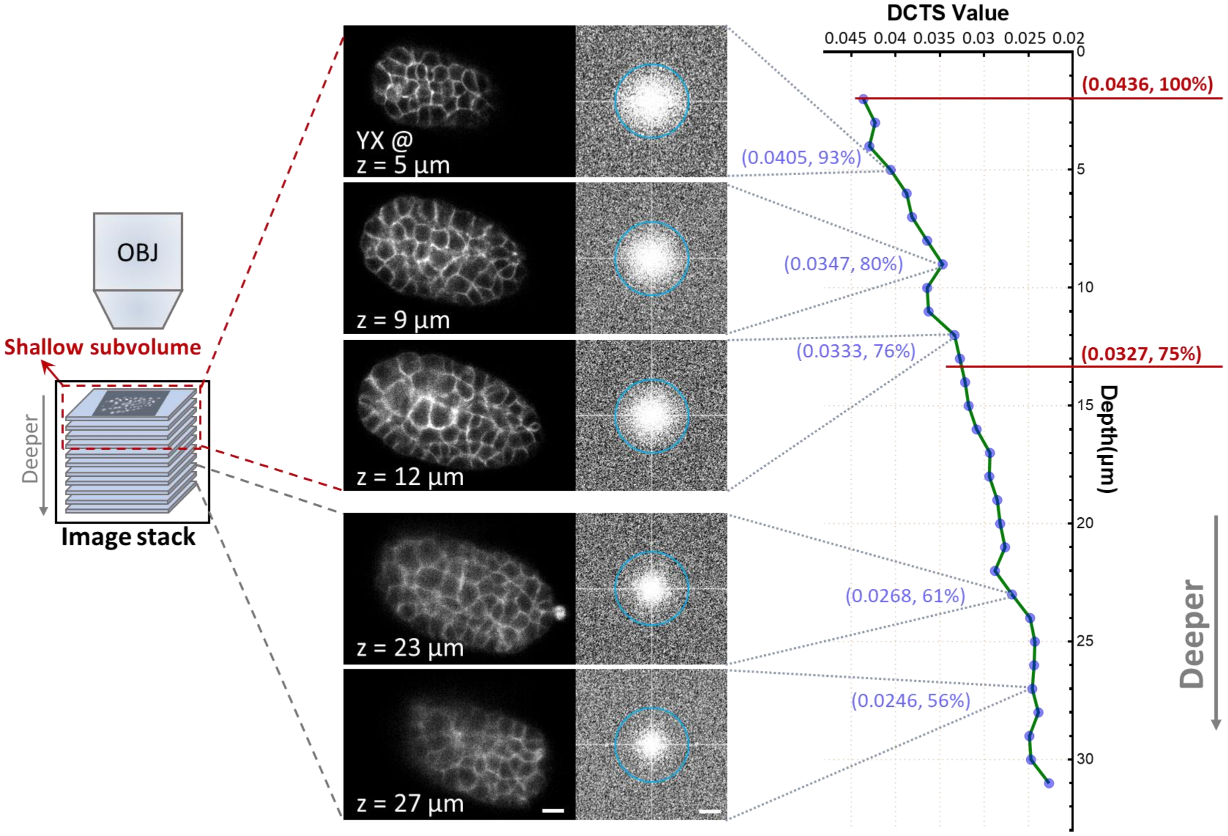


**Supplementary Fig. 17, As in Supplementary Fig. 16, but using Normalized Discrete Cosine Transform Shannon Entropy (DCTS) as a measure of degradation.** Right-hand side shows DCTS (higher values are better) as a function of depth, with absolute values and %age of initial value shown in blue and red. Blue circles in the right panel indicate 1/0.6 $\mu m^{-1}$ spatial frequency. Red lines indicate DCTS value at shallowest plane (top) and at the same plane as in **Supplementary Fig. 16** (bottom, corresponding to value at which resolution degrades by 20%). Scale bars: 5 μm (left) and 1/2.5 $\mu m^{-1}$(right). Data shown are representative samples from N = 3 experiments.

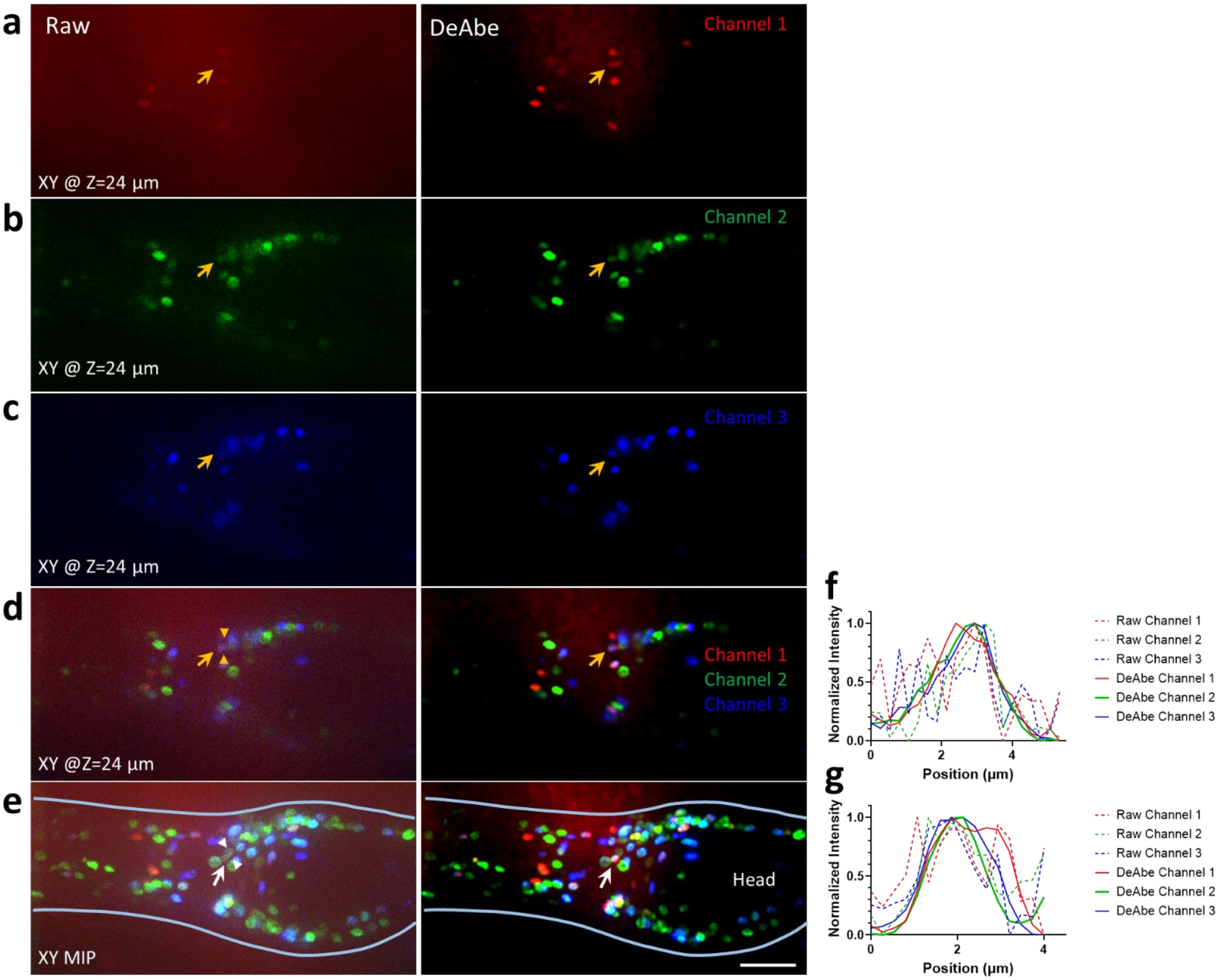


**Supplementary Fig. 18, DeAbe compensates for aberrations in spinning-disk confocal microscopy.** Adult *C. elegans* expressing NeuroPAL;GcaMP6s was imaged with spinning-disk confocal microscopy (**a-e)**, left) and our DeAbe model used to restore the data (**a-e)**, right). Individual planes 24 µm from the surface of the worm are shown in each color channel **a-c)**), in merged overlay **d)**, or a maximum intensity projection (MIP) over the whole volume **e)** are shown. In the MIP, the boundary of the worm has also been traced (blue lines) and the head region indicated for context. Orange (**a-d**) or white **e)** arrows highlight nuclei; double-headed arrowheads show line profiles (**f, g**) over the same nuclei comparing raw images (dashed lines) to DeAbe model prediction (solid lines). The network prediction improves signal-to-noise ratio and contrast in each channel compared to the raw data. Intensity profiles in (**f, g**) are each normalized to the maximum value in their profile. See also **Supplementary Video 4**. Scale bar: 20 µm. Data shown are representative samples from N = 3 experiments.

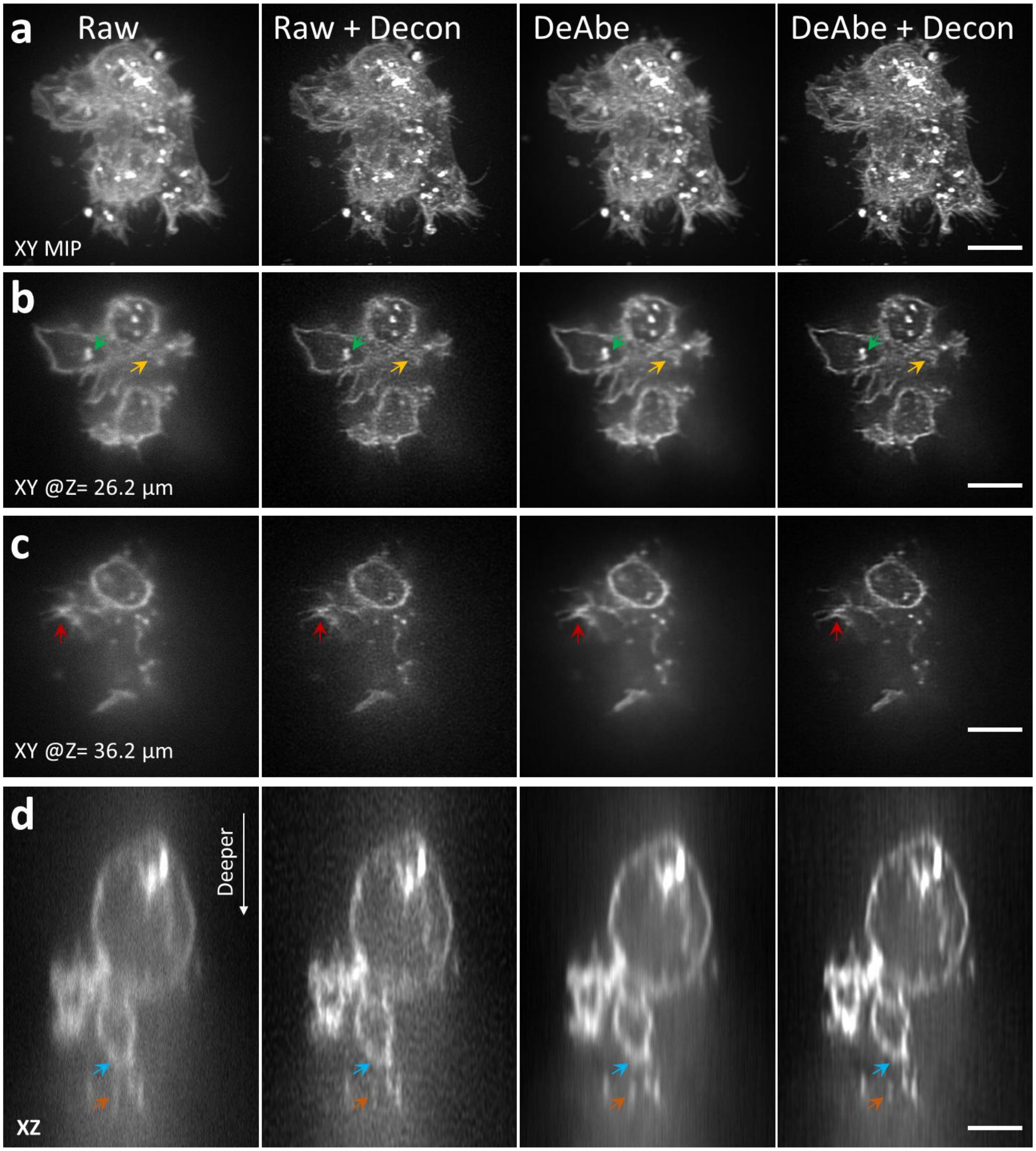


**Supplementary Fig. 19, Progressive improvements in image quality in NK-92 cells imaged with iSIM.** Cells were fixed and stained for wheat germ agglutinin as in **Fig. 3b-d**. Columns show (left to right) representative images of raw data, deconvolved data, DeAbe prediction on raw data, and DeAbe prediction followed by deconvolution (20 iterations Richardson-Lucy). The latter provides the clearest lateral (**a-c**) and axial (**d)** views, best highlighting fine features in the data (arrows). MIP: maximum intensity projection. Scale bars: 5 µm. See also **Supplementary Video 5**. Data shown are representative samples from N = 3 experiments.

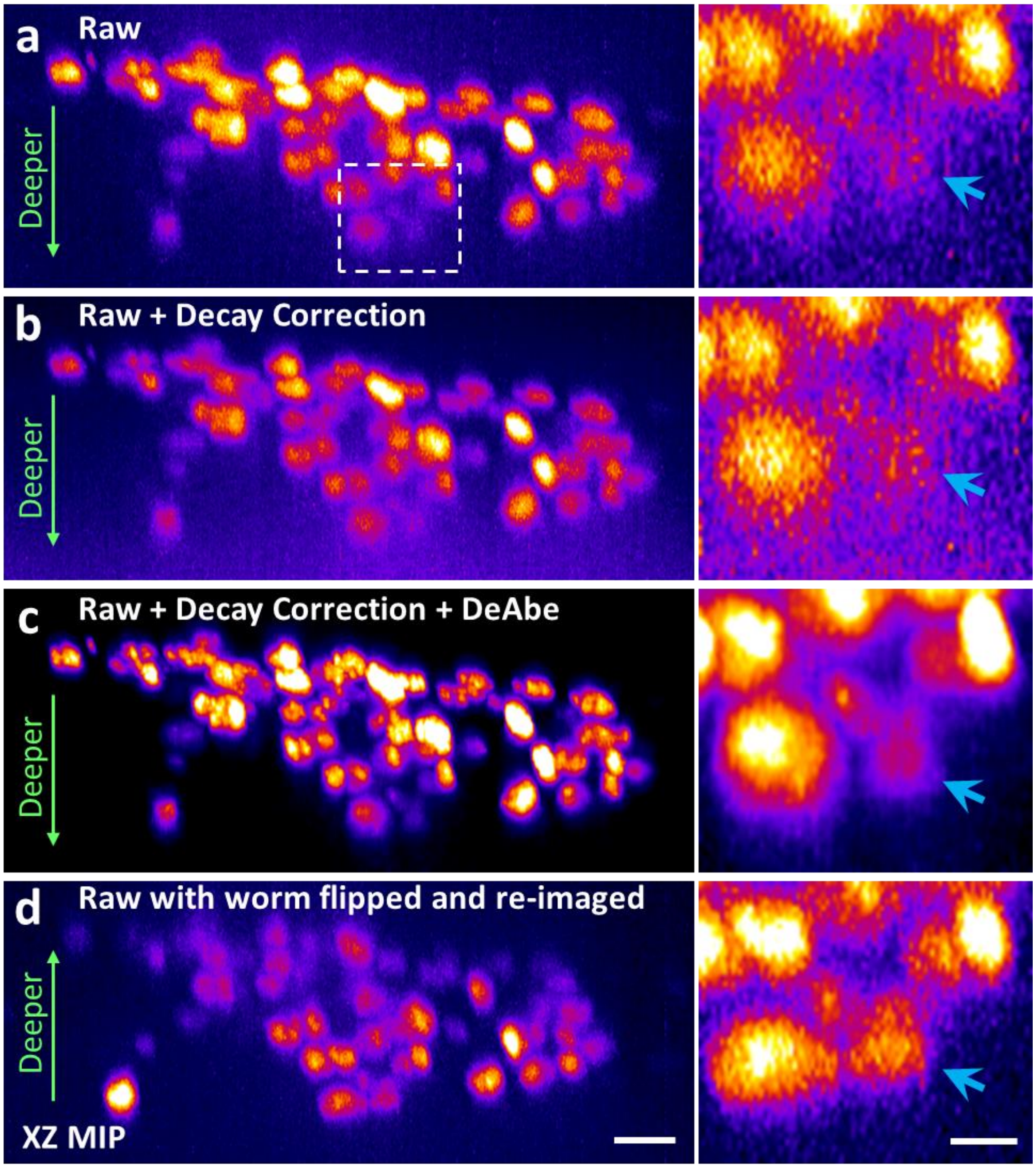


**Supplementary Fig. 20, DeAbe recovers nuclei on the 'far side' of a worm, as verified by 'flipping' the worm.** Adult nematodes expressing mTagRFP-T-tagged neuronal nuclei were anesthetized, immobilized between two surfaces, and imaged with instant structured illumination microscopy. **a)** Raw image data of worm, showing depth-dependent degradation in image quality. **b)** Raw data after applying an exponential weighting to account for depth-dependent loss in intensity. **c)** As in **b)**, but after additionally applying DeAbe. **d)** As in **a)**, but 'flipping' the worm so that depth-dependent degradation occurs in the opposite direction. Higher magnification views at right correspond to white rectangular region in **a)**; arrows highlight nuclei for visual inspection. Axial maximum intensity projections (XZ MIP) are shown, the 'fire' look up table in ImageJ was used to highlight large dynamic range within images. Scale bars: 5 µm and 2 µm for insets. Data shown are representative samples from N = 3 experiments.

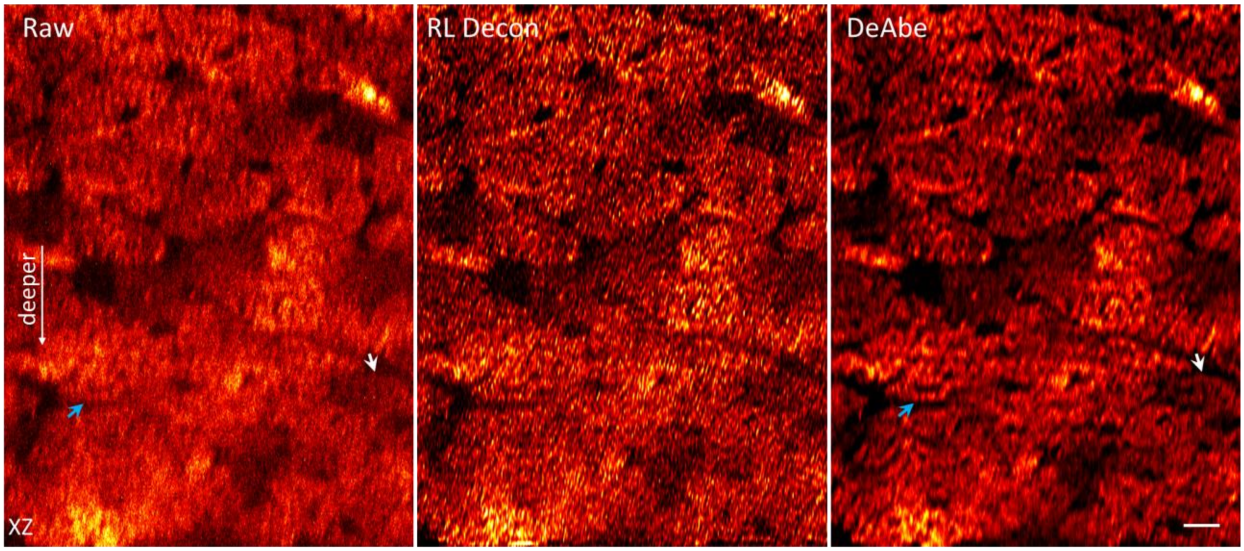


**Supplementary Fig. 21, DeAbe compensates for aberrations in two photon microscopy.** Live cardiac tissue containing cardiomyocytes expressing Tomm20-GFP imaged with two photon microscopy. Left: axial view of raw data with the direction of increasing depth indicated with white arrow. Middle: Richardson-Lucy deconvolution on the raw volume (20 iterations). Right: DeAbe model prediction. Arrows highlight space between cells or cell boundaries better delineated in prediction than raw data. See also **Supplementary Video 6**. Scale bar: 10 μm. Data shown are representative samples from N = 3 experiments.

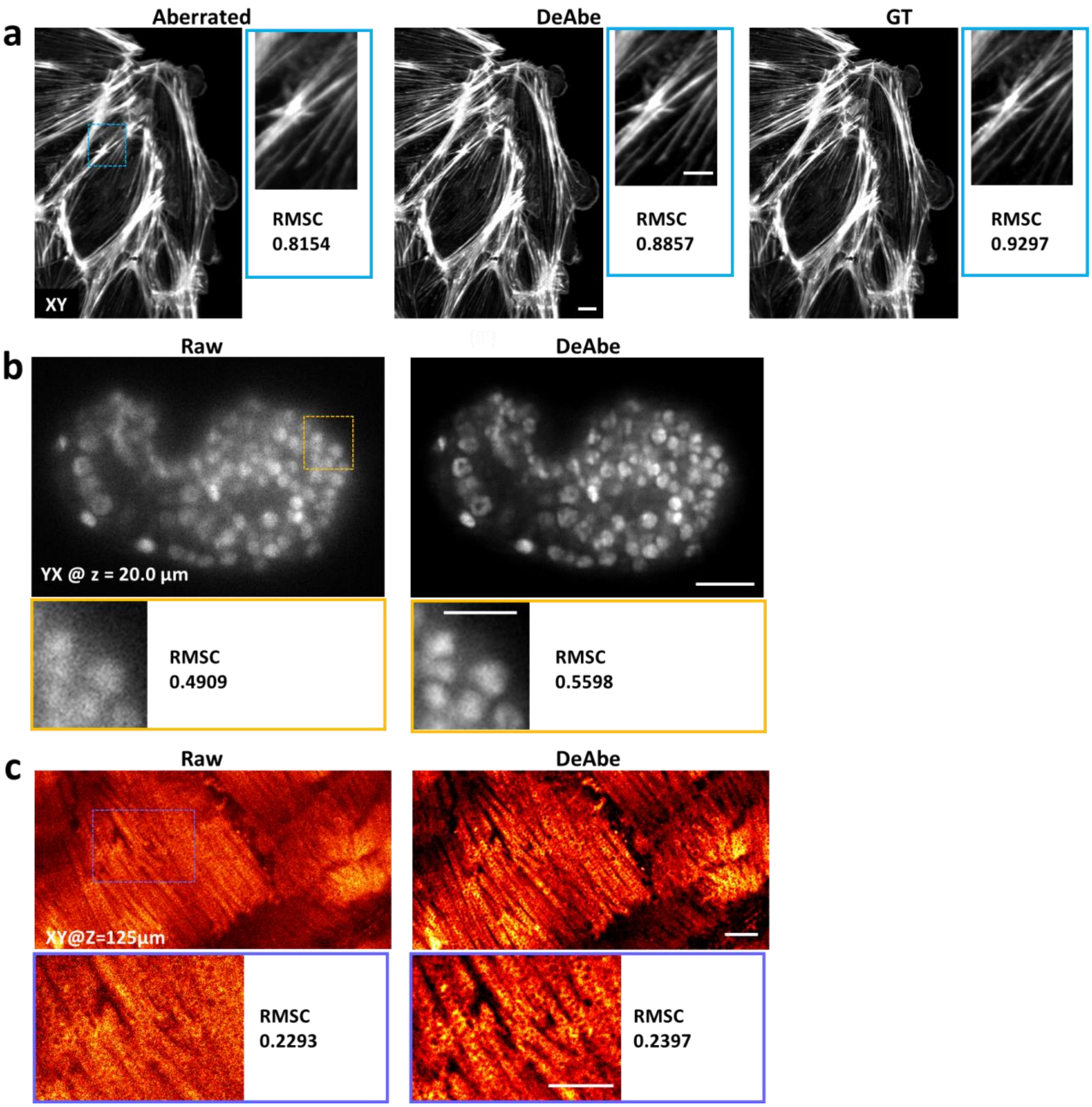


**Supplementary Fig. 22, Root mean square contrast enhancement (RMSC) provided by DeAbe.** The RMSC (see also **Methods**) is calculated based on the ROIs indicated by the rectangles in each subfigure **a-c)**, corresponding to **Figs. 2a,d, 3a, and 3e.** Higher values of RMSC are better. Scale bars: 10 µm for lower magnification views and 5 µm for insets. Data shown are representative samples from N = 3 experiments.

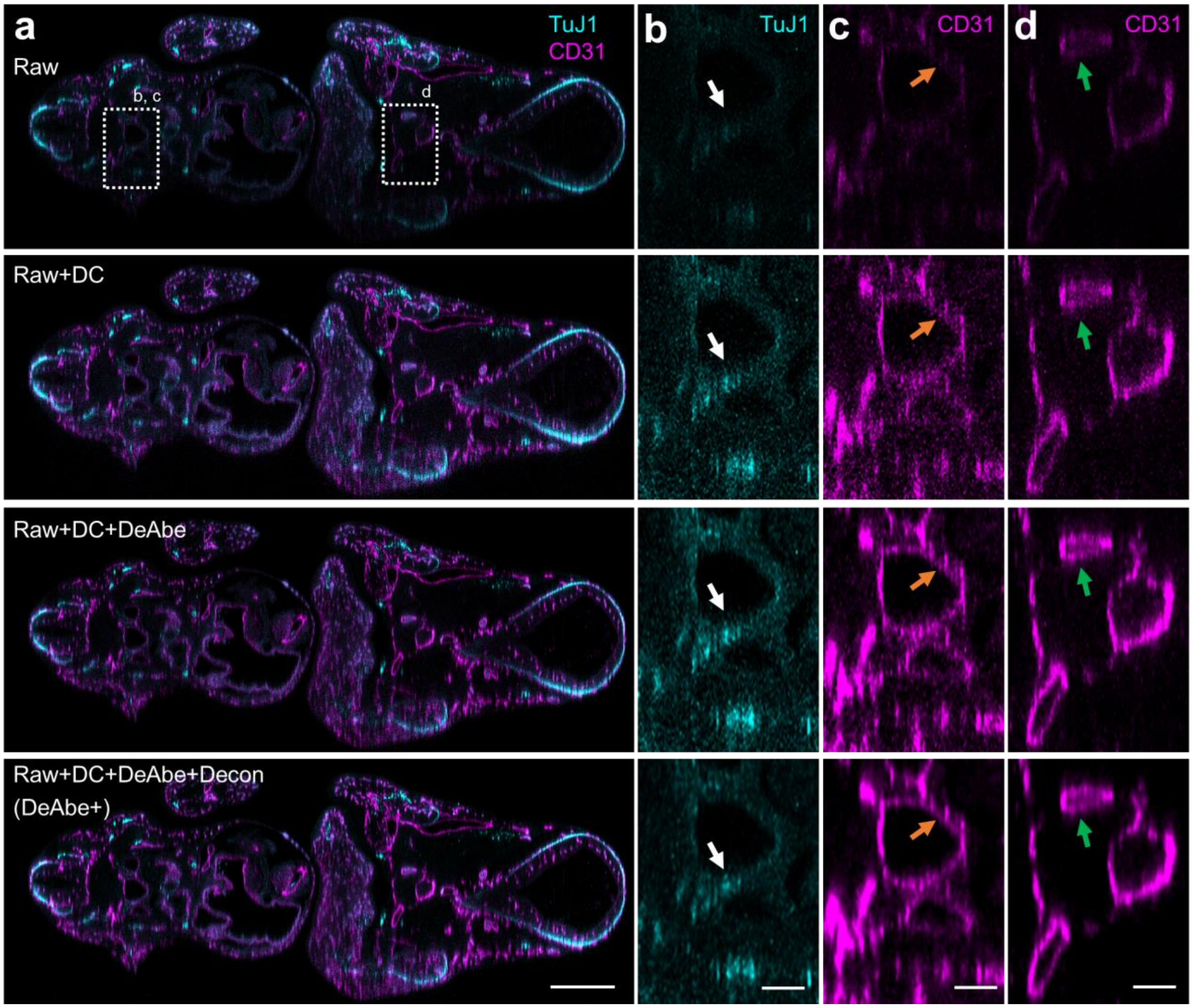


**Supplementary Fig. 23, Progressive improvements in image quality in cleared mouse embryo images.** **a)** Axial views of cleared mouse embryo corresponding to data in **Fig. 4b**, comparing (top to bottom) raw data; intensity decay compensated (DC, see also **Methods**); DC followed by application of DeAbe; and DC, DeAbe and deconvolution ('DeAbe+', 20 iterations Richardson-Lucy). Higher magnification views of dashed rectangular regions in **a)** are shown in TuJ1 channel (**b**) and CD31 channel (**c, d**). Arrows highlight boundaries that are progressively more clearly delineated after each restoration step. See also **Supplementary Video 8**. Scale bars: 500 μm **a)**; 100 μm **b, c, d)**. Data shown are representative samples from N = 3 experiments.

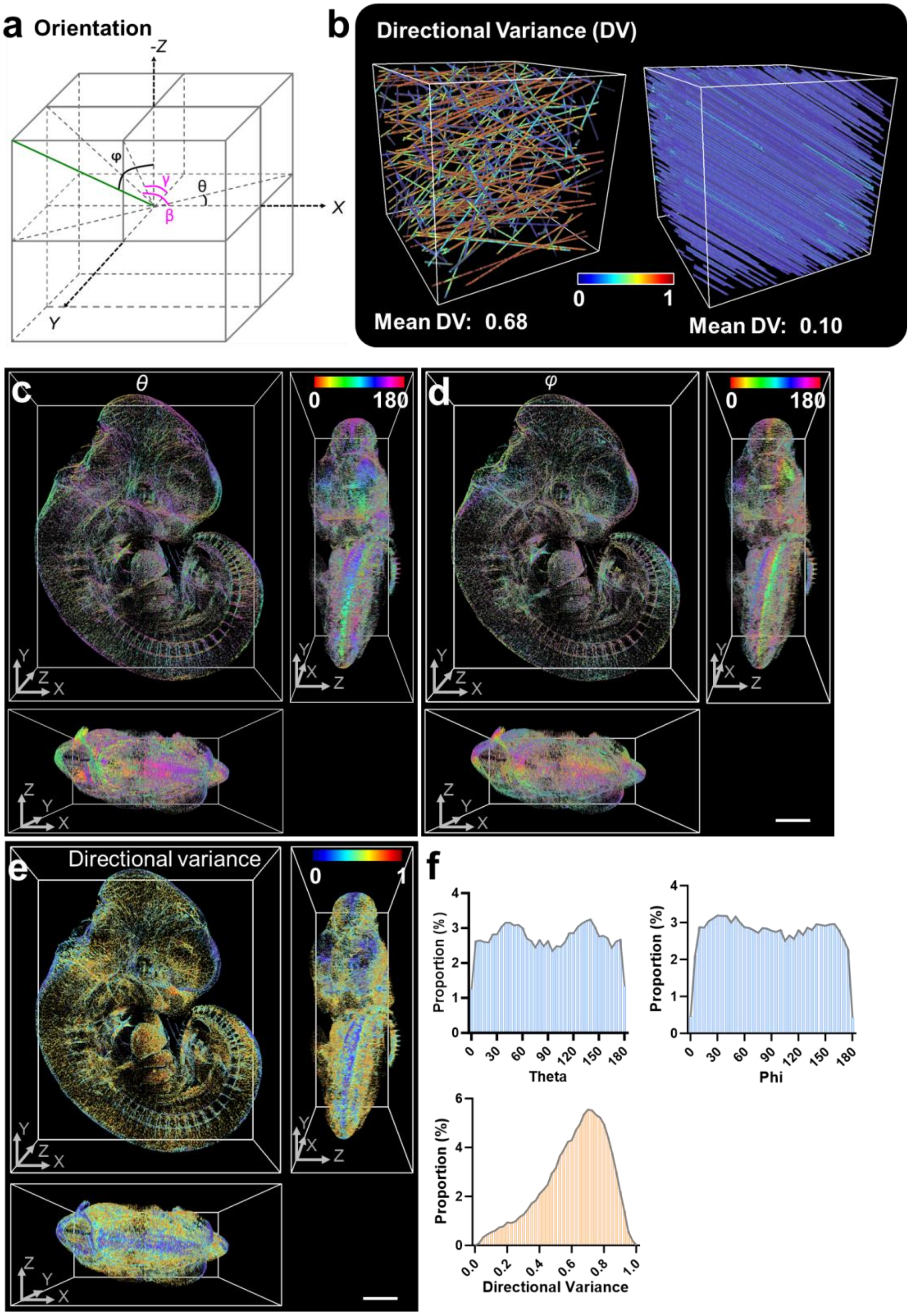


**Supplementary Fig. 24**, **Key variables in orientational vessel analysis. a)** Defining the azimuthal angle $\theta$ and polar angle $\varphi$ variables for a certain orientation (the green line) with respect to Cartesian coordinate system, with two additional two azimuthal angles, $\beta$ and $\gamma$, to assist the calculation of $\varphi$. **b)**

simulation of oriented lines with more (left) or less (right) mean directional variance (DV). More directional variance implies lines are more randomly oriented. **c-e)** Orientation analysis on blood vessel channel of DeAbe processed data of CLARITY-cleared E11.5-day mouse embryo, immunostained for neurons (TuJ1) and blood vessels (CD31).  Perspective views of $\theta$ (**c**), $\varphi$ (**d**) and DV (**e**) distributions in 3D are shown. **f**) Histograms of $\theta$ (top left), $\varphi$ (top right), and DV (bottom). See also **Fig. 4e-g, Supplementary Video 9.** Scale bars: 500 $\mu$m. Data shown are representative samples from N = 3 experiments.

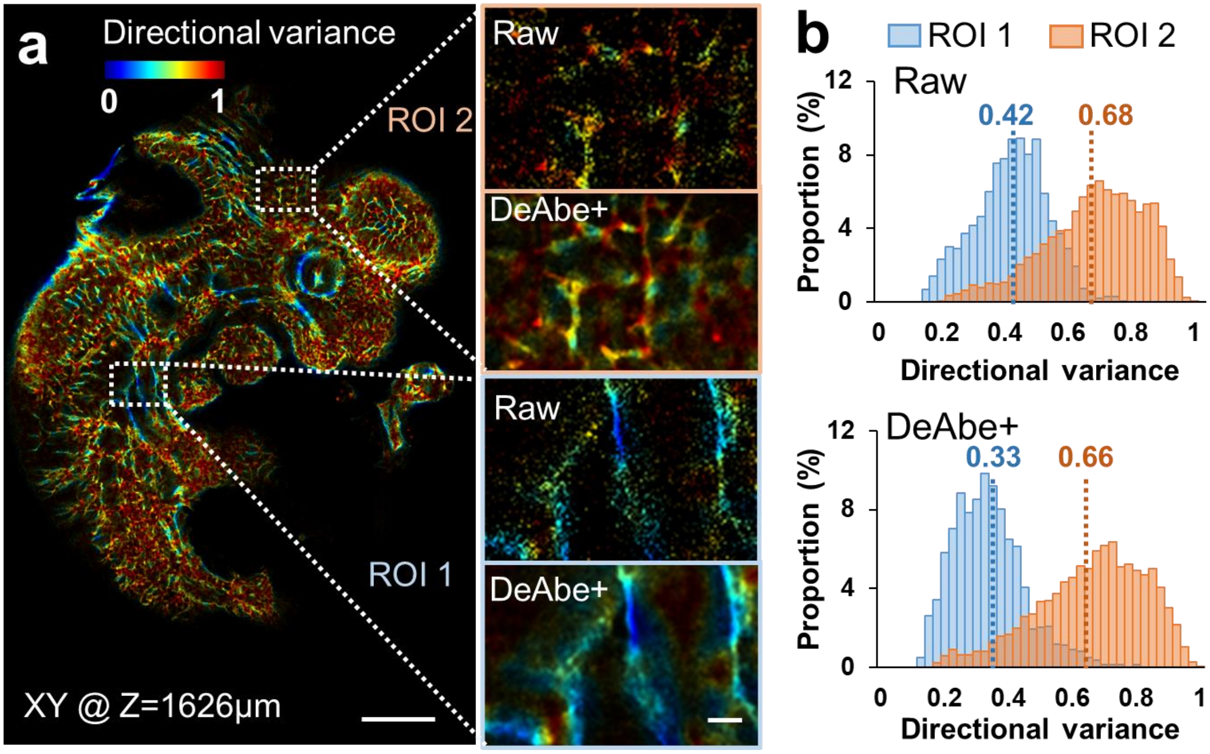


**Supplementary Fig. 25, Comparing directional variance between raw and DeAbe+ restoration. a)** Data are as in **Fig. 4g**, but higher magnification views of each ROI also show directional variance computed from raw data in addition to DeAbe+. Note noisier appearance in raw data, which results in less separation between histograms (**b**). Scale bars: 300 μm **a)**, 50 μm inset. Data shown are representative samples from N = 3 experiments.

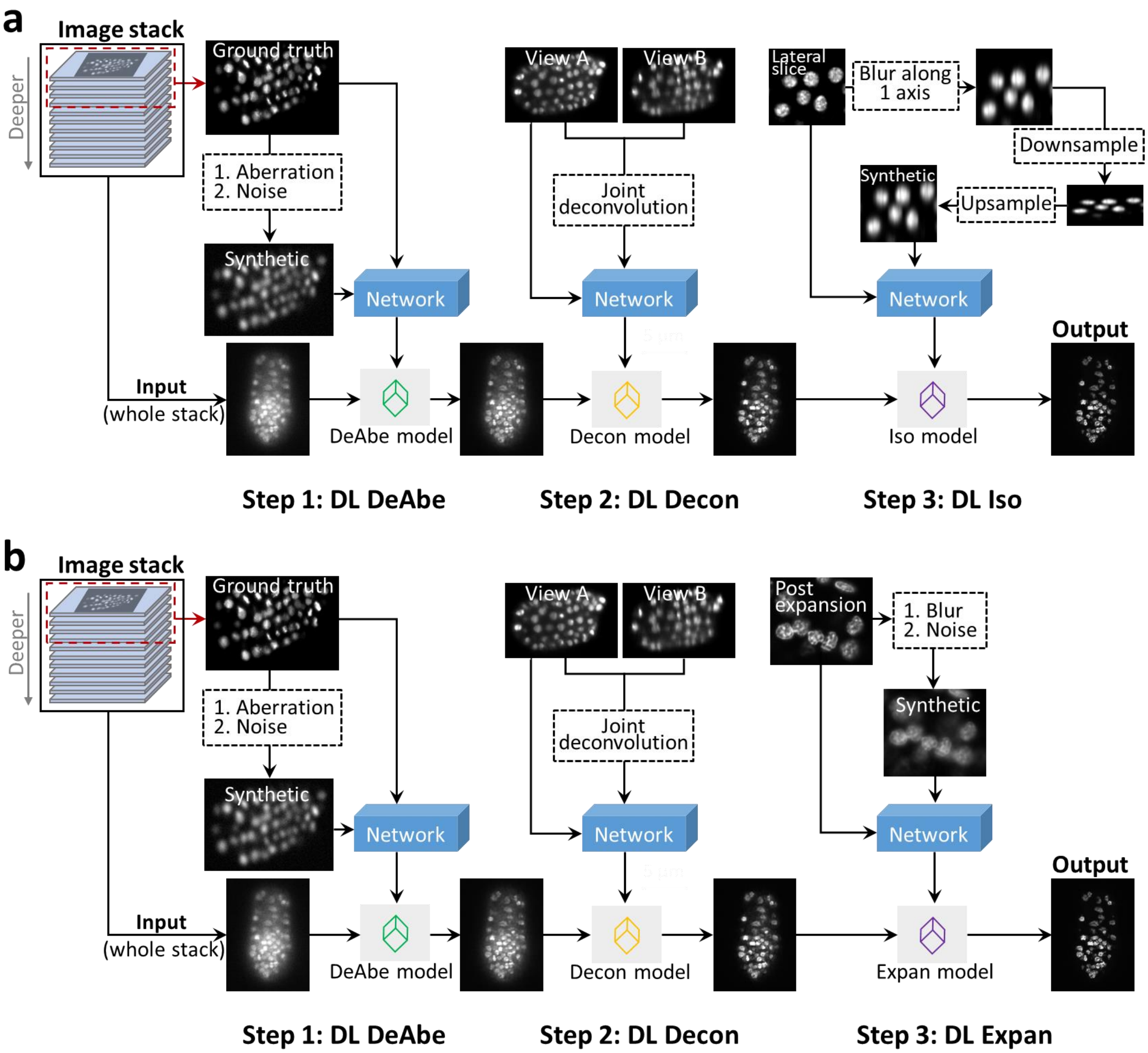


**Supplementary Fig. 26, Multi-step image restoration schemata.** We trained independent neural networks to compensate for aberrations (Step 1: DL DeAbe, see also **Fig. 1a**), deconvolve the data (Step 2: DL Decon), and to super-resolve the data (Step 3: DL Iso, **a)** or DL Expan, **b)**). For the deconvolution networks, we used high quality joint deconvolution based on orthogonal views (after DL DeAbe processing) from diSPIM as ground truth and trained the network to predict this joint deconvolution given only single-view input. For the isotropic enhancement model in **a)**, we synthetically blur lateral views to resemble axial views and downsample and upsample the data to mimic the coarser pixel size in z. We then train a neural network to reverse this degradation. For the expansion microscopy network in **b)**, we synthetically degraded high-resolution images from expanded samples until the synthetic data resembled conventional images acquired on the diSPIM and trained a network to reverse this degradation. Serial application of each network produced the final prediction. See also **Methods**.

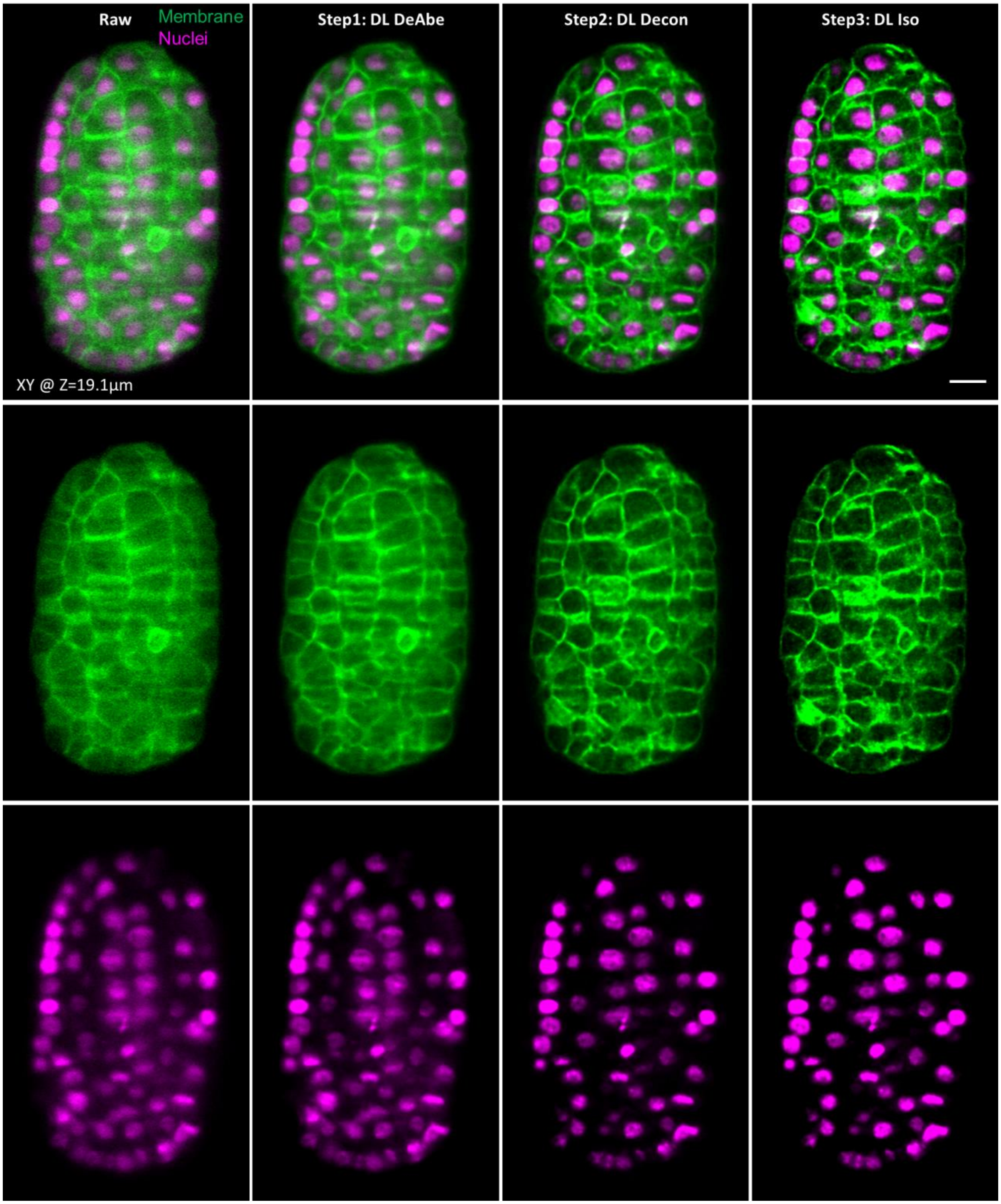


**Supplementary Fig. 27, Improvements in lateral views after multi-step deep learning.** Data from *C. elegans* embryos expressing membrane and nuclear markers as in **Fig. 5a**, emphasizing improvements to lateral views (single plane 19.1 μm into the volume) corresponding to successive steps in restoration (columns). Merged views (top), membrane channel (middle), and nuclear channel (bottom) are shown. Scale bar: 5 μm. Data shown are representative samples from N = 3 experiments.

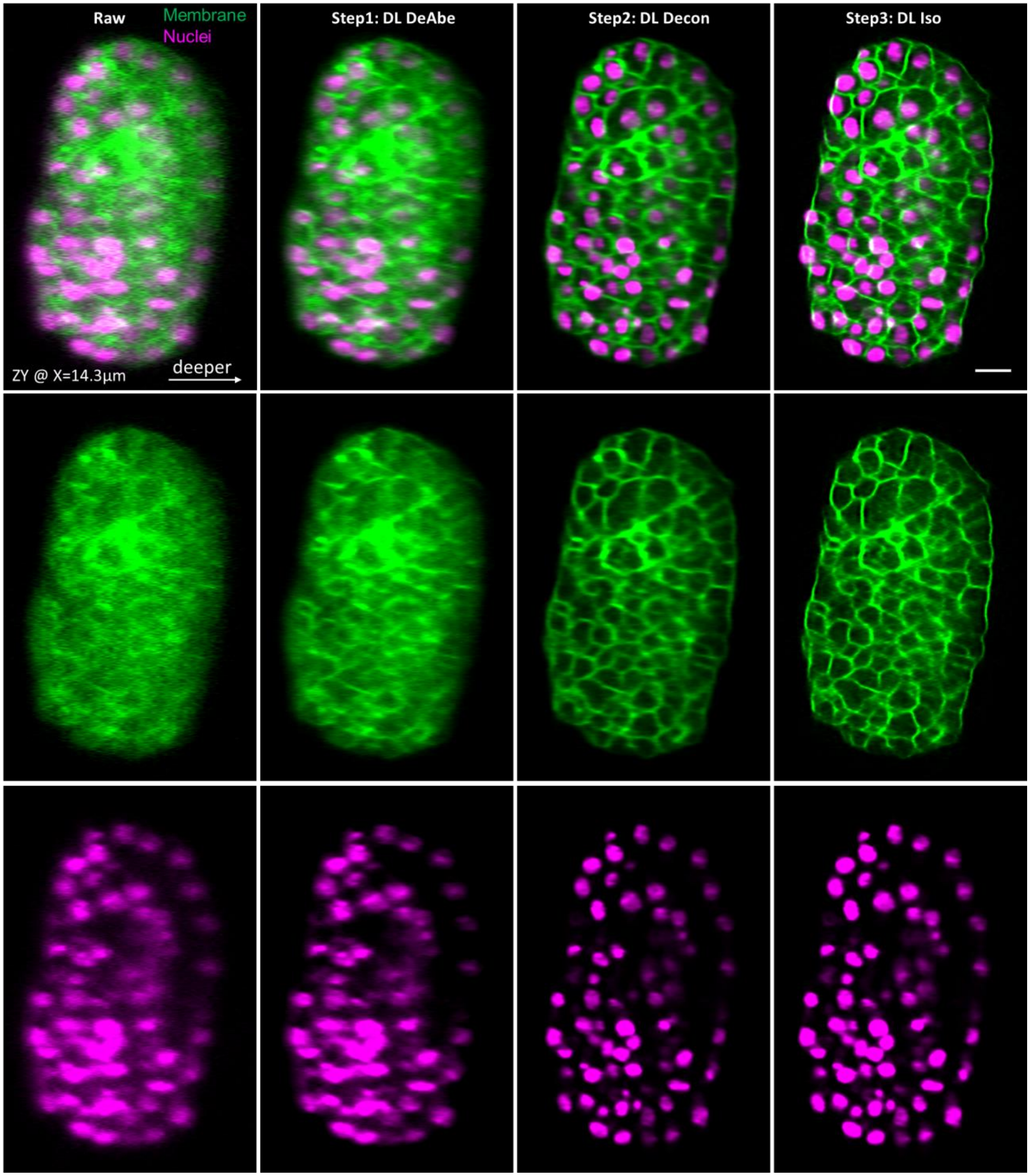


**Supplementary Fig. 28, Improvements in axial views after multi-step deep learning.** Data from *C. elegans* embryos expressing membrane and nuclear markers as **Fig. 5a**, emphasizing improvements to axial views (single plane 14.3 μm into the volume) corresponding to successive steps in restoration (columns). Merged views (top), membrane channel (middle), and nuclear channel (bottom) are shown. Scale bar: 5 μm. Data shown are representative samples from N = 3 experiments.

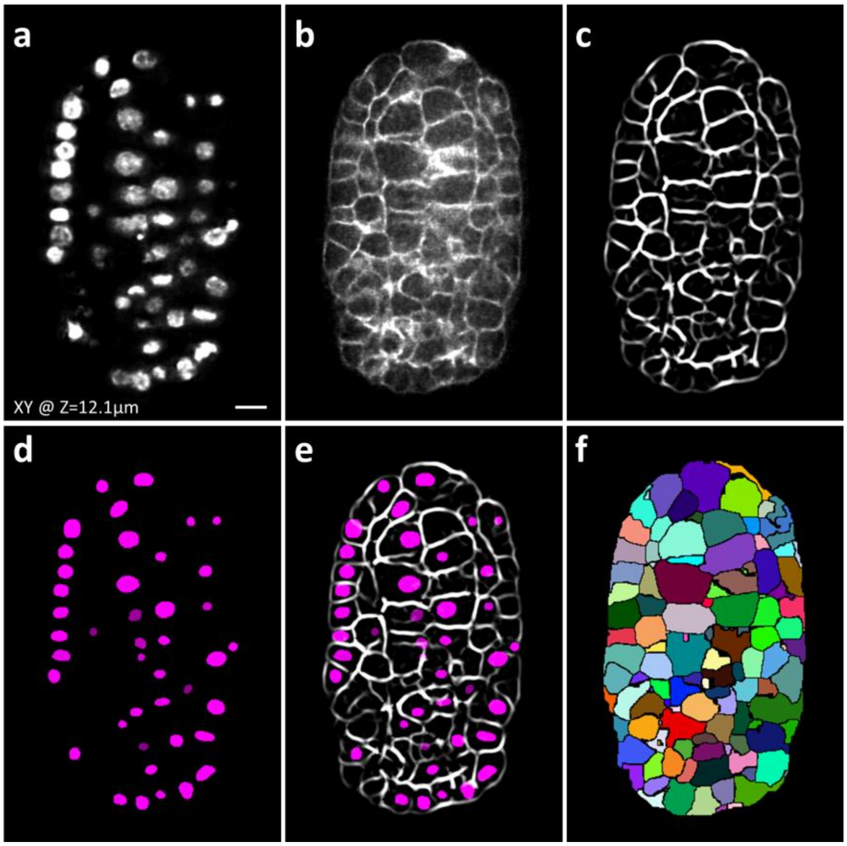


**Supplementary Fig. 29, Automated membrane segmentations on restored image volumes derived from dual-color (pan-nuclear and membrane) *C. elegans* embryos. a, b)** Image planes from nuclear and membrane channels from representative volume as in **Fig. 5a-d**, after multistep image restoration. **c)** Membrane-enhanced image after applying vascular structure enhancement filter to **b**. **d)** Segmented nuclei with Mask RCNN. **e)** Overlay of enhanced membrane and segmented nuclei. **f)** The associated multi-instance segmentation of cell boundaries. Scale bar: 5 μm. Data shown are representative samples from N = 3 experiments.

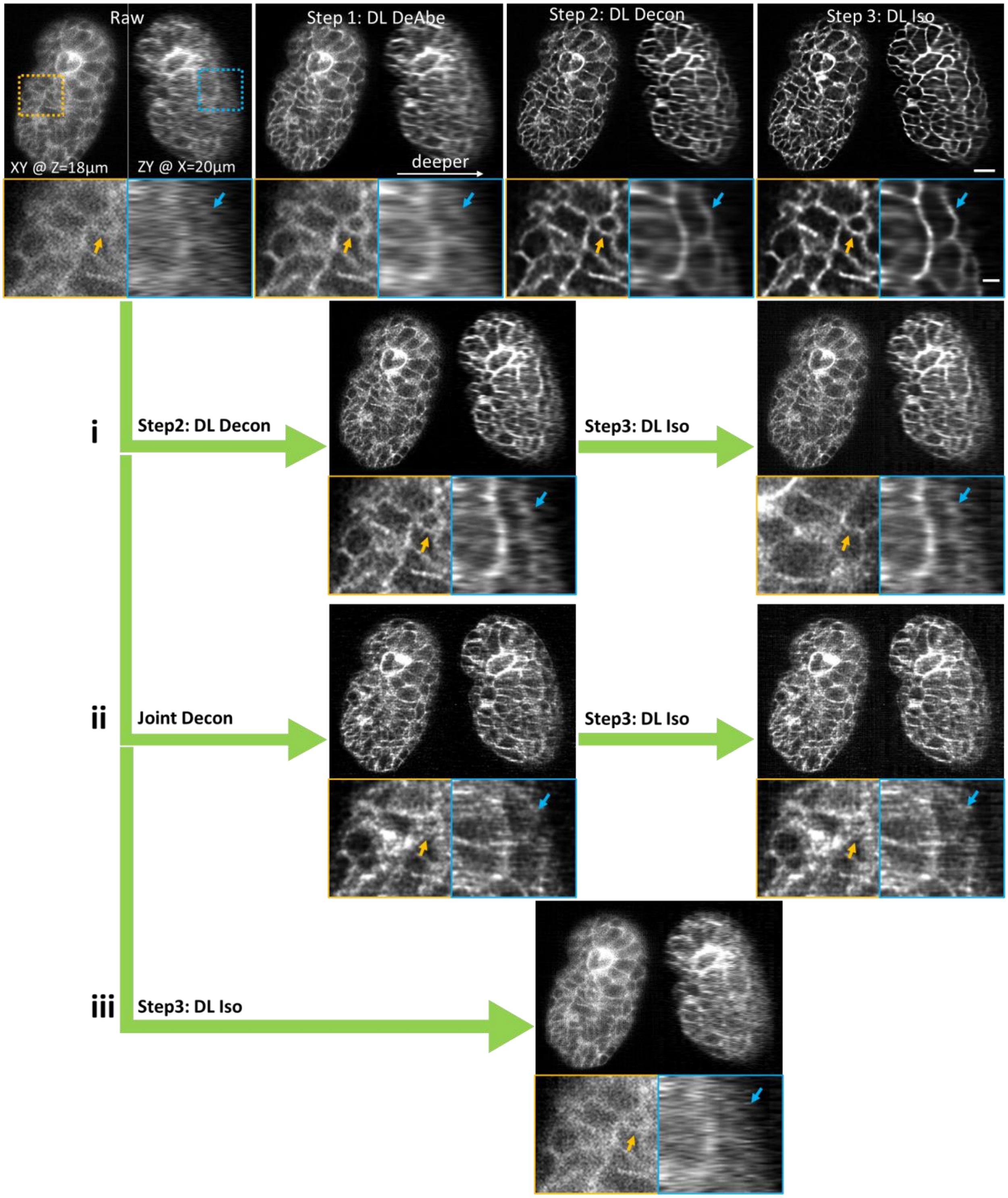


**Supplementary Fig. 30, Ablation experiments highlighting value of DeAbe model in multi-step restoration.** Top rows show data from *C. elegans* embryos expressing membrane marker, highlighting progressive improvements in image quality after each step of multi-step restoration, including higher magnification insets in second row corresponding to dashed rectangular regions in top row. Data are also shown (i) if DeAbe (Step 1) is ablated, proceeding directly from raw data to Steps 2 and 3; (ii) if

DeAbe is ablated, and joint deconvolution used in lieu of the DL Decon (Step 2); (iii) if steps 1 and 2 are both ablated, and raw data is used directly as in the input for the DL Iso network (Step 3). The final prediction is degraded in schemes (i) – (iii) relative to the original method, as membranes are either noisy or not well resolved (arrows). Scale bars: 5 μm, 2 μm inset. Data shown are representative samples from N = 3 experiments.

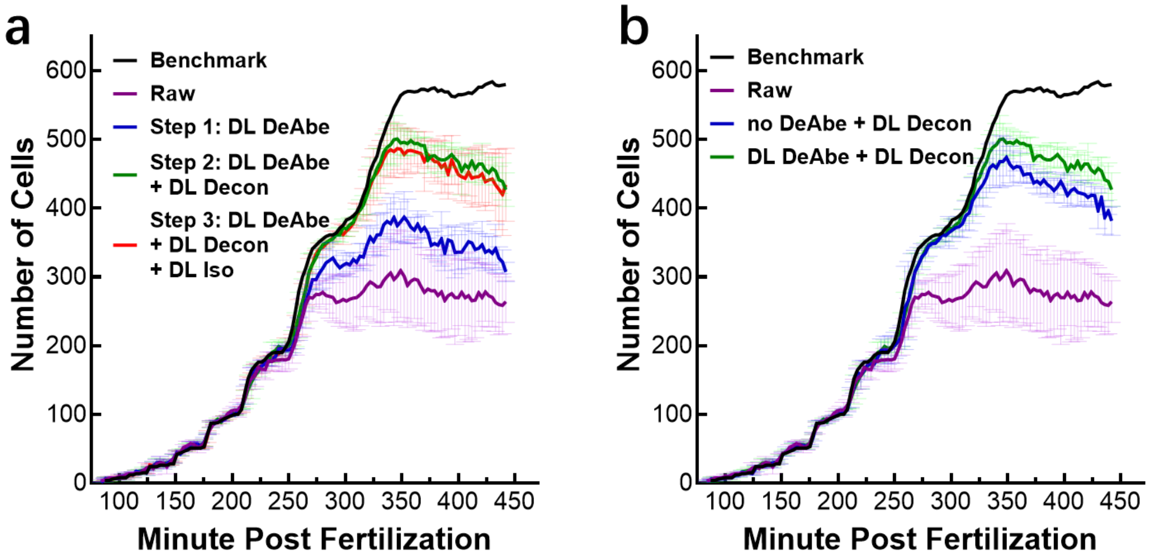


**Supplementary Fig. 31, Evaluating the contribution of different networks to cell segmentation.** Analysis corresponds to cell segmentation shown in **Fig. 5a-d). a)** # of cells segmented manually ('Benchmark'), from raw data ('Raw'), using DeAbe prediction (Step 1), using DeAbe + DL Decon (Step 2), or using DeAbe+DL Decon +DL Iso (Step 3). Step 3 offers no improvement relative to Step 2. **b)** Ablation study showing that DeAbe improves cell segmentation (~10% more cells are segmented in green vs. blue curve) compared to using only DL Decon network. Means and standard deviations from three embryos are shown. Statistics are derived from N = 3 different embryos.

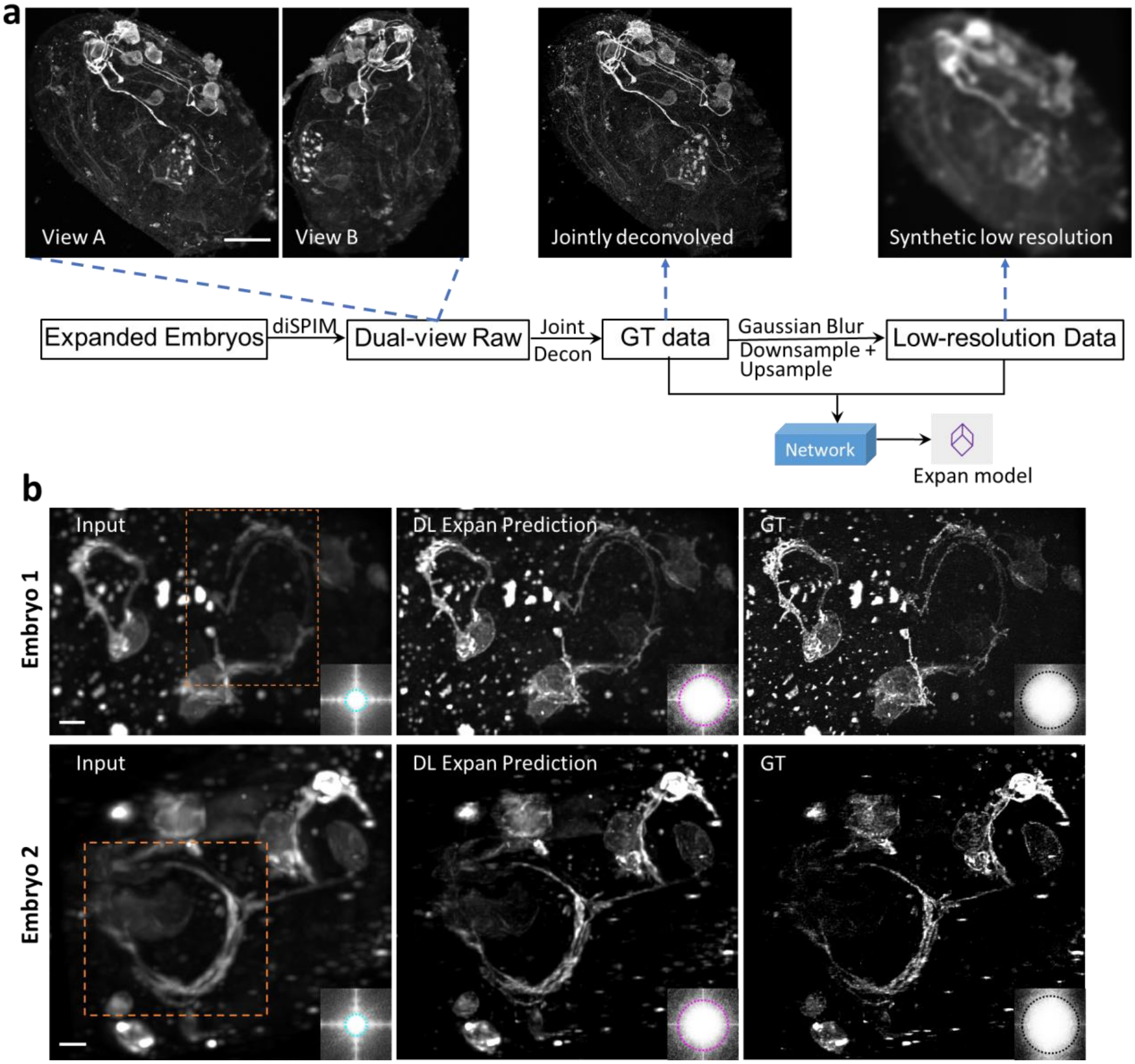


**Supplementary Fig. 32, Using expanded embryos as high resolution ground truth for training resolution enhancement model. a)** Expansion was used to enlarge embryos expressing ttx-3B-GFP 3.6-fold and the samples were then imaged using diSPIM. The two views (A and B) were registered, and joint deconvolution applied to produce volumes with more nearly isotropic resolution. These volumes were synthetically degraded via blurring and downsampling to produce semisynthetic data with resolution more like the raw data prior to expansion, and used to train a network (Expan model) for resolution enhancement. In these images, GFP signal was boosted using immunolabeling. **b)** Example images from two embryos highlighting nerve ring region (orange dashed rectangles), comparing synthetic, low-resolution input data (left), prediction from DL Expan model, and ground truth data. The Fourier transform images are shown in the insets, with cyan, pink, and black dotted circles corresponding to the spatial resolutions of 330 nm, 130 nm, and 105 nm respectively. Scale bars: 2 μm.

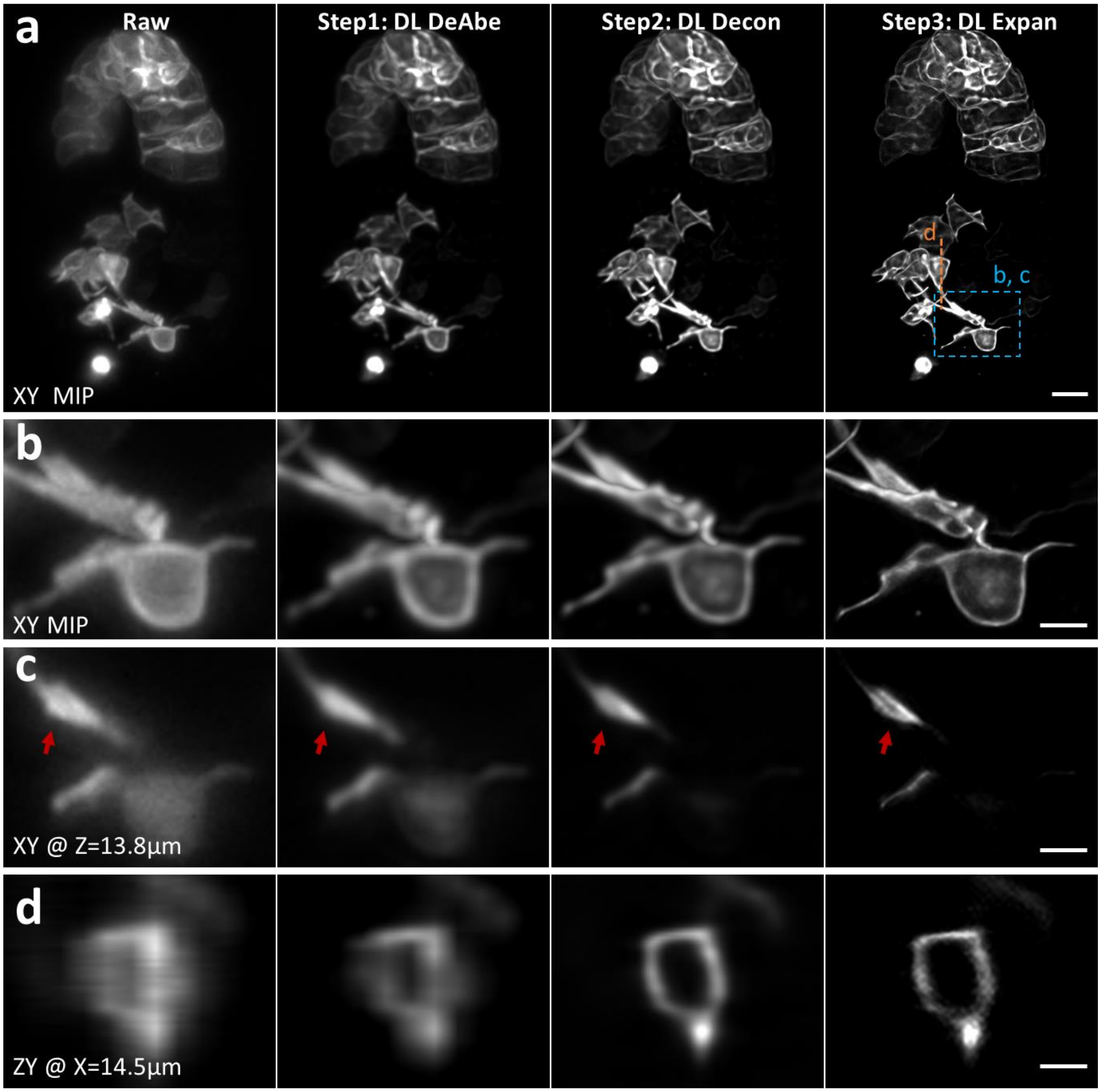


**Supplementary Fig. 33, Progressive improvements in image quality in multi-step image restoration. a)** *C. elegans* embryos expressing membrane markers in neurons and gut cells were imaged as in **Fig. 5e** and passed through our multi-step image restoration pipeline. Higher magnification lateral **b, c)** and axial **d)** views of dashed blue rectangle and orange line in **a)** are also shown. Red arrows highlight features resolved after the third step. MIP: maximum intensity projection. Scale bars: 500 µm **a)**; 100 µm **b, c, d)**. Data shown are representative samples from N = 3 experiments.

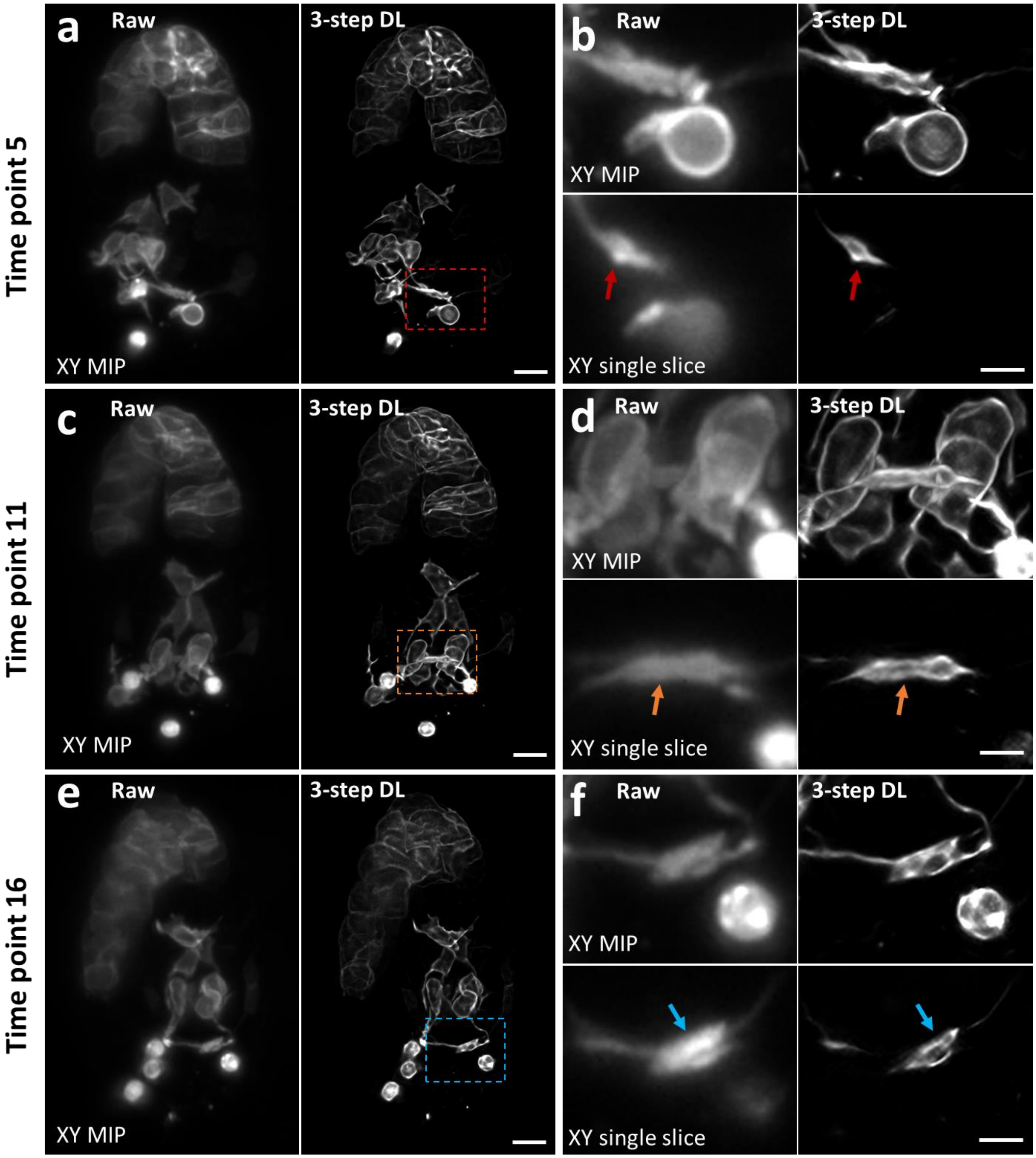


**Supplementary Fig. 34, Additional examples of improved image quality after multi-step image restoration**. **a)** Selected time points from the dataset highlighted in **Fig. 5e-i**, further comparing raw data (left) and multi-step image restoration (3-step DL, right). Higher magnification views of dashed red **b)**, orange **c)**, and blue **d)** rectangles in **a)** are also shown in lateral maximum intensity projections (top) or single planes (bottom). Arrows indicate features that are obscured in raw data but better resolved after image restoration. Scale bars: 500 μm **a, c, e)**; 100 μm **b, d, f)**. Data shown are representative samples from N = 3 experiments.

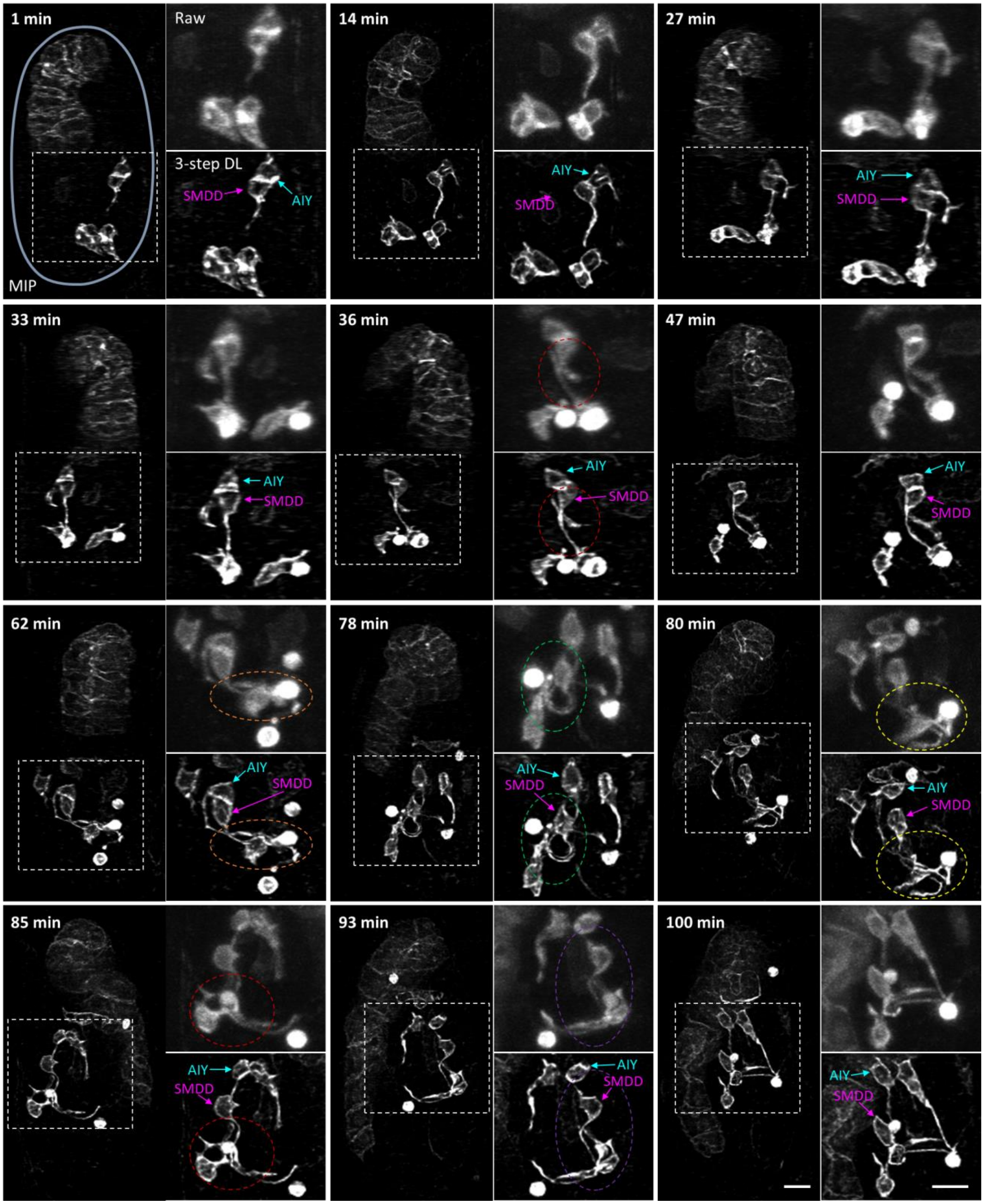


**Supplementary Fig. 35, Visual inspection of AIY and SMDD neurites is facilitated after multi-step image restoration.** Example images in support of **Fig. 5i** at indicated time points, highlighting additional clarity after image restoration. Higher magnification views of dashed white rectangles are shown at right, comparing raw (upper) and restored (lower) data. AIY and SMDD cell bodies have been indicated with arrows for clarity; dashed circles highlight associated neurites as they innervate the nerve ring area. Scale bars: 5 μm. Data shown are representative samples from N = 3 experiments.

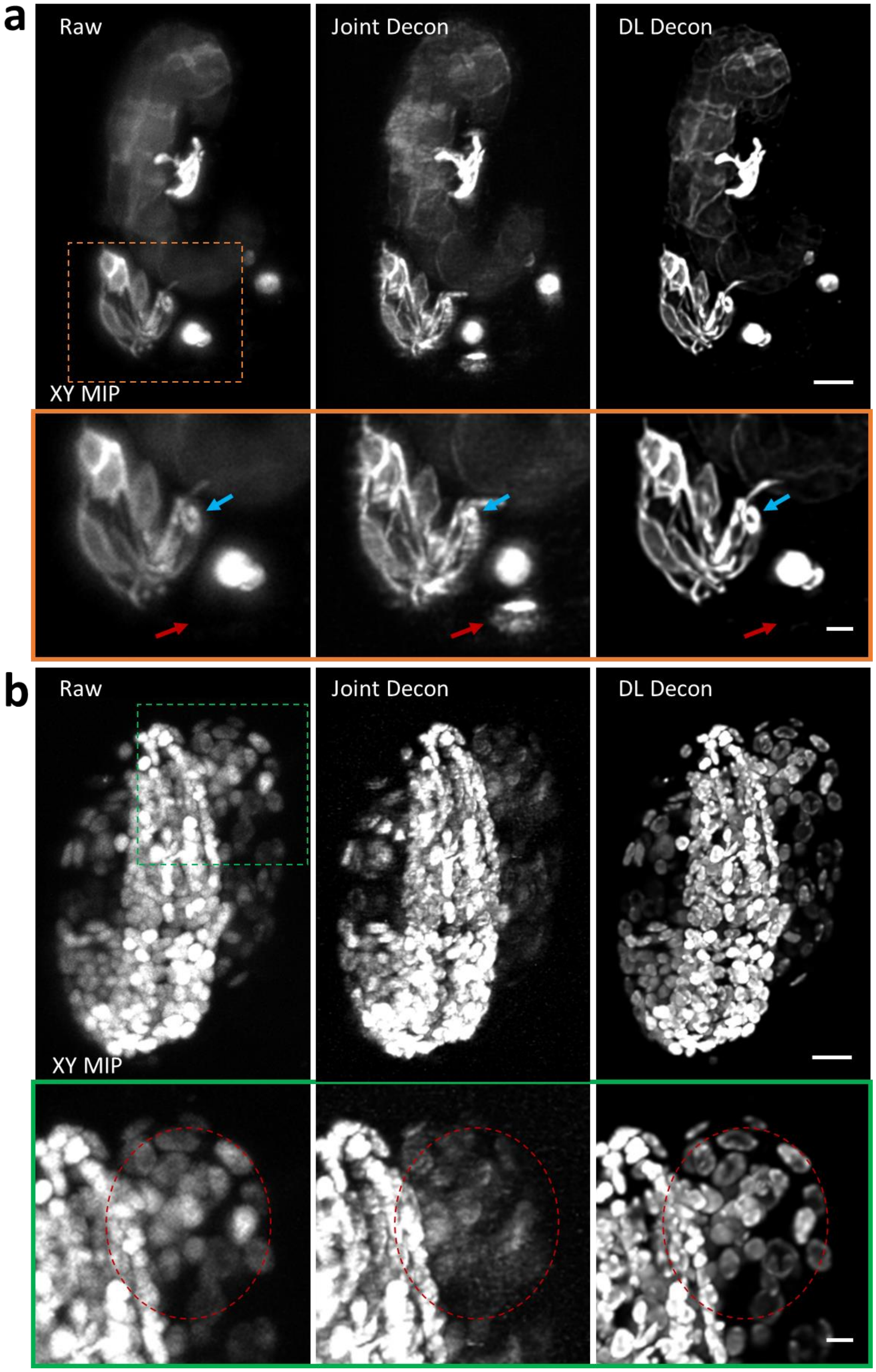


**Supplementary Fig. 36, Multi-step image restoration does not suffer motion artifacts like multiview fusion. a)** Maximum intensity projection (MIP) images of *C. elegans* embryos expressing ttx-3B-GFP, imaged with diSPIM, comparing raw single-view recordings (Raw, left), dual-view fusion result via traditional joint deconvolution (Joint Decon, middle) and deep learning-based deconvolution after application of DeAbe (DL Decon, right). Higher magnification lateral views of orange dotted rectangular region are shown in the bottom panel, with cyan and red arrows highlighting motion induced artifacts present in Joint Decon images but not Raw or DL Decon images. **b**) Images of *C. elegans* embryos expressing pan-nuclear GFP marker, imaged with symmetric diSPIM. Higher magnification lateral views of green dotted rectangular region are shown in the bottom panel, with red dotted circles highlighting the motion induced artifacts present in Joint Decon images but not Raw or DL Decon images. Scale bars: 5 μm, 2 μm insets. Data shown are representative samples from N = 3 experiments.

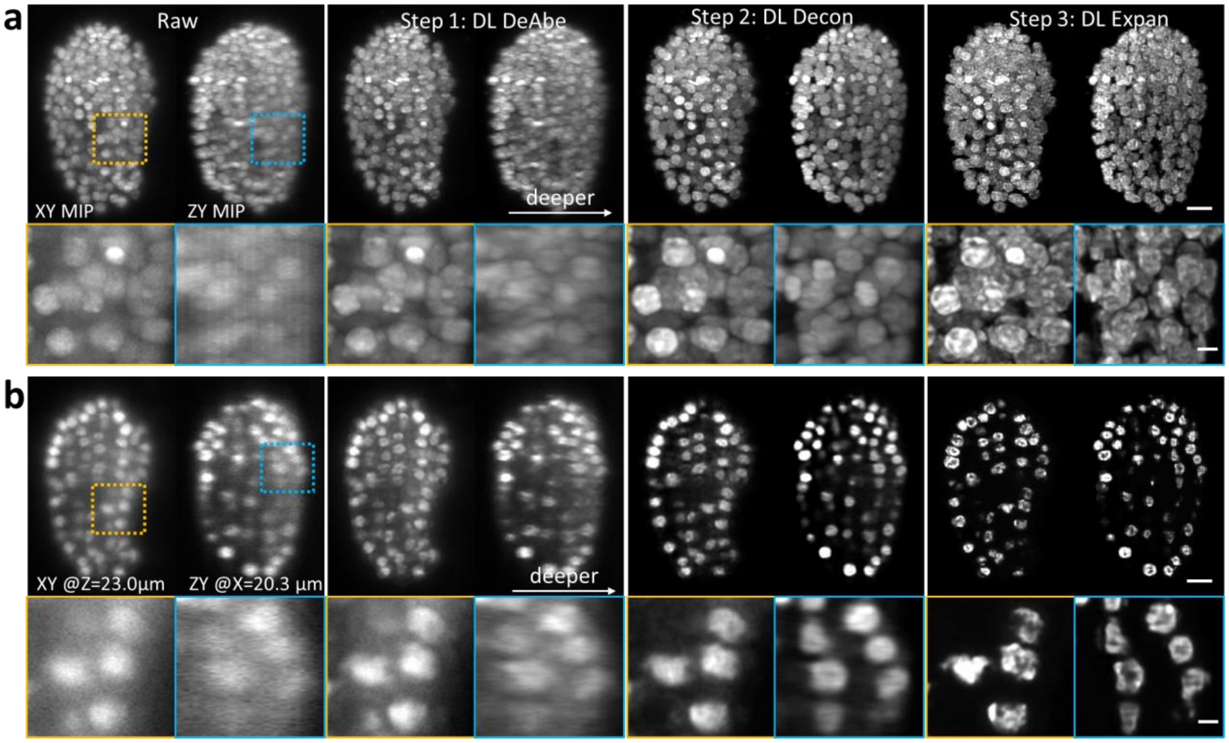


**Supplementary Fig. 37, Additional example of multi-step restoration. a)** *C. elegans* embryos expressing pan-nuclear GFP marker were imaged with high numerical aperture diSPIM and input into our multi-step imaging restoration pipeline. Top row: Comparing raw, after DeAbe (Step 1), after additionally applying deconvolution network (Step 2), and finally after additionally applying DL Expan model (Step 3). See also **Methods** for further details on sample preparation and imaging system. Higher magnification views of lateral (left, corresponding to dashed orange rectangle in top row) and axial (right, corresponding to dashed blue rectangle in top row) maximum intensity projections (MIP) are also shown. **b)** As in **a)**, but highlighting single lateral and axial imaging planes instead of MIPs. Scale bars: 5 μm, 2 μm insets. See also **Supplementary Videos 13, 14.** Data shown are representative samples from N = 3 experiments.

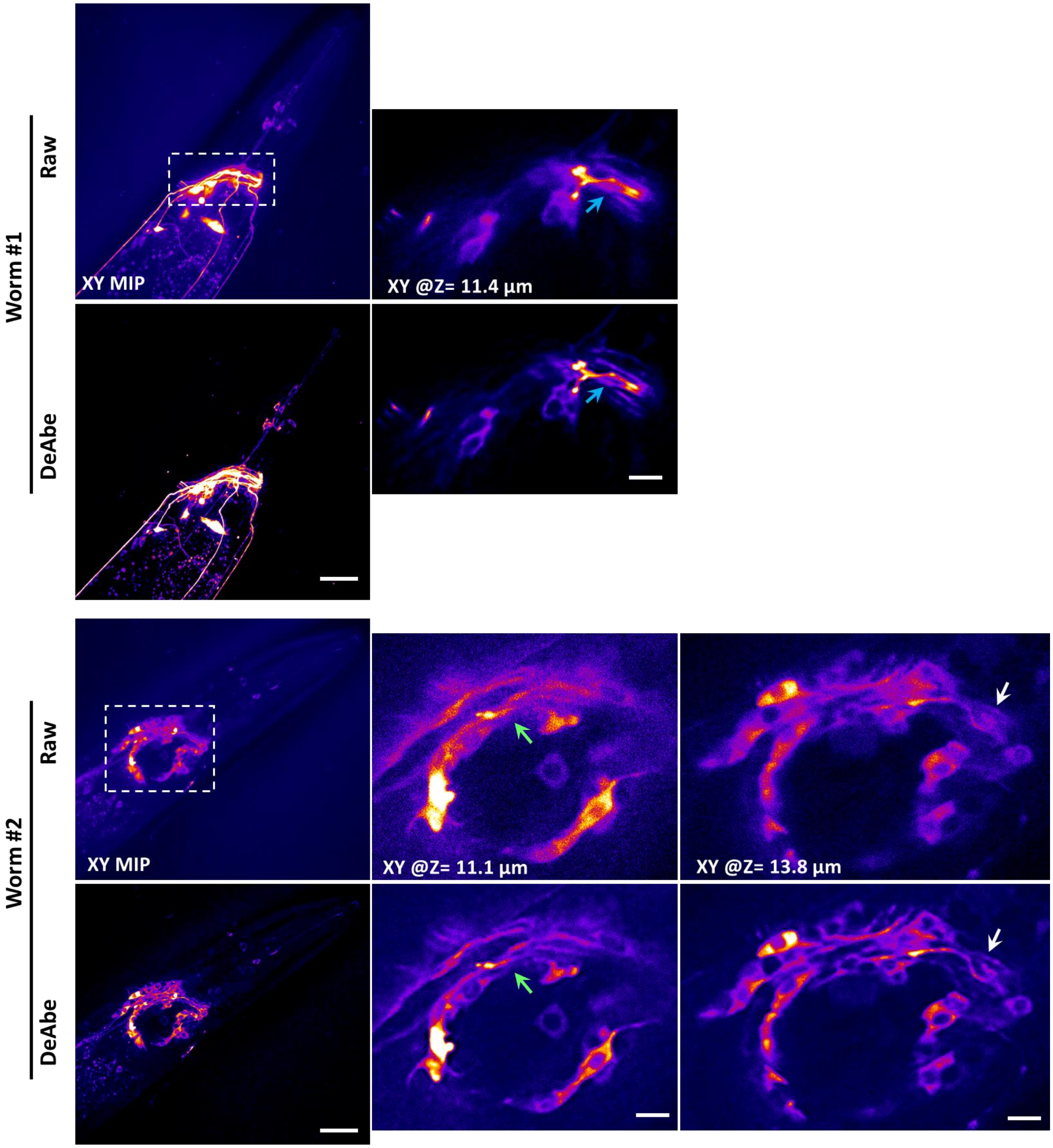


**Supplementary Fig. 38, Demonstration of DeAbe on iSIM images of anesthetized adult worms expressing a GCaMP marker targeted to neurons (pceh-24 promoter).** Left: lower magnification maximum intensity projection views of two worms show the improved image quality provided by DeAbe. Right insets: higher magnification views of single planes at different depths corresponding to the white dashed rectangular regions in left figures, with the arrows highlighting structural details that are better resolved by DeAbe vs. raw data. Scale bars: 20 µm in main figures and 5 µm in higher magnification views. Data shown are representative samples from N = 3 experiments.

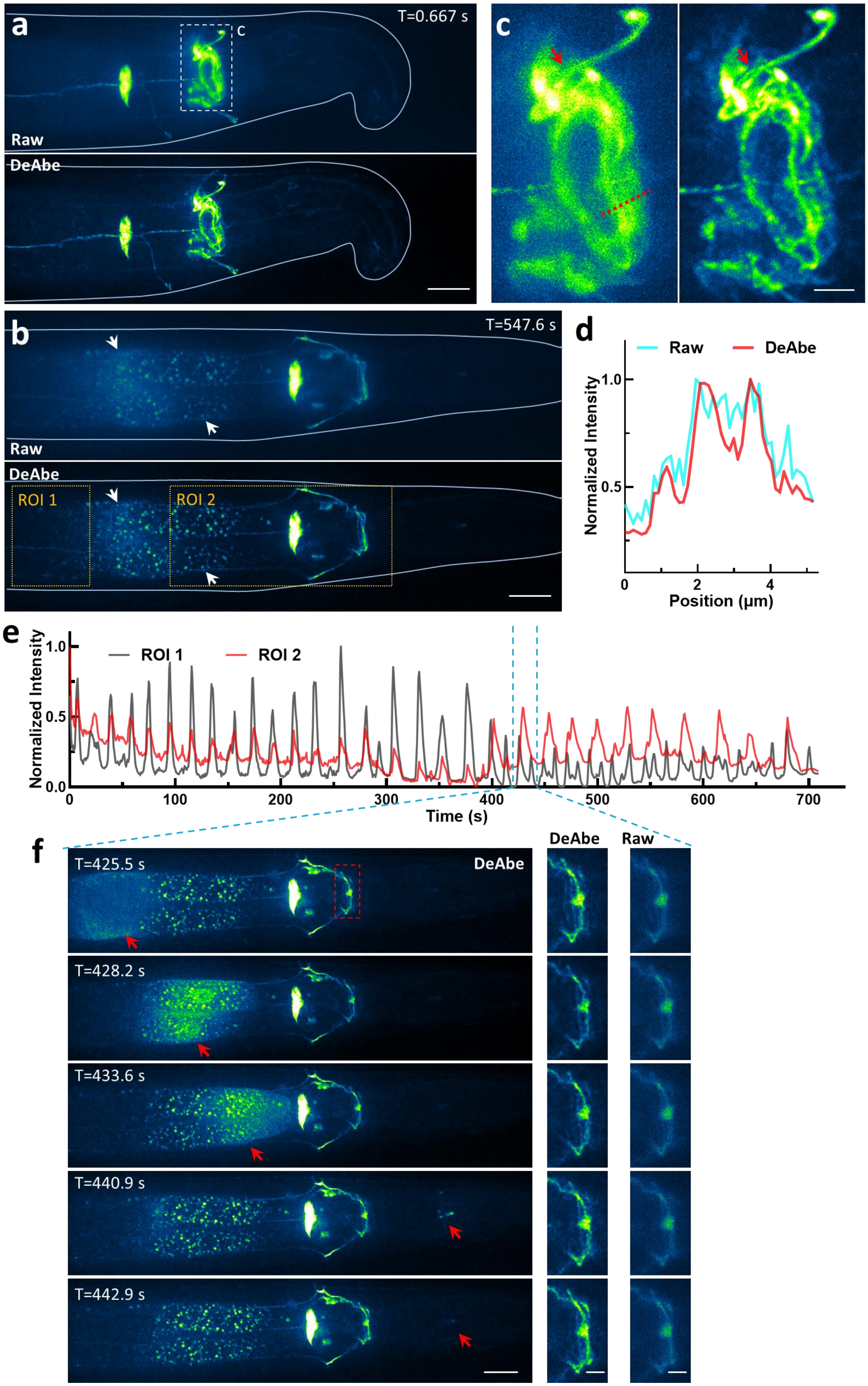

a
T=0.667 s
c
Raw
DeAbe
c
b
T=547.6 s
Raw
DeAbe
ROI 1
ROI 2
d
Raw
DeAbe
Normalized Intensity
1.0
0.5
0
2
4
Position (μm)
e
ROI 1
ROI 2
Normalized Intensity
1.0
0.5
0.0
0
100
200
300
400
500
600
700
Time (s)
f
DeAbe
Raw
T=425.5 s
DeAbe
T=428.2 s
T=433.6 s
T=440.9 s
T=442.9 s

**Supplementary Fig. 39, Demonstration of DeAbe on time-lapse iSIM images of a live worm expressing a GCaMP marker targeted to neurons (pceh-24 promoter), acquired at 1.5 volumes/s. a), b)** comparison of the raw and DeAbe images 0.667 s **(a)** and 547.6 s **(b)** after acquisition commences, with white arrows in **b)** indicating features resolved in the DeAbe prediction but obscured in the raw data. **c)** higher magnification view of the white dashed rectangular region in **a)**, with the red arrows highlighting neurites that are better resolved by DeAbe. **d)** line profiles corresponding to the red dashed line in **c)**. **e)** Calcium signal traces over time for regions of interest (ROIs, orange dashed rectangular regions in **b)** by averaging intensity in the ROIs and normalizing to the range of 0-1. Note distinct calcium signals comparing nerve ring region (ROI 2) compared to gut (ROI 1). **f)** Selected frames within the dashed blue line area in **e)**, showing the transient calcium wave propagation from posterior to anterior of the worm (brighter intensity indicated by the red arrows) and clearer delineation of signal in nerve ring neurites provided by DeAbe vs raw data (righthand columns). Maximum intensity projection views are shown for all data. Scale bars: **a, b**) 20 µm; **c**) 5 µm; **f**) 20 µm, 5 µm for insets. See also **Supplementary Videos 15-17.** Data shown are representative samples from N = 3 experiments.

**Supplementary Table 1, Sample information and parameters used in generating DeAbe models.**

| **Samples** | **Synthetic phantoms** | **NK-92 cells** | ***C. elegans* embryos** | | | | | **Adult *C. elegans*[3]** |
|---|---|---|---|---|---|---|---|---|
| **Figures/Videos** | Fig 1b, Supp Figs 1-9 & Supp Videos 1,2 | Fig 3b-d, Supp Fig 19 & Supp Video 5 | Fig 3a, Supp Videos 3, 13,14 | Fig 5e-i, Supp Figs 33-35, 36a & Supp Video 12 | Supp Fig 36b | Supp Figs 16, 17, 30 | Fig 5a-d, Supp Figs 27-29, Supp Videos 10,11[3] | Supp Fig 18 & Supp Video 4 |
| **Label** | -- | Wheat germ agglutinin (WGA) | Nuclei | Neurites | Nuclei | Membranes | Membranes, Nuclei | Neurons, 4 color channels |
| **Microscopy** | Light sheet microscopy | iSIM | iSPIM | iSPIM | iSPIM | iSPIM | iSPIM | Spinning disk confocal microscopy |
| **Detection NA** | 1.1 | 1.3 | 1.1 | 0.8 | 0.8 | 0.8 | 1.1 | 1.2 |
| **Maximum depth of raw volumes** | 33 μm | 50 μm | 33 μm | 33 μm | 33 μm | 33 μm | 33 μm | 32 μm |
| **Depth of shallow subvolumes** | 33 μm | ~25 μm | 12 μm | 12 μm | 12 μm | 12 μm | 15 μm | 18 μm |
| **Size of shallow subvolumes** | 256x256x256 voxels x 50 subvolumes | ~542x540x68 voxels x 18 subvolumes | ~324x410x20 voxels x 465 subvolumes | ~256x384x16 voxels x 623 subvolumes | ~248x354x14 voxels x 386 subvolumes | ~290x364x16 voxels x 622 subvolumes | ~256x470x32 voxels x 404 subvolumes | 512x256x12 voxels X 153 subvolumes |
| **Training data size[1] (32 bit/voxel)** | 62 GB | 27 GB | 88 GB | 73.4 GB | 28 GB | 87 GB | 145 GB | 18 GB |
| **Training time cost** | 27 h | 18 h | 26 h | 20 h | 14 h | 16 h | 37 h | 10 h |
| **Applying volume size[2]** | 256x256x256 voxels | 367x376x200 voxels | 340x460x59 voxels | ~256x384x50 voxels | ~208x311x41 voxels | ~270x370x45 voxels | 340x510x70 voxels | 512x256x21 voxels |
| **Applying time cost** | 39 s | 31 s | 23 s | 7 s | 3 s | 7 s | 23 s | 11 s |

**Supplementary Table 1 (continued).**

| **Samples** | **Ptk2 cells** | **Zebrafish** | **Adult *C. elegans*[3]** | **Adult *C. elegans*** | **Live mouse cardiac tissue** | **Fixed mouse liver tissue** | **Cleared mouse embryo[3]** |
|---|---|---|---|---|---|---|---|
| **Figures/Videos** | Fig 2a-f, Supp Figs 10-13 | Fig 2g-m, & Supp Fig 14 | Supp Fig 20 | Supp Figs 38, 39 & Supp Videos 15-17 | Fig. 3e, f, Supp Fig 21 & Supp Video 6 | Supp Video 7 | Fig. 4, Supp Figs 23-25 & Supp Videos 8, 9 |
| **Label** | Actin | Membranes | Neurons, 5 color channels | Neurites | Mitochondria | Membranes | Vessels, neurons |
| **Microscopy** | AO Lattice Light sheet microscopy | AO Lattice Light sheet microscopy | iSIM | iSIM | Two-photon microscopy | Two-photon microscopy | Confocal microscopy |
| **Detection NA** | 1.0 | 1.0 | 1.15 | 1.15 | 1.0 | 1.0 | 0.5 |
| **Maximum depth of raw volumes** | 28 μm | ~140 μm | 32 μm | 32 μm | 200 μm | 72 μm | ~2370 μm |
| **Depth of shallow subvolumes** | ~28 μm | ~20 μm | ~15 μm | ~20 μm | 35 μm | ~30 μm | ~1000 μm |
| **Size of shallow subvolumes** | ~430x770x132 voxels x 50 subvolumes | ~256x512x101 voxels x 45 subvolumes | 1028x500x50 voxels X 45 subvolumes | 568x490x100 voxels X 24 subvolumes | 1024x1024x64 voxels x 24 subvolumes | ~1024x1024x50 voxels x 24 subvolumes | ~740x620x180 voxels x 26 subvolumes (From 4 embryos) |
| **Training data size[1] (32 bit/voxel)** | 66 GB | 23 GB | 54 GB | 27 GB | 120 GB | 95 GB | 99 GB |
| **Training time cost** | 27 h | 15 h | 17 h | 18 h | 44 h | 39 h | 59 h |
| **Applying volume size[2]** | 1230x1500x370 voxels | 256x512x101 voxels | 1512x580x167 voxels | 1512x996x122 voxels | 1024x1024x304 voxels | 1024x1024x140 voxels | 1440x1740x433 voxels |
| **Applying time cost** | 75 s | 20 s | 2 min | 2.6 min | 5.7 min | 2.5 min | 15 min |

All samples and datasets used for training the DeAbe models in this paper. The time costs are reported based on tests on a Windows 10 workstation (CPU: Intel Xeon, Platinum 8369B, two processors; RAM: 256 GB; GPU: NVIDIA GeForce RTX 3090 with 24 GB memory). With additional footnotes:

1: The training data consists of paired GT images and synthetically aberrated images.

2: Representative single volume for testing the applying time cost.

3: For multicolor images, data size and time cost are reported for single color channel.

**Supplementary Note 1**, Synthetic generation of aberrated data from fluorescence images

***Basic concept***

The key insight of our approach is that fluorescence images acquired on the 'near side' of three-dimensional volumes are often close to diffraction limited. Such images thus represent ground truth data that can be used to train neural networks to reverse the effect of synthetically introduced aberrations. For this purpose, we intentionally synthetically aberrate the images acquired by fluorescence microscopes with a 'forward model'.

We assume a scalar imaging model with pupil function $H(r,\theta)$ given by

$$H(r,\theta) = |P(r,\theta)|e^{i\phi(r,\theta)} \tag{1}$$

$$r = \sqrt{u^2 + v^2}, \tag{2}$$

where $u, v$ are pupil plane coordinates, $r$ and $\theta$ are the radial and angular coordinates in the pupil plane, $\phi(r,\theta)$ is the phase aberration introduced either in the data or synthetically, and $P(r,\theta)$ is a binary pupil mask that spans the set of spatial frequencies transmitted by the imaging system:

$$P(r,\theta) = \begin{cases} 1,\ for\ r \leq \dfrac{NA}{\lambda} \\ 0, otherwise\ , \end{cases} \tag{3}$$

where $NA$ is the objective numerical aperture and $\lambda$ is the imaging wavelength.

The phase aberration, or wavefront distortion, can be conveniently expressed using Zernike basis functions $\phi_m(r,\theta)$ and associated coefficients $c_m$ :

$$\phi(r,\theta) = \sum_{m=0}^{M} c_m \phi_m(r,\theta)\,. \tag{4}$$

The corresponding Root Mean Square (RMS) wavefront distortion is calculated as

$$RMS = \sqrt{\frac{1}{\int P(r,\theta)drd\theta}\int \left[\phi(r,\theta) - \overline{\phi(r,\theta)}\right]^2 drd\theta} = \sqrt{\sum_{m=3}^{M} {c_m}^2}\,, \tag{5}$$

where $\overline{\phi(r,\theta)}$ is the mean wavefront, and the first three terms representing piston (m=0), tip (m=1) and tilt (m=2) are ignored.

The three-dimensional coherent point-spread function associated with this pupil function is given by[1]

$$h(x,y,z) = \mathcal{F}^{-1}\{H(r,\theta)e^{iz\gamma(r)}\}, \tag{6}$$

$$\gamma(r) = \frac{2\pi n}{\lambda}\sqrt{1 - \left(\frac{\lambda r}{n}\right)^2}\,, \tag{7}$$

where $x, y$ denote lateral coordinates, $z$ the axial coordinate and $n$ the refractive index. The incoherent PSF (appropriate for widefield fluorescence microscopy) is then given by

$$s(x,y,z) = |h(x,y,z)|^2. \tag{8}$$

Given an object $f(x, y, z)$ we can then describe the forward imaging model as:

$$d(x, y, z) = s(x, y, z) \otimes f(x, y, z), \tag{9}$$

where $\otimes$ is the convolution operation.

Omitting the $(x, y, z)$ coordinates for simplicity, the forward imaging model

$$d = s \otimes f \tag{10}$$

can be Fourier transformed ($\mathcal{F}(\square)$) to obtain

$$\mathcal{F}(d) = \mathcal{F}(s)\mathcal{F}(f), \tag{11}$$

where the Fourier transform of the PSF, $\mathcal{F}(s)$ is also known as the optical transfer function (OTF). If $\phi(r)$ = 0, i.e., the wavefront is 'flat' or unaberrated, we obtain the diffraction-limited image $d_0$ from the ideal PSF $s_0$ and object $f$ as

$$\mathcal{F}(d_0) = \mathcal{F}(s_0)\mathcal{F}(f). \tag{12}$$

Eliminating the object term using the last two equations,

$$\mathcal{F}(d) = \mathcal{F}(d_0)\left[\frac{\mathcal{F}(s)}{\mathcal{F}(s_0)}\right]. \tag{13}$$

To prevent division by zero, we modify the denominator by adding a small value $\alpha$ ($\alpha = 0.01$ for all datasets in this paper). Inverse Fourier transforming and optionally adding a noise term we obtain finally

$$d = \mathcal{F}^{-1}\left\{\mathcal{F}(d_0)\left[\frac{\mathcal{F}(s)}{\mathcal{F}(s_0) + \alpha}\right]\right\} + noise. \tag{14}$$

This equation provides a prescription for deriving an aberrated image $d$ from a diffraction-limited image $d_0$ and the ratio of aberrated OTF $\mathcal{F}(s)$ to diffraction limited OTF $\mathcal{F}(s_0)$. The degree of aberration can be tuned by adjusting the Zernike functions $\phi_m(r)$ and magnitude of their associated coefficients $c_m$.

In practice, we use images from the near side of fluorescence microscopy volumes as $d_0$ and tune $c_m$ until we obtain $d$ that resemble aberrated images that occur on the 'far side' of the image stack (see **Supplementary Table 1** for relevant parameters used in this paper).

***Extension to different microscopes***

To adopt this concept for other microscopes, we modify the PSF in equation (8), extending the wide-field PSF as needed to more accurately model light-sheet microscopy, confocal microscopy, instant SIM, and two-photon microscopy. Correspondingly, we re-write the equation (8) by adding a subscript "WF" to explicitly indicate it as using the wide-field PSF:

$$s_{WF} = s_{WF}(x, y, z) = |h(x, y, z)|^2. \tag{15}$$

We leave $s(x, y, z)$ as the generalized system PSF, which is constructed by considering the excitation PSF and emission PSF:

$$s = PSF_{sys} = PSF_{exc} \times PSF_{em}. \tag{16}$$

By substituting equation (16) into equation (10), we obtain the forward model of each microscope.

*1) Light sheet microscopy*

In light sheet microscopy, the emission PSF is equivalent to the wide-field PSF and the excitation PSF is often (as in these experiments) modeled as a virtual sheet constructed by scanning a low numerical aperture Gaussian beam across the field of view. We model the excitation sheet as uniform in the lateral directions but Gaussian in the axial direction (lateral and axial are defined from the perspective of the detection objective):

$$PSF_{exc} = \frac{1}{\sigma\sqrt{2\pi}} e^{-\frac{z^2}{2\sigma^2}}. \quad (17)$$

Then the final system PSF is

$$s_{LS} = PSF_{exc} \times PSF_{em} = \frac{1}{\sigma\sqrt{2\pi}} e^{-\frac{z^2}{2\sigma^2}} \times s_{WF} = A s_{WF} e^{-\frac{z^2}{2\sigma^2}}, \quad (18)$$

where $A$ is a constant which is discarded as we normalize the system PSF by integrating its intensity to 1. In practice, by measuring the thickness of the light sheet, i.e., the full width at half maximum (FWHM) in the axial direction at the beam waist, $\sigma$ can be estimated based on the assumption that the beam is Gaussian as:

$$\sigma = \frac{FWHM_{LS}}{2\sqrt{2ln2}} = \frac{FWHM_{LS}}{2.3548}. \quad (19)$$

*2) Confocal microscopy*

We construct the confocal PSF as:

$$s_{confocal} = PSF_{exc} \times PSF_{em} = s_{WF(\lambda_1)} \times \left(s_{WF(\lambda_2)} \otimes s_{Pinhole}\right), \quad (20)$$

where $s_{WF(\lambda 1)}$ and $s_{WF(\lambda 2)}$ are wide-field PSFs with excitation wavelength $\lambda_1$ and emission wavelength $\lambda_2$, respectively and $\otimes$ is the convolution function. $s_{Pinhole}$ models the physical pinhole in a confocal system as a binary circular mask at the z=0 plane:

$$s_{Pinhole}(x, y, z) = \begin{cases} 1, \ for\ z = 0\ and\ \sqrt{x^2 + y^2} \le Pinhole\ size \\ 0, \ \ otherwise. \end{cases} . \quad (21)$$

In practice, we implement the convolution operation in equation (20) in the Fourier domain, giving:

$$s_{confocal} = s_{WF(\lambda_1)} \times \mathcal{F}^{-1}\left\{\mathcal{F}\left(s_{WF(\lambda_2)}\right) \times \mathcal{F}\left(s_{Pinhole}\right)\right\}. \quad (22)$$

Note that we do not model the multifocal and pinhole lattice in spinning-disk confocal microscopy, so the model used for this form of microscopy is the same as what is used in point-scanning confocal microscopy.

*3) Instant SIM*

To simplify calculations, we neglected to model the multifocal illumination and pinhole lattice, instead modeling the PSF as a confocal PSF with infinitely small pinhole (a Dirac Delta function). In this case, the final super-resolution PSF is approximated by:

$$s_{iSIM} = \left(s_{WF(\lambda_1)} \otimes s_{Pinhole}\right) \times s_{WF(\lambda_2)} = s_{WF(\lambda_1)} \times s_{WF(\lambda_2)}. \quad (23)$$

Again, $s_{WF(\lambda 1)}$ and $s_{WF(\lambda 2)}$ are the wide-field PSFs with excitation wavelength $\lambda_1$ and emission wavelength $\lambda_2$, respectively.

*4) Two-photon microscopy*

For two-photon microscopy, the excitation PSF is the square of the wide-field PSF with the excitation wavelength. The emission PSF is treated as uniform as there is typically no confinement or modulation on the emission side, and this constant can be discarded as the final system PSF is normalized as described above. Therefore, we have:

$$s_{2P} = PSF_{exc} \times PSF_{em} = PSF_{exc} = {s_{WF(\lambda_1)}}^2. \quad (24)$$

For most datasets employing single photon microscopy, we used 488 nm excitation and 532 nm excitation; for two-photon microscopy data, we used 960 nm wavelength excitation.


1 Hanser, B. M., Gustafsson, M. G. L., Agard, D. A. & Sedat, J. W. Phase retrieval for high-numerical-aperture optical systems. *Optics Letters* **28**, 801-803 (2003).
